# Hypoxia Promotes Early Tumorigenesis through Dynamical-Mechanical Balance and Genetic Instability

**DOI:** 10.1101/2024.12.31.630973

**Authors:** Zhengduo Wang, Li Tian, Bo Li

## Abstract

Understanding early tumor formation provides insights for cancer prevention. However, in vivo studies focus on later stages, leaving initial tumorigenesis poorly understood. Hypoxia is a key microenvironmental factor in cancer, yet its impact on single-cell tumor initiation and dynamics remains unclear. To address this, we developed an integrated live-imaging platform for hypoxic conditions, enabling first direct observation of 3D tumor spheroid development from a single cell. We show hypoxia accelerates tumorigenesis. Beyond growth, directional movement promotes spheroid fusion. This is driven by optimized actin polymerization and cell-cell adhesion. Hypoxia induces specialized, fusion-competent cells with outstretched F-actin and tiny nuclei that bridge spheroids. Concurrently, hypoxia triggers genetic instability, evidenced by tiny nuclei and aneuploidy. The interplay between dynamic and genetic instability optimizes spheroid mechanical properties, creating a positive feedback loop fuels proliferation. Our findings offer insight into hypoxia’s role in early tumorigenesis and highlight potential of targeting dynamic-genetic balance for prevention.

**Statement of Significance:** Hypoxia accelerates early tumorigenesis by promoting directional 3D tumor spheroid fusion via optimized actin dynamics and mechanical balance, while inducing micronuclei formation and aneuploidy-linked genetic instability, forming a positive feedback loop that boosts proliferation and malignant progression.

## Introduction

Cancer becomes life-threatening after multiple stages of transformation. Hallmarks of advanced stages, such as metastasis, typically indicate a poorer prognosis (1). Thus, characterizing tumor features earlier is critical for effective detection and prevention (2,3). The prevailing model of early tumorigenesis posits that mutations confer a survival advantage, enabling transformed cells to proliferate and form primary tumors (4). However, this model remains indirectly supported. In vivo studies in animal models or patient samples rely predominantly on retrospective analyses, leaving early tumorigenesis poorly understood. Live cell imaging platforms enable direct observation of cellular processes within precisely controlled environments (5,6). Recent advances in 3D culture systems now enable ex vivo study of more complex processes (7,8), offering new avenues to investigate early tumorigenesis.

Hypoxia is a key microenvironmental factor that promotes cancer initiation and progression (9–11). While classical hypoxia signaling pathways influence diverse cellular processes (12–17), their role in early tumorigenesis remains unclear, largely due to a lack of direct evidence. Molecular analyses alone cannot capture the dynamics and spatial changes—like cell movement and fusion—that are critical in development (18) and metastasis (19). Furthermore, tumor cells interact with multiple environmental factors. Although mechanical forces influence tumorigenesis by modulating mechanical properties and adhesion (20–22), their interplay with hypoxia is poorly understood (23).

We developed an integrated platform to investigate early tumor formation, addressing a critical gap in understanding tumorigenesis. Hypoxia dramatically accelerates this process (Fig. 1). Contrary to molecular-centric models, we demonstrate that directional spheroid dynamics and fusion drive early tumorigenesis (Fig. 2). This fusion is facilitated by optimized mechanical-dynamical-molecular couplings and structurally supported by outreaching F-actin with underlying YAP signaling (Fig. 3). A concurrent emergence of genetic instability is marked by a new population of micronuclei cells, aneuploidy, and increased mitochondria number (Fig. 4). Post-fusion, a mechanics-genetic instability coupling promotes proliferation via p53 pathway modulation, further enhancing hypoxic growth (Fig. 5). These findings redefine hypoxia’s role in early tumor formation and challenge classical models focused solely on molecular cell-cycle pathways.

**Fig. 1.**
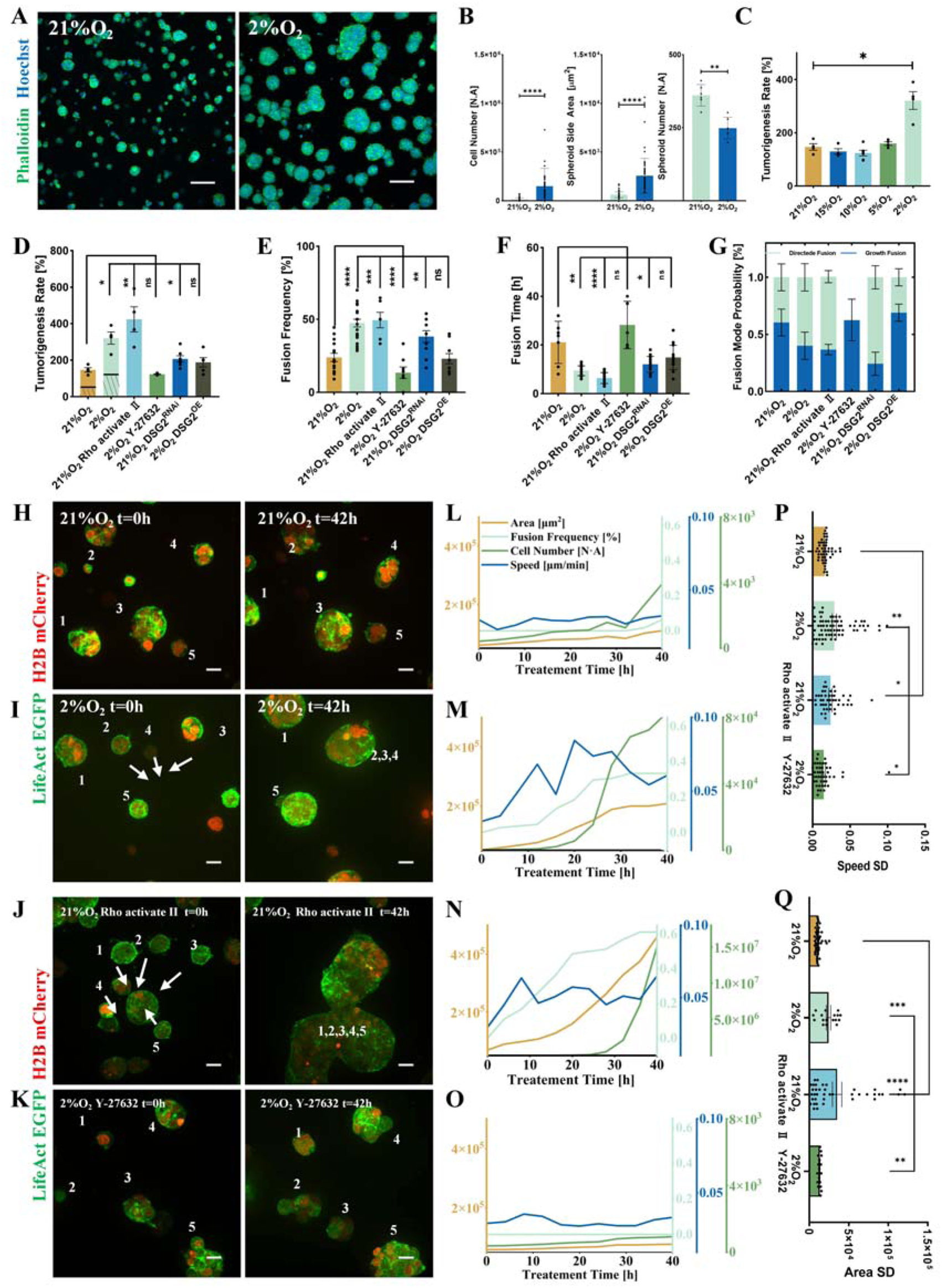
Hypoxia promotes early tumorigenesis by directional fusion. (**A**), Confocal images of HEK293T cells cultured in 3D Matrigel under 21% oxygen (left) and 2% oxygen (right) for 48 hours. The images are the typical reflection of more than three independent experiments. The green and blue channels are Phalloidin and Hoechst, respectively. The scale bar is 100 µm. (**B**), Cell number, maximum cross-sectional area of tumor spheroids, and total number of spheroids after 48 hours of 3D culture. (**C**), Tumorigenesis rate of HEK293T cells under different oxygen concentrations. (**D**), Tumorigenesis rate of HEK293T cells under different conditions of normoxia, hypoxia, and PHAi treatments. The shaded area in the first two columns represents the parts of tumorigenesis contributed by growth. (**E**), Fusion frequency of tumor spheroids under different culture conditions. (**F**), Time required for the directional fusion under different culture conditions of normoxia, hypoxia, PHAi, and RNAi treatments. (**G**), Proportion of directional and growth modes in the fusion under different culture conditions of normoxia, hypoxia, PHAi, and RNAi treatments. (**H-K**), Representative confocal images of more than three independent experiments that reflect the tumor spheroids’ fusion under different culture conditions. The left panels are images taken at 0 hours, and the right panels are taken at 42 hours after the treatment. Compared with the normal 21% oxygen control (**H**), directional fusion is promoted under 2% hypoxia conditions (**I**) or Rho activator II treatment (**J**). The directional fusion under hypoxia disappears after Y-27632 treatment (**K**). Green and red channels are LifeAct and H2B, respectively. The scale bar is 20 µm. (**L-O**), Time evolution of maximum cross-sectional area, fusion frequency, cell number, and speed of the spheroids under conditions of 21% Oxygen (**L**), 2% Oxygen (**M**), 21% Oxygen with Rho Activator II (**N**), and 2% Oxygen with Y-27632 (**O**). (**P, Q**), Standard deviation of the spheroid speed (**P**) and maximum cross-sectional area (**Q**) across different culture conditions. For all statistical plots, an unpaired Student’s t-test (n > 3 experiments, data are mean ± SEM) was conducted for all data with **** P < 0.0001, *** P < 0.001, ** P < 0.1, * P <0.5, ns P > 0.5. The error bars are the standard error of the mean of the data.

**Fig. 2.**
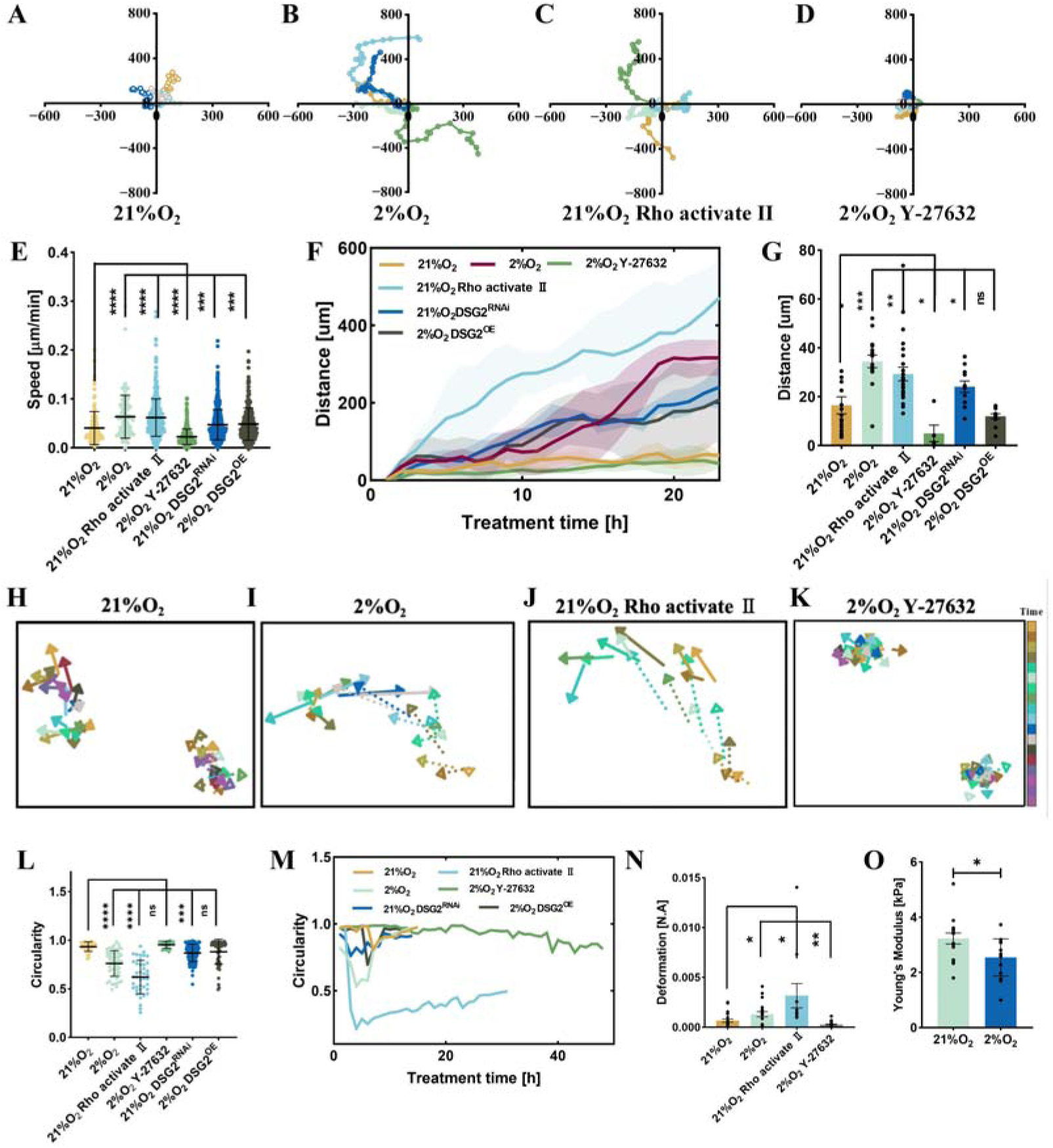
Dynamics of the tumor spheroids during the directional fusion. (**A-D**), Typical trajectories of tumor spheroids in 24 hours under different culture conditions of normal 21% oxygen (**A**), 2% hypoxia (**B**), Rho-activate II treatment under normoxia (**C**), and Y-27632 treatment under hypoxia (**D**). The data is a typical reflection of three independent experiments. The colors represent different tumor spheroids. (**E**), Speed in 3-dimension of tumor spheroids under different culture conditions of normoxia, hypoxia, PHAi, and RNAi treatment. (**F**), The distance between the non-fused tumor spheres and their initial positions over time under various conditions of normoxia, hypoxia, PHAi, and RNAi treatment. (**G**), The distance between two tumor spheroids before directional fusion occurs under various conditions of normoxia, hypoxia, PHAi, and RNAi treatment. (**H-K**), Typical trajectories of two fusing tumor spheroids under various conditions of 21% normal oxygen (**H**), 2% hypoxia (**I**), normoxia treated with Rho-activate II (**J**), and hypoxia treated with Y-27632 (**K**). The data is a typical reflection of three independent experiments. The dashed and solid vectors represent the trajectory of the two spheroids each. The direction and length of the vectors indicate the movement direction and speed of the spheroids at each time point, respectively. The color represents the time. (**L**), Circularity of the tumor spheroid during directional fusion under different culture conditions. (**M**), Time evolution of the tumor spheroid circularity during the directional fusion under various culture conditions of normoxia, hypoxia, PHAi, and RNAi treatments. (**N**), Degree of deformation of the tumor spheroid after laser ablation under different culture conditions of 21% normal oxygen, 2% hypoxia, normoxia treated with Rho-activate II, and hypoxia treated with Y-27632. The wavelength of the laser is 405 nm, and the power of ablation is 50 mW. (**O**), Cellular Young’s modulus under normoxic or hypoxic culture conditions. Further promotion of actin polymerization or the inhibition of intercellular interactions under hypoxic conditions suppresses early tumorigenesis. For all statistical plots, an unpaired Student’s t-test (n > 3 experiments, data are mean ± SEM) was conducted for all data with **** P < 0.0001, *** P < 0.001, ** P < 0.1, * P <0.5, ns P > 0.5. The error bars are the standard error of the mean of the data.

**Fig. 3.**
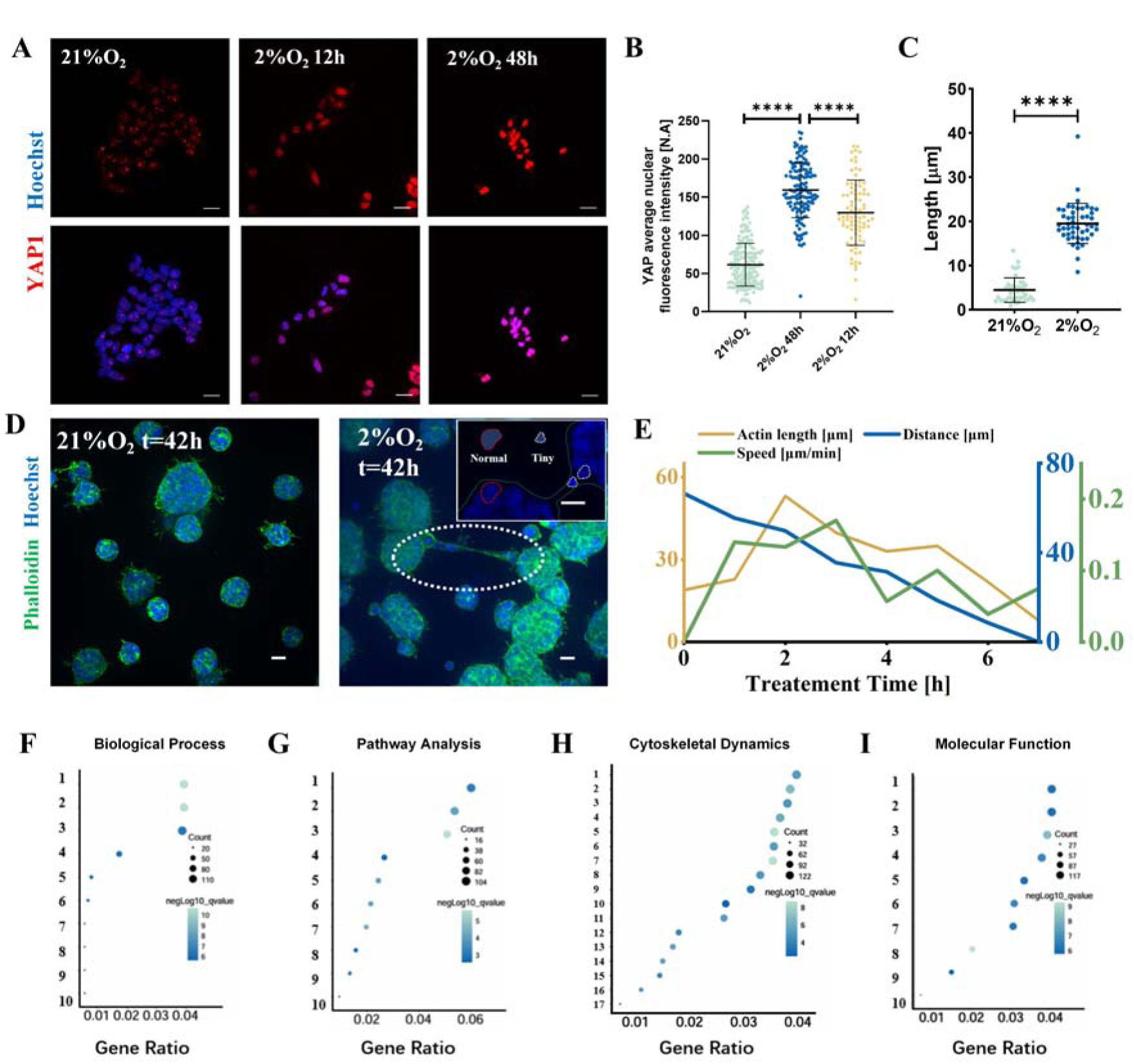
Outreaching F-acting guides the directional fusion. (**A**), Representative confocal images of YAP1 immunofluorescence in 3D cultured cancer cell spheroids under normoxic conditions for 48 h (left), hypoxic conditions for 12 h (middle), and hypoxic conditions for 48 h (right). YAP1 immunostaining is shown in red, and nuclei were counterstained with Hoechst (blue). The scale bar is 20 µm. (**B**), Quantification of mean YAP1 fluorescence intensity in the nucleus of 3D cultured cancer cell spheroids under normoxic conditions for 48 h, hypoxic conditions for 12 h, and hypoxic conditions for 48 h. (**C**), The length of out-reaching F-actin from the spheroid surface in normoxic and hypoxic conditions. (**D**), Typical immunostaining images of the tumor spheroids under normoxia (Left) and hypoxic (Right) conditions. The green channel is Phalloidin, and the blue channel is Hoechst. The dashed white circles indicate the outreaching F-acting structure that bridges two fusing tumor spheroids. The inset on the right is an enlarged view of the outstretched F-actin channel between two spheroids. The dashed green line depicts the envelope of the spheroids and the channel. The dashed white circles demonstrate the tiny nuclei, and the dashed red circles demonstrate the normal nuclei. The scale bar is 20 µm. (**E**), Time evolution of the F-actin bridge length, the distance between the spheroid and the spheroid’s speed during directional fusion. (**F-I**), Differential analysis of 3-dimensional HEK293T transcriptomic data under hypoxia and normoxia conditions. (**F**), KEGG biological process enrichment analysis of differentially expressed genes. The bubble plot shows the top ten enriched biological processes. The y-axis is described as follows. 1: extracellular structure organization, 2: extracellular matrix organization, 3: gland development, 4: nucleoside diphosphate metabolic process, 5: NAD metabolic process, 6: glucose catabolic process, 7: glucose catabolic process to pyruvate, 8: canonical glycolysis, 9: NADH regeneration, 10: glycolytic process through glucose−6−phosphate. (**G**), KEGG pathway enrichment analysis of differentially expressed genes. The bubble plot shows the top ten KEGG pathways. The y-axis is described as follows. 1: PI3K−Akt signaling pathway, 2: MAPK signaling pathway, 3: Calcium signaling pathway, 4: Oxidative phosphorylation, 5: Protein digestion and absorption, 6: ECM−receptor interaction, 7: Biosynthesis of amino acids, 8: Glycolysis / Gluconeogenesis, 9: Arginine and proline metabolism, 10: Pentose phosphate pathway. (**H**), KEGG enrichment analysis of differentially expressed genes related to cytoskeletal dynamics. The y-axis is described as follows. 1: cell growth, 2: cell−cell junction, 3: regulation of GTPase activity, 4: actin binding, 5: cell−substrate junction, 6: actin filament organization, 7: focal adhesion, 8: positive regulation of GTPase, 9: activity Ras GTPase binding, 10: tubulin binding,11: embryonic organ morphogenesis, 12: actin filament binding, 13: site of polarized growth, 14: adherens junction, 15: Rho GTPase binding, 16: actin filament, 17: Rac GTPase binding. (**I**), KEGG molecular function enrichment analysis of differentially expressed genes. The y-axis is described as follows. 1: channel activity, 2: passive transmembrane transporter activity, 3: metal ion transmembrane transporter activity, 4: ion channel activity, 5: monovalent inorganic cation transmembrane transporter activity, 6: cation channel activity, 7: gated channel activity, 8: extracellular matrix structural constituent, 9: growth factor binding, 10: extracellular matrix structural constituent conferring tensile strength. For all the KEGG plots, the size of each bubble represents the number of genes involved, while the color indicates the significance level (p-value). The mRNA samples are extracted from cells in 3-dimensional culture after 24 hours of 2% (hypoxia) or 20% (normoxia) oxygen treatment. All data are from three independent experiments. For all statistical plots, an unpaired Student’s t-test (n > 3 experiments, data are mean ± SEM) was conducted for all data with **** P < 0.0001, *** P < 0.001, ** P < 0.1, * P <0.5, ns P > 0.5. The error bars are the standard error of the mean of the data.

**Fig. 4.**
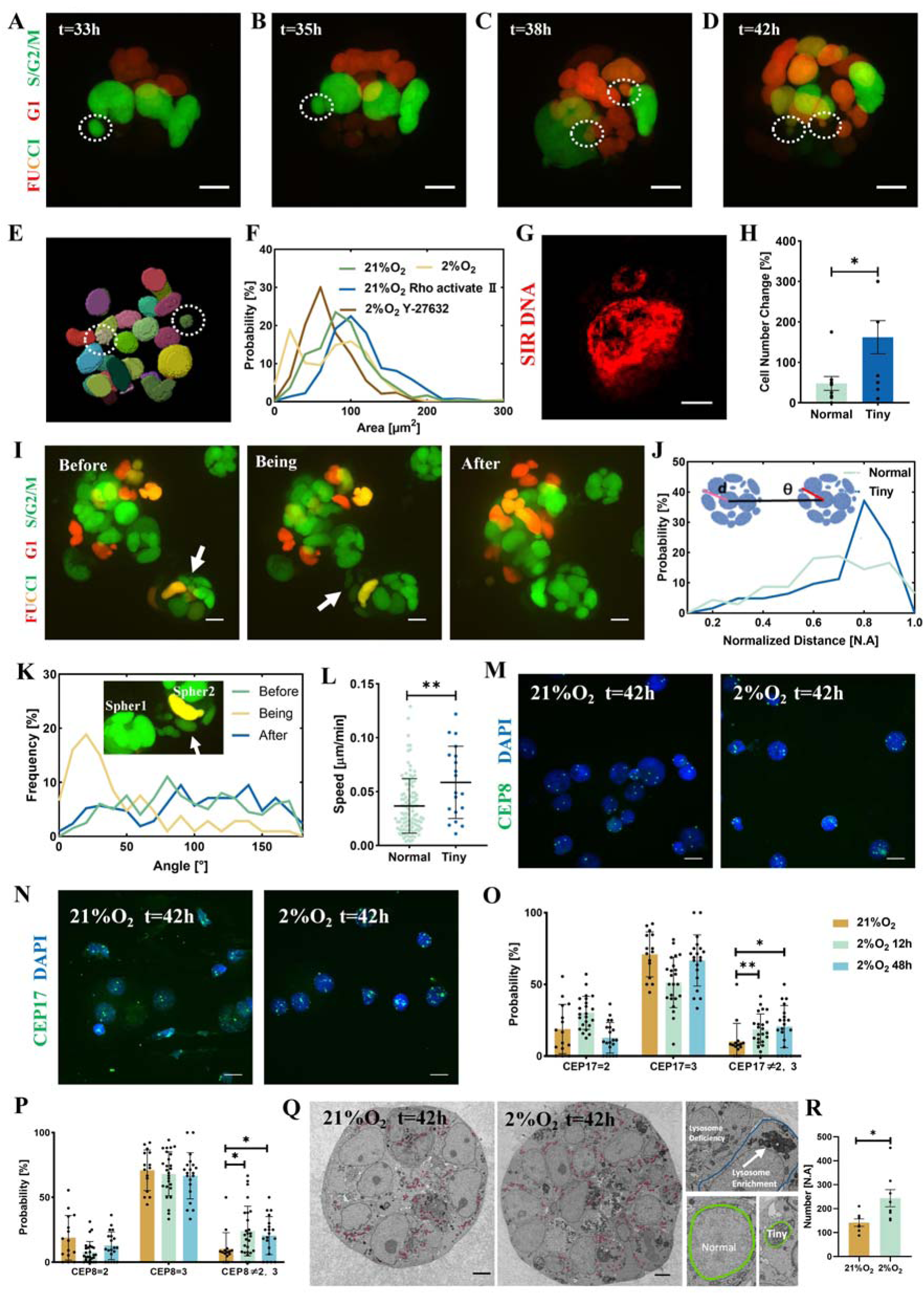
Micronuclei, aneuploidy, and mitochondria hyperplasia. (**A-D**), Typical confocal images from more than three independent experiments of tumor spheroid composed of HEK293T-FUCCI cells in 3D culture for 42 hours under 2% oxygen conditions. The cells are labeled with FUCCI, in which the color indicates different stages of the cell cycle. Red marks hCdt1, indicating cells are in the G1 phase, and green marks hGem, indicating cells are in the S/G2/M phases. The white dotted circle indicates the nucleus of the tiny cells. The scale bar is 10 µm. (**E**), 3D segmentation results of tumor spheroids via Cellpose; white circles represent the segmented small nuclei. (**F**), Probability distribution of nucleus sizes under different 3D culture conditions of normoxia, hypoxia, normoxia treated with Rho-activate II, and hypoxia treated with Y-27632. The dashed line indicates a bimodal distribution in the probability of nuclear size under hypoxic conditions, suggesting the presence of nuclei with different sizes. (**G**), Representative stimulated emission depletion (STED) images of cells stained with SIR-DNA to label nuclei and tiny nuclei. The scale bar is 2.5 µm. (**H**), Relative changes in the number of tiny and normal nuclei over the same period of 42 hours. (**I**), Confocal images of HEK293T-FUCCI tumor spheroids during the directional fusion process. The tiny cells appear at the proximal gap between the two spheroids during the fusion, but are not observed before and after. The white arrow in (**I**) indicates the nucleus of the tiny cells. The green and red channels are hCdt1 and hGem, respectively. (**J**), Probability distribution of the distance (“d” in the inset) between the tiny and normal cells at the center of tumor spheroids during directional fusion. A distance of 1 indicates positioning at the spheroid’s edge. The inset of (**J**) schematically demonstrates the relative positions between the tiny cells and the spheroids. (**K**), Probability distribution of the angle (“θ”) in the inset of (**K**) between the tiny cell nucleus–spheroid center axis and the spheroid center-spheroid center axis during the directional fusion. A smaller θ value means a closer location to the proximal ends of the axis that connects the mass centers of the two spheroids. The inset of (**K**) is the enlarged part of the gap region between two fusing spheroids. The white arrow indicates the nucleus of the tiny cells. (**L**), Speed of tiny and large nuclei in cells. (**M, N**), Representative confocal images showing DNA FISH labeling of the centromere of chromosome 8 (**M**) and chromosome 17 (**N**) in 3D-cultured cancer cell spheroids under normoxia (48 h, left) and hypoxia (48 h, right). The centromeres are visualized with green fluorescent labeling. Nuclei were counterstained with Hoechst (blue). The scale bar is 20 µm. (**O, P**), Statistical analysis of the probabilities of single cells having exactly 2, exactly 3, or a number not equal to 2 or 3, for chromosome 8 (**P**) and chromosome 17 (**O**), after treatment under normoxia for 48 h, hypoxia for 12 h, or hypoxia for 48 h. (**Q**), Representative scanning electron microscopy (SEM) images of cancer spheroids under three-dimensional culture conditions, cultured for 42 h under 21% (left) and 2% (right) oxygen. Mitochondria are indicated by red circles. Right upper inset: A magnified view of two adjacent cells within the tumor spheroid, showing a notable difference in the number of lysosomes. The white arrows indicate regions of lysosome accumulation. Right lower inset: A schematic diagram illustrating the maximum cross□section of a micronucleus and a normal nucleus from the same tumor spheroid at the same scale. The scale bar is 5 µm. (**R**), Quantitative analysis of the mitochondrial number per cell in cancer spheroids cultured under 21% and 2% oxygen for 42 h. For all statistical plots, an unpaired Student’s t-test (n > 3 experiments, data are mean ± SEM) was conducted for all data with **** P < 0.0001, *** P < 0.001, ** P < 0.1, * P <0.5, ns P > 0.5. The error bars are the standard error of the mean of the data.

**Fig. 5.**
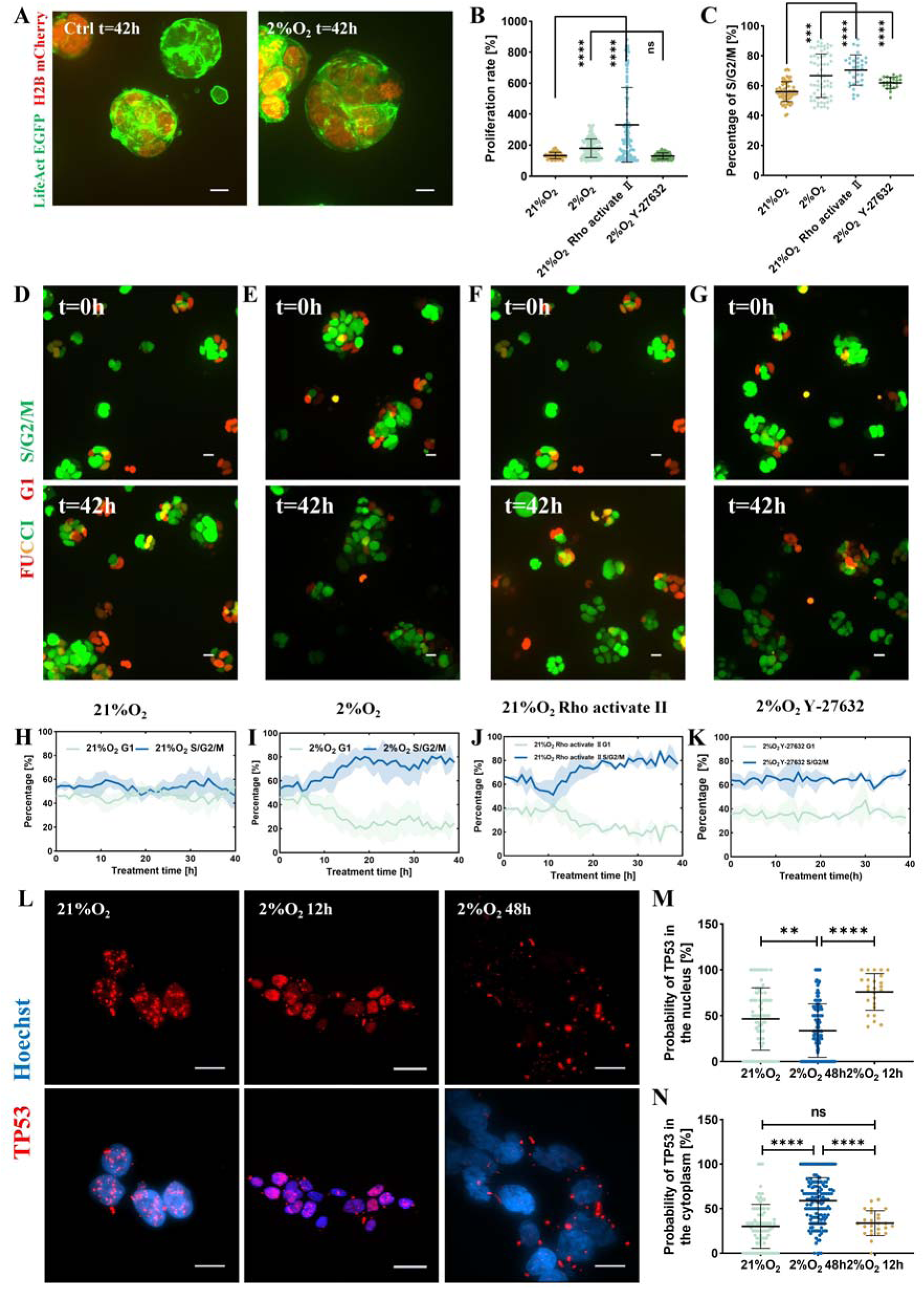
Hypoxia promotes spheroid growth in 3D by modulating the cell cycle. (**A**), Representative Confocal images of tumor spheroid under 21% oxygen (left) and 2% oxygen (right) conditions after 42 hours of 3D culture. The images are the typical reflection of more than three independent experiments. The green and red channels are LifeAct and H2B, respectively. The scale bar is 10 µm. (**B**), Proliferation rate of HEK293T cells under different 3D culture conditions of normoxia, hypoxia, and PHAi experiments. (**C**), Proportion of HEK293T cells in the S/G2/M phases under different 3D culture conditions of normoxia, hypoxia, and PHAi experiments. (**D-G**), Typical confocal images of tumor spheroids composed of HEK293T-FUCCI cell line under various culture conditions of normoxia control (**D**), hypoxia (**E**), normoxia with Rho activate II treatment (**F**), and hypoxia with Y-27632 treatment (**G**). The upper column is images taken at 0 hours, and the lower column is images taken at 42 hours. The green and red channels are hCdt1 and hGem, respectively. The scale bar is 20 µm. (**H-K**), Proportion of cells in G1 phase or S/G2/M phases as a function of time normoxia (**H**), hypoxia (**I**), normoxia with Rho activate II treatment (**J**), and hypoxia with Y-27632 treatment (**K**). (**L**), Representative structured illumination microscopy (SIM) and confocal images of TP53 immunofluorescence in 3D-cultured cancer cell spheroids following 48 h of normoxia (left), 12 h of hypoxia (middle), or 48 h of hypoxia (right). Red fluorescence indicates TP53, and blue fluorescence corresponds to Hoechst-stained nuclei. The scale bar is 10 µm. (**M, N**), Statistical analysis showing the probability of TP53 localizing to the nucleus (**M**) or to the cytoplasm (**N**) in cells subjected to 48 h of normoxia, 12 h of hypoxia, or 48 h of hypoxia. For all statistical plots, an unpaired Student’s t-test (n > 3 experiments, data are mean ± SEM) was conducted for all data with **** P < 0.0001, *** P < 0.001, ** P < 0.1, * P <0.5, ns P > 0.5. The error bars are the standard error of the mean of the data.

The research platform includes several components (Fig. S1). First, a human kidney-derived tumor cell line, genetically modified for protein visualization, was used to analyze underlying mechanisms (Fig. S2 and Tables S1, S2). The HEK293T tumor cell line (7,8,24–26) was seeded in a Matrigel suspension, where cells grew as 3D spheroids by occupying cavities larger than the spheroids (Fig. S3A and B). Notably, only in 3D culture did cancer and cancer stem cell markers express fully (Fig. S4A-C), rationalizing the system as a model for tumorigenesis (25,27). A spinning-disk confocal microscope, coupled with a three-gas incubator, enabled long-term live-cell imaging with minimal phototoxicity. By adjusting the nitrogen gas supply, tumorigenesis was observed under different oxygen conditions, from hypoxia (2%) to normoxia (21%). Finally, live-cell imaging data were analyzed using AI-assisted segmentation (Fig. S3C and D) alongside molecular cell biology data (Tables S3, S4) to elucidate the mechanism. Further experimental details are provided in the Methods section.

## Results

### Hypoxia promotes tumorigenesis through directional fusion

After 42 hours of culture, tumor spheroids formed under 2% oxygen were significantly larger than those grown under normoxic conditions (Fig. 1A and Movie S1). Quantitative analysis shows a fourfold increase in tumor size and total cell number under hypoxia, with a moderate decrease in spheroid number by 31% (Fig. 1B). Under different oxygen concentration treatments, only the 2% oxygen concentration cultivation condition significantly upregulated the formed tumor spheroid area (Fig. 1C and Fig. S5A-C). For suspended NCI-H69 cells cultured under 2% oxygen, we further observed a marked elevation in the number of tumor spheroids under hypoxic relative to normoxic conditions (Fig. S5D and E). These results clearly indicate that hypoxia accelerates early tumorigenesis. Two dynamic processes contribute to this effect: fusion between tumor spheroids and growth driven purely by cell proliferation. For both normoxia and hypoxia, fusion accounts for approximately 70% of tumorigenesis, significantly outweighing the ∼30% contribution from proliferation alone (Fig. 1D). Under hypoxia, fusion frequency doubles (Fig. 1E), and fusion event duration decreases by 55% (Fig. 1F), leading to increased tumor size in a shorter time (Movie S2).

Fusion events can be further classified into two modes (Fig. 1G). In the directional mode, tumor spheroids move freely within the Matrigel network until their outer surfaces make contact (Fig. 1I, Movie S3). We refer to this as “directional fusion”. In the growth mode, spheroid centers oscillate near their initial positions, and fusion results as a by-product of growth (Fig. 1H and Movie S4). Under hypoxic conditions, directional fusion predominates (Fig. 1G), indicating that hypoxia drives early tumorigenesis by increasing spheroid dynamics. Based on these results, we conclude that directional fusion events and associated spheroid dynamics are key drivers of enhanced tumorigenesis.

Tumor cells within spheroids exhibit greater survival than isolated cells in 3D cultures (28,29), fostered by a protective microenvironment of close cell-cell contacts. This enhanced viability promotes cell dynamics and proliferation, establishing a positive feedback loop that accelerates tumorigenesis. Time-resolved analysis supports this model (Fig. 1L-O). Under hypoxia, spheroid speed increases immediately, confirming dynamics as central to tumorigenesis. This rapidly lowers barriers to contact, elevating fusion frequency after a brief 6-hour lag. Proliferation rates rise ∼10 hours later, reflecting improved survival in larger spheroids, with a marked increase in overall spheroid size signifying robust tumorigenesis after ∼24 hours (Fig. 1M). This cascade is absent under normoxia, where speed, fusion, and proliferation remain stable (Fig. 1L). Thus, hypoxia initiates a cascade where enhanced dynamics drive fusion, which in turn promotes growth.

Accompanying the directional fusion process, notable heterogeneity in both the dynamics and morphology of spheroids emerges. Under hypoxic conditions, the standard deviations of both spheroid speed and size increase significantly compared to those observed in normoxia (Fig. 1P and Q). As two spheroids are undergoing directional fusion, their dynamics slow down, aligning with the sharp decrease in speed observed approximately 10 hours after hypoxia treatment (Fig. 1M). Since not all spheroids undergo fusion, those growing solely through proliferation naturally remain smaller than those formed via directional fusion, leading to a broader distribution of spheroid sizes under hypoxia (Fig. 1Q).

### Dynamical and mechanical balance facilitates spheroid fusion

Known its critical role in tumorigenesis, we examine the cellular and molecular mechanisms underlying the directional movements. Representative spheroid trajectories (Fig. 2A-D) show navigation ranges prior to fusion. Under hypoxia, spheroids become unpinned from their original positions, moving freely in 3D space, with speed increases by 57.4 % (Fig. 2E), allowing fusion to occur over greater distances (Fig. 2F and G). Remarkably, the motion of fusing spheroids shifts from a random walk to directional movement toward each other (Fig. 2H-K), efficiently enabling fusion and enhancing its frequency.

Hypoxic spheroids modulate their mechanics and morphology to facilitate directional fusion. They exhibit reduced circularity (Fig. 2L), greater deformation over time (Fig. 2M), and undergo more extensive structural reorganization after laser ablation of single cells (Fig. 2N and Movie S5). Nano-indentation confirms that individual cells within these spheroids are softer (Fig. 2O). Collectively, these mechanical adaptations enhance spheroid dynamics (Fig. 2E-G) and shorten fusion time (Fig. 1F). To define the molecular basis of these dynamical-mechanical couplings, we perturbed cell mechanics pharmacologically and genetically (Table S5). Enhancing actin polymerization phenocopies hypoxia, increasing tumor formation and fusion frequency (Fig. 1D, E, and Movie S6), while its inhibition suppresses them. This manipulation also recapitulates hypoxic effects on circularity, fusion time, and heterogeneity (Fig. 1F, P, Q, and Fig. 2L-N). Chronological analysis confirms that actin polymerization drives a dynamic sequence identical to hypoxia (Fig. 1N and O), establishing it as a key molecular regulator.

The functional role of actin polymerization in early tumorigenesis observed in the present experiments implies the activation of those signal pathways that control cell responses to mechanical cues under hypoxia (30), although at the time of promoted actin polymerization, no upregulation of Hypoxia-Inducible Factor 1 Alpha Subunit (HIF1α) yet occurs (Fig. S4I). Among those, Hippo signaling and its core regulator, Yes-associated protein (YAP), play pivotal roles by regulating actin cytoskeleton dynamics and cellular mechanics (31,32). Immunofluorescence staining reveals significant YAP1 nuclear accumulation under hypoxia (Fig. 3A and B). As the interaction between YAP and actin polymerization is mutual (33,34), YAP1 can, in turn, facilitate the fusion of tumor spheroids.

To define the structural basis of directional fusion, we analyzed subcellular dynamics. Immunostaining under hypoxia reveals that F-actin forms distinctive outreaching structures, specifically bridging adjacent fusing spheroids (Fig. 3D, Fig. S6A-H, and Movie S7). These extensions are polarized, localizing to the inter-spheroid gap, and can attain lengths comparable to a spheroid’s diameter (Fig. 3C), vastly extending their sensing range. Live-cell imaging confirms that F-actin protrusions dynamically elongate toward a target spheroid hours before fusion (Fig. S6E-H and Movie S8). Chronological analysis shows actin elongation coincides with accelerated spheroid movement and distance reduction, confirming their correlation (Fig. 3E). In addition, laser ablation of these F-actin extensions reduces hypoxic tumorigenesis (Fig. S7E-H), proving their essential role as the structural basis for remote sensing and directional fusion.

To further assess the roles of cell-cell adhesion and cell-ECM interaction, we performed RNA interference (RNAi) and Overexpression (OE) manipulations of Desmoglein-2 (DSG2) and integrin β-3 (ITGB3), both of which have been reported to be hub modulators in tumorigenesis (35,36). Here we briefly report that tumor spheroids reduce intercellular adhesion to facilitate greater deformation and enhance dynamics, while cell-ECM interaction plays a minor role in the hypoxic environment (Fig. S8). More detailed analysis of these experiments can be found in the Supplementary Information. In 2D cultures, however, cell-ECM interactions influence both tumorigenesis and directional fusion (Fig. S9), highlighting fundamental differences in dynamical-mechanical couplings between 2D and 3D culture systems.

The fusion and deformation of tumor spheroids represent complex adaptations beyond equilibrium physics. While greater deformation typically signifies reduced stiffness in equilibrium systems (37), we observe that actin polymerization—generally associated with increased stiffness (38)—paradoxically enhances deformation and dynamics here (Fig. 1D-G and Fig. 2E-K). Furthermore, neither additional actin polymerization nor DSG2 knockdown under hypoxia further enhances fusion (Fig. S8I-L), indicating an optimal concentration for each molecule to maintain spheroid function. Transcriptomic analyses reveal hypoxia-induced upregulation of gene clusters involved in metabolism, cancer signaling, cytoskeletal dynamics, and ECM remodeling (Fig. 3F-I). These findings show that cells finely optimize their biomechanical and biochemical properties in response to hypoxia.

### Genomic instability manifested as tiny cells and aneuploidy

Following hypoxia treatment, a distinct cell type with notably small nuclei emerges within the tumor spheroids (Fig. 4A-D, Fig. S6I-P and Movie S9). These “tiny” cells, with nuclear sizes averaging 23.9 % of that of typical nuclei, are virtually absent under normoxic conditions. The nuclei size distribution shifts to a bimodal pattern under hypoxia, contrasting with the continuous distribution observed in normoxia (Fig. 4F). Chromosome immunostaining verifies these tiny nuclei as ring-shaped cells (Fig. 4G) rather than organelle-like entities, and their number-increase rate is nearly quadruple that of normal cells (Fig. 4H). The emergence of distinct nuclear size distribution and number increased rate provides strong evidence that these tiny nuclei represent a novel cell subtype within the tumor spheroid. This adaptation to environmental stress suggests underlying genetic instability, a known hallmark of cancer (39).

Tiny cells are pivotal for directional fusion. They localize predominantly at the spheroid surface, clustering specifically at the interface toward a neighboring spheroid to bridge the gap (Fig. 4I-K). These cells colocalize with outreaching F-actin, which forms connecting channels (Inset of Fig. 3D). Their functional importance is confirmed by laser ablation, which reduces tumorigenesis (Fig. S7A-D). We propose that directional fusion arises from cytoskeletal-nuclear coordination: F-actin extends the sensing range, while motile tiny cells (Fig. 4L) act as “sentinels,” traversing these channels and facilitating mass transport. Their high motility enables rapid positional adjustment and influences overall spheroid dynamics (Fig. 1P and Q). Thus, the coupling between dynamical and genetic properties jointly determines the tumorigenesis rate.

At the chromosomal level, the genetic instability is manifested as aneuploidy. DNA fluorescence in situ hybridization (FISH) experiments show that under both normoxic and hypoxic conditions, more than 50% of the cells exhibited three copies of chromosomes 8 and 17 rather than the normal diploid number of two (Fig. 4M-P). This result is consistent with the real space observation (Fig. 4M-P) and sequencing analysis (Fig. S4A-C) that the 293T cells present cancer stem cell-like character under 3D-cultured conditions, namely, the 3D environment has already induced some degree of genetic instability to the tumor cells. Hypoxia treatment further increased the proportion of cells with chromosome 8 and 17 copy numbers other than two or three by about one fold, indicating elevated genomic instability at the chromosomal level. In particular, the genes that encode Tumor Protein 53 (TP53) and Junction Plakoglobin (JUP) lie on chromosome 17 (40,41), the abnormality of which is consistent with the loss of proliferation control by the functional impairment associated with localization changes of p53 structure; and regulation of cell-cell adhesion, respectively. Note that the increase in chromosome number has already been detectable as early as 12 hours after hypoxia treatment, proving it to be a causal effect, just as the directional fusion, for the enhanced tumorigenesis. The aneuploidy persisted throughout the whole time course of the experiments since its occurrence. Usually, the genetic instability in cancer is discussed at a much later stage after the primary tumor has been constructed (39). It is remarkable that it occurs so early, accompanying the process by which tumors are formed from a few transformed cells.

Genetic stability also emerges at the organelle level. Mitochondria and lysosomes carry genes that regulate the cell metabolism (42) and biochemical reactions (43), respectively, and thus act as the centers for bioactivities. Scanning electron microscopy revealed that the number of mitochondria per cell under hypoxic conditions is 30 % higher, compared with that of normoxia (Fig. 4Q and R). The elevated mitochondrial count could help provide additional energy to support directional movement and enhanced cell proliferation under hypoxia. Under hypoxia, cells in the spheroids divide into two subgroups regarding lysosomal content. Approximately 35% of cells contained abundant lysosomes, while lysosomes can hardly be observed in the remaining ones (Fig. 4Q, inset and Supplementary Information). The existence of tiny cells is further confirmed by the electron microscopy (Fig. 4Q, inset)

### Stimulated cell cycle enhances proliferation and spheroid growth

Following directional fusion, the rate of cell proliferation shows a marked increase, indicating a coupling between spheroid dynamics and cell proliferation (Fig. 1M). Although cell proliferation-driven growth of the tumor spheroid is less prominent than fusion, it still accounts for approximately 20 % of the overall tumorigenesis rate (Fig. 1D). Under hypoxic conditions, this contribution rises by 40 % (Fig. 1D), alongside a substantial increase in cell proliferation rate (Fig. 5A and B), underscoring the importance of proliferation-driven growth in tumor spheroid development.

To visualize cell cycle dynamics, we employed the FUCCI technique, enabling precise tracking of cell cycle phases (Supplementary Information, Table S2). Statistical analysis reveals that under hypoxia, the proportion of cells in the S/G2/M phases is approximately 10 % higher than under normoxia (Fig. 5C), suggesting a hypoxia-induced acceleration of cell proliferation. Over time, this effect amplifies, with nearly all cells entering the S/G2/M phases after 42 hours of hypoxia exposure (Fig. 5E, I, and Movie S10). Notably, the rate of increase in the S/G2/M cell proportion is non-linear. Initially, the proportion of cells in G1 and S/G2/M remains stable, but after a critical threshold, the S/G2/M phase proportion begins a rapid ascent (Fig. 5I), marking a proliferation rate jump confirmed by cell counts (Fig. 1M). This analysis supports the hypothesis that larger spheroids, which are better suited for cell survival in 3D culture, stimulate cell proliferation, contributing to overall spheroid growth.

Mechanistically, the stimulation of the cell cycle within the nucleus is linked to cytoskeletal dynamics, echoing previous studies at the molecular level (44). When actin polymerization is activated, the proportion of cells in S/G2/M phases increases, even in the absence of hypoxia (Fig. 5J). Conversely, inhibiting actin polymerization prevents this increase in S/G2/M phase cells under hypoxia (Fig. 5K). Tiny cells, which number increase at a rate four times that of normal cells, fail to form when actin polymerization is suppressed (Fig. 4F). Thus, in addition to hypoxia, specific cytoskeletal conditions are essential for the generation of tiny cells and the enhanced proliferation they drive. Therefore, the proliferation within tumor spheroids is modulated by cytoskeletal mechanics, linking external mechanical properties to nuclear behavior.

Immunostaining reveals the mechanism for stimulated cell cycling via p53 (Fig. 5L-N). As a known guardian of genome integrity (45), p53’s role in tumorigenesis is context-dependent (46,47). We find that under normoxia and early hypoxia, nuclear p53 acts to restrict proliferation in already-transformed HEK293T cells, which exhibit spheroid growth and aneuploidy even at baseline (Fig. 1E,2B, 2E, 2F, 4O and 4P). However, after 48 hours of hypoxia, prolonged stress induces p53 functional impairment associated with localization changes, disrupting its nuclear localization and function (Fig. 5L-N). This loss of proliferative control accelerates growth, as confirmed by FUCCI imaging. Thus, proliferation emerges from dynamic-genetic coupling: hypoxia-driven fusion and genetic instability increase cell numbers, with larger spheroids enhancing survival, while p53 impairment releases cell-cycle checkpoints, collectively propelling a malignant growth state.

## Discussion

In summary, this study provides experimental insights into how hypoxia accelerates early tumorigenesis (Fig. S10). By establishing a 3-dimensional culture, we are able to construct an ex vivo system in which tumor spheroids are formed from individual cancer stem cells. We find that the enhanced directional dynamics, whose underlying balance with the cytoskeletal mechanics and cell cycle kinetics are the major driving factors of tumorigenesis. The genetic instability fully involves early tumorigenesis by coupling with the dynamical processes. These findings not only challenge traditional linear models of early tumorigenesis that often reduce it to simple proliferation and growth but also offer potential prevention or therapeutic strategies aiming at the dynamo-mechano-physio-coupling, in addition to those targeting classical signal pathways (48). The scenario proposed by our experiment is valid for the primary tumor formation from individual cancer stem cells (49), as well as for the metastatic formation from the disseminated cells at the distal end (50).

Our findings also raise new research questions. Dynamics, a concept typically outside the focus of molecular cell biology, proves critical for understanding tumorigenesis in this context. However, the transcriptomic analysis fails to identify classical signaling pathways in existing databases (Fig. S11), although chemical hypoxia induced by CoCl2 replicates similar spheroid behaviors (Fig. S11A-E) – suggesting unknown molecular mechanisms underlying the observed dynamic processes. The relation between phenotypic heterogeneity and genetic instability needs to be further elucidated. Moreover, the consequence of the behaviors in early tumorigenesis, namely, their potential involvement in later stages of cancer development, such as metastasis, should be tested by an in vivo model. Last but not least, whether the model proposed by this work is general to tumorigenesis needs to be tested in other specific cancer cell lines.

## Methods

### Cell line maintenance and culture

HEK293T (ATCC) cells were grown in DMEM (Adamas life, C8013-500mL) supplemented with 10% heat-inactivated FBS (Gibco). NCI-H69 (ATCC) cells were grown in RPMI 1640 (Adamas life, C8016-500mL) supplemented with 10% heat-inactivated FBS (Gibco). Cells were maintained in an incubator at 37□°C and 5 % CO2 and passaged every 2 to 3□days. Before imaging, the medium was changed or Pharmaceutical interference (PHAi) treatments were applied according to experimental requirements, including DMSO (Adamas Life, 75927AA, 30 µM), Rho Activator II (Cytoskeleton, Cat. # CN03, 2 µg/mL), or Y-27632 (Yesen, 53006ES08, 30 µM). The prepared cell samples were then transferred to a live-cell workstation, where confocal imaging was performed. The mycoplasma testing was conducted weekly on the cell lines using the Mycoplasma Detection Kit (Solea, CA1080). We use Corning Matrigel (356234-5mL) to create a 3D culture environment that assists the tumor cells in forming spheroids (51,52). After thawing the Matrigel according to the manufacturer’s recommendations, mix it thoroughly with a pipette to ensure homogeneity. Dilute the Matrigel to a 50% concentration with a serum-free medium. Then, pour the diluted Matrgel on the confocal dish and incubate at 37°C for 15 minutes to allow solidification. Finally, seed the cells on the confocal dish with solidified Matrigel and add complete culture medium.

### Pharmaceutical interference experiment

Rho Activator II (CN03) from Cytoskeleton (53,54), is stored in a desiccated state at 4°C for no more than 6 months. For reconstitution, briefly centrifuge to collect the product at the bottom of the tube and resuspend each vial in 200 µL of sterile water, place on ice for 10 minutes prior to mixing, and mix by gently pipetting up and down to yield a concentration of 0.1 µg/µL. During use, the compound was diluted in a culture medium to achieve a final concentration of 2 µg/mL for cell treatment. Y-27632 was purchased from Yeasen Biotechnology and is provided as a powder with a purity exceeding 98 % (55,56). The powder should be aliquoted and stored at -20 °C in a dry environment for no more than 2 years. To prepare a stock solution, 10 mg of Y-27632 was dissolved in 312 µL of DMSO to create a 100 mmol/L solution, which was then diluted 10-fold with DMSO to achieve a concentration of 10 mmol/L. This solution was stored at -20 °C. For cell treatment, the solution was further diluted in culture medium to a final concentration of 30 µM.

### Construction of monoclonal genetically modified cell lines

We utilized overexpression lentiviral vectors to add fluorescent signals to the target protein for visualization imaging (57), RNA interference (58–62), and of target proteins (63). The lentiviral vectors used in this study were constructed and synthesized by GeneChem (https://www.genechem.com.cn/about.html). We packaged the synthesized vectors into lentivirus. The GeneChem lentiviral system consists of the GV lentiviral vector series, pHelper 1.0, and pHelper 2.0 plasmids. After mixing the plasmids, we performed transient transfection into HEK293T cells using the E-trans DNA transfection reagent (GeneChem, 10038555). Following gentle mixing, the cells were incubated for 48 hours, and the supernatant was collected. Lentivirus was purified using the GeneChem lentiviral particle purification kit and stored at -80 °C. The day before infection, cells were counted, and approximately 10,000 cells were seeded in a 12-well plate. After an overnight culture, the virus was diluted in serum-free medium and added to the plate to infect the cells at the recommended MOI. After 12 hours, the virus-containing solution was removed, and a complete medium was added for cell culture. After 48 hours, fluorescence intensity was observed under a fluorescence microscope. Corresponding antibiotics were added for cell selection, and monoclonal clones were picked for subsequent experiments. Five stable cell lines are constructed for this study: HEK293T-LifeAct-H2B to visualize F-actin and H2B, HEK293T-FUCCI to visualize cell cycle, HEK293T-DSG2OE to overexpress DSG2, HEK293T-DSG2RNAi to knockdown DSG2, and HEK293T-ITGB3RNAi to knockdown ITGB3. More details about the construction procedure and characteristics of the cell lines are offered in the Supplementary Information.

### Immunostaining Experiments

For the samples in 3D, the culture medium was aspirated, and the samples were washed with PBS. They were then fixed at room temperature with 0.4 % glutaraldehyde (National Medicines, 30092436) for 15 minutes, followed by another wash with PBS. Next, the samples were treated with 0.1 % sodium borohydride (Kemiou, 80115818KMO) at 4 °C for one hour to eliminate the effects of glutaraldehyde on imaging. After washing with PBS again, the samples were permeabilized at room temperature with 0.1 % Triton X-100 (Solarbio, T8200-500ml) for 30 minutes. They were then blocked at room temperature for 2 hours using a solution of 3% BSA (Pumeike, PMK0181-250g) prepared in 0.1 % Triton X-100. The samples were incubated overnight at 4°C in the dark with Alexa Fluor™ 488 phalloidin (Thermo Fisher, A12379) diluted 2000-fold in 3 % BSA for actin (64). The diluted antibody was incubated with the 3D samples overnight at 4 °C. After washing twice with PBS, an anti-fade mounting medium containing DAPI (Pumei Biological, PMK0272) was added, and the samples were incubated at room temperature for 15 minutes before imaging.

For the samples in 2D, the culture medium was aspirated and washed once with PBS. They were then fixed at room temperature with 4% paraformaldehyde (Pumei Biological, PMK2040) for 15 minutes, followed by another wash with PBS. The samples were permeabilized at room temperature with 0.5% Triton X-100 (Solarbio, T8200-500ml) for 10 minutes and then washed again with PBS. Next, a blocking solution of 3% BSA (Pumei Biological, PMK0181-250g) prepared in PBS was applied for 1 hour at room temperature. The p53 primary antibody (Abcam, catalog no. ab32389) was diluted 500-fold with 3 % BSA. The YAP1 primary antibody (Abcam, catalog no. ab205270) was diluted 500-fold with 3 % BSA. On the following day, the antibody was removed, and the samples were washed three times with PBS, each wash lasting 10 minutes. The Goat anti-Mouse IgG (H+L) secondary antibody was then diluted 1000-fold in 3% BSA and incubated with the samples for 2 hours at room temperature in the dark. Finally, the samples were washed three times with PBS, each wash lasting 10 minutes (65,66). After washing twice with PBS, Hoechst 33258 (Pumei Biological, PMK0965) was added for a 30-minute incubation at room temperature to stain the DNA (67).

### DNA FISH

Cells were cultured under three-dimensional conditions in normoxic or hypoxic (2% O□) environments for 12□h or 48□h, after which the medium was removed, and the cultures were washed once with PBS. Subsequently, 1□mL of 0.25% trypsin was added, and the matrix network was dissociated by gentle pipetting, followed by digestion at 37□°C for approximately 10□min until cell spheroids were dispersed into single cells. Digestion was terminated by adding 1□mL of culture medium. The cell suspension was transferred to a microcentrifuge tube and centrifuged at 1000□rpm for 5□min. The supernatant was discarded, and the pellet was resuspended in 0.075□M KCl, transferred to a 15□mL centrifuge tube, and diluted with 10□mL of 0.075□M KCl. After gentle inversion to mix, the tube was incubated at 37□°C for 30□min. Next, 2□mL of freshly prepared fixative (methanol: glacial acetic acid, 3:1) was slowly added to the suspension, mixed by inversion, and centrifuged at 1500□rpm for 10□min. The supernatant was removed, leaving approximately 0.5–1.0□mL above the pellet, which was gently resuspended. Fixative (5□mL) was slowly added, mixed, and the sample was centrifuged again under the same conditions. After repeating this washing step, the pellet was resuspended in fixative and adjusted to an appropriate concentration for slide preparation; fixed samples could be stored at –20□°C for up to 12 months. Subsequently, 7–10□μL of the cell suspension was vertically dropped onto pre-cleaned microscope slides and air-dried or oven-dried. Slides were then washed by immersion in 2× SSC for 5□min at room temperature, repeated once, followed by dehydration through a graded ethanol series (70%, 85%, and 100%, 2□min each). For hybridization, 10□μL of pre-mixed and centrifuged probe was applied to the target area under light-protected conditions, covered with a coverslip, avoiding bubbles, and sealed along the edges. After the sealant dried, slides were hybridized in a hybridization oven (or humidified chamber) under the following conditions: denaturation at 82□°C for 10□min and hybridization at 45□°C for 2□h. Post-hybridization, the sealant was carefully removed with forceps, and slides were immersed in ddH2O to gently lift off the coverslip. Slides were then washed in pre-warmed (68□°C) wash buffer (0.3% NP-40/0.4× SSC) for 2–6□min, rinsed in ddH2O for 1□min, immersed in 100% ethanol for 1□min, and air-dried. Finally, samples were counterstained with DAPI, mounted, and observed under a microscope.

### Live cell dye staining experiments

SiR-actin (CY-SC001, Cytoskeleton) is a fluorescent probe based on Silicon Rhodamine (SiR) conjugated with the actin-binding natural product Jasplakinolide. This combination enables specific and low-background labeling of F-actin in live cells (68). Hoechst 33258 was diluted 1000-fold in culture medium and incubated at 37 °C for 1 hour (67). A 1 mM SiR-actin or Hoechst 33258 stock solution was prepared in anhydrous DMSO or PBS and stored at -20 °C. For labeling, 3D-cultured cancer cell spheroids were collected, and the SiR-actin or Hoechst 33258 stock solution was diluted to 500 nM in culture medium. The cells were then incubated at 37 °C with 5% CO□ for 1 hour, with the incubation time adjustable based on the staining outcome. For cell lines with high levels of efflux pump expression, the addition of 10 µM Verapamil to the staining solution may improve the staining quality. After incubation, imaging was performed using a live-cell workstation with a spinning-disk confocal microscope. The laser wavelength was set to 640 nm (100 mW) with 20% laser intensity and an exposure time of 200 ms.

### Live cell imaging

For 3D imaging, we established a live cell imaging platform by integrating a 3-gas controller (Oko-lab, 3GF-MIXER-HYPOXIA) and a temperature incubator (Oko-lab, H301-MINI) onto a spinning disk confocal microscope (SpinSR, Evident Olympus). The microscope incubators maintain a temperature of 37°C for live cell culture, while the gas controller regulates the flow rates of air, carbon dioxide, and nitrogen to maintain a carbon dioxide concentration of 5% and an oxygen concentration of either 21% or 2%, depending on experimental requirements. Cell samples in a confocal dish (Nest, 801001) were placed on the live cell workstation. Typically, we set the laser intensity to 20% and the exposure time to 200 ms. To capture the dynamics of tumor spheroids, we utilized a laser with a wavelength of 488 nm to image the cytoskeleton and a laser with a wavelength of 561 nm to image the nuclei. To capture cell cycle, the 488 nm laser was used to image nuclei in the S/G2/M phases, while the 561 nm laser was employed to image nuclei in the G1 phase (69). To enhance experimental throughput, we selected a quad-well confocal dish (Cellvis, D35C4-20-1.5-N) for imaging, where each well operates independently without interference. Typically, three imaging positions are chosen within each well. When imaging 3D samples, at each position, a total 60 µm range along the z-axis was scanned at an interval of 5 µm, resulting in a total number of 15 images for one stack. When imaging 2D samples, images are captured at each position along the Z-axis at 3 µm intervals, resulting in a total of 7 images for one stack and a total z-range of 18 µm. The super-resolution images are obtained under the corresponding mode of the spinning disk confocal microscopy, using a 60x objective. Images were captured every hour for a total duration of 42 hours, with no significant phototoxicity observed (70–72).

### Data reconstruction and 3D visualization

The reconstruction of imaging stacks from confocal microscopy is conducted using Cell Sens Dimension Desktop 4.2.1 (https://lifescience.evidentscientific.com.cn/en/software/cellsens/). In the top planar view, the Maximum Intensity Projection (MIP) algorithm is adopted to overlay all images in the z-stack into a single image. In the side view, the Marching Cubes (Surface Reconstruction Algorithms) algorithm is used to yield the stereoscopic spheroids.

### Analysis of live cell imaging data

To analyze the spheroid, we segmented the tumor spheroids using CellPose (73), which provided the mask file of the spheroids. XY coordinates of each point on the spheroid’s edge are then calculated based on the mask data. Subsequently, using the segmentation data, we employed custom-written MATLAB code to analyze dynamic parameters, including area, perimeter, circularity, movement direction, and size of the tumor spheroids. Time-series image stacks were imported into ImageJ. The Tracking Plugin Manual Tracking (74) was utilized to obtain the movement trajectories, centroid spatial coordinates, distance, and speed of the tumor spheroids. To give a closer look into cells, we first segment the cells for one imaging using CellPose 2.0 (75), determining the optimal input parameters. Then, we utilized home-built Python code to perform batch segmentation for a series of image stacks. Once the batch-processed data were obtained, we re-imported them into CellPose for fine-tuning the segmentation parameters and obtaining the mask data of the cells. After acquiring the cell edge coordinates from the mask data, we employed home-built MATLAB scripts to analyze various properties of the cells, including centroid coordinates, area, roundness, fluorescence intensity, motility, and direction. Subsequently, statistical analysis and visualization were performed using OriginLab and ImageJ, respectively.

### Transcriptome sequencing and analysis

The tumor spheroids cultured in three dimensions are embedded in the matrix gel and cannot be separated individually. Therefore, when extracting RNA from these three-dimensional tumor spheres, we directly added Trizol into the matrix gel containing the tumor spheroid for extraction (76). In contrast, cells cultured in two dimensions can be lysed directly using Trizol. After culturing cancer cell spheroids under normoxic or hypoxic conditions for 42 hours, the culture medium was aspirated, and TransZol (TransGen, ET101-01-V2) was added directly to the mixture containing matrix gel and cells. RNA was then extracted and purified using the EasyPure® RNA Kit (TransGen, ER501-01-V2). The purified RNA was sent to Beijing Genomics Institution for library construction and sequencing, yielding sequencing data (77). We subsequently utilized R packages and Galaxy (https://usegalaxy.cn/) to align the sequencing data with human genomic information. To elucidate patterns and regulatory mechanisms of gene expression, regular analyses have been conducted, including sequence alignment, gene expression quantification, and differential expression analysis. Furthermore, we conducted GO and KEGG enrichment analyses of differentially expressed genes to gain deeper insights into the molecular mechanisms and regulatory networks underlying biological processes.

### Laser ablation experiments

To assess the mechanical property of tumor spheres, we employed a high-power laser to ablate individual cells within the tumor spheroids and monitored the response (78). Tumor spheroids cultured were first placed under a confocal microscope. Before laser ablation, ensure that tumor spheroids in different samples were at comparable z0 heights and had similar sizes. The intensity of the 405 nm laser was set to 100 % (50 mW), with a duration of 10 seconds for each ablation. A single cell at the edge of the tumor sphere was randomly selected for ablation. After laser exposure 4-5 times, the selected cell was ablated. During the ablation process, real-time imaging of the actin was conducted using a 488 nm laser, allowing for simultaneous live cell recording. Subsequent analysis focused on the deformation of the tumor spheroids over time.

### Nano-indentation measurements

The nano-indentation instrument (PIUMA Chiaro) can characterize the Young’s modulus of cell surfaces with high precision without deconstructing the cells (79). We measured the Young’s modulus of cell surfaces under both hypoxia and normoxia conditions. The sample preparation method was identical to that used for two-dimensional live-cell imaging. The confocal dish was placed under the microscope when the cell density reached approximately 70-80%. We first observe the sample under the objective, placing the interested cell at the center of the viewing field. Then a nanoindentation probe with the size of 2 µm is lowered onto the cell surface, giving feedback to the in situ force it senses. Nine random locations were selected on each confocal dish for measurement with the same nanoindentation probe.

### Scanning electron microscopy

Cells were seeded in Matrigel and cultured under either 21% or 2% oxygen for 48 hours. After collection, the samples were washed once with PBS and fixed with 0.4% glutaraldehyde (National Medicines, 30092436) at room temperature for 15 minutes, which preserved both the spheroid morphology and the Matrigel network structure. Subsequently, the samples were processed for electron microscopy. They were immersed in a mixture containing 2% paraformaldehyde (PFA) and 2.5% glutaraldehyde at 4°C overnight, followed by three washes with 0.1 M cacodylate/PB buffer (pH 7.4) at 4°C for 10 minutes each. Post-fixation was performed with 2% osmium tetroxide and 1.5% potassium ferrocyanide on ice for 1 hour. The samples were then rinsed with 0.1 M cacodylate/PB buffer on ice for 30 minutes, and washed three times with ice-cold ddH2O for 5 minutes each. This was followed by treatment with 1% thiocarbohydrazide (TCH) at 40°C in the dark for 30 minutes, and subsequent washing with ice-cold ddH2O three times for 10 minutes each. The samples were then treated with 2% osmium tetroxide on ice for 1 hour, washed four times with ice-cold ddH2O for 10 minutes each, and stained with 1% uranyl acetate (UA) in ddH2O at 4°C overnight. After warming at 50°C for 2 hours, the samples were washed four times with ice-cold ddH2O for 10 minutes each, treated with lead aspartate at 60°C for 1 hour, and again washed four times with ice-cold ddH2O for 10 minutes each. Dehydration was carried out using a graded ethanol series, followed by two rinses with 100% acetone for 20 minutes each at room temperature. Finally, the samples were embedded in resin for further processing.

Resin-embedded tumor spheroids were mounted onto an aluminum pin and trimmed to a cubic shape using an ultramicrotome (Leica UC7) until the tissue was exposed. Samples were subsequently coated with a 40-nm platinum layer using a high-vacuum sputter coater (Leica EM ACE600). The aluminum pin was then transferred to a Gatan 3View2 system within a serial block-face scanning electron microscope (SBF-SEM, Zeiss Gemini 300). Sample imaging was performed at an acceleration voltage of 1.5 kV with a 30μm aperture and a dwell time of 1 μs. Images were acquired at a resolution of 11,000× 11,000 pixels (5-nm pixel size), with serial sectioning at 70nm z-intervals.

## Acknowledgments

We thank Min Bao from Oujiang Laboratory for helpful discussions; Ruirui Gao and Yeqing Leng from Wuhan University for sharing monoclonal transfected cell lines; Chaoyong Liu from Wuhan University for his assistance in 3D image processing; Meng Shao, Limin Zheng, and Yilin Sun from Westlake University for scanning electron microscopy imaging; Yinping Gao from Westlake University and Jie Wang from Oujiang Laboratory for STED imaging; Danyan Yang from CSR biotech for SIM imaging; and ChatGTP for grammar checking and text polishing.

## Funding

National Natural Science Foundation of China (NSFC): Bo Li Distinguished Young Scholars Grant (Overseas)

Wuhan University (WHU): Bo Li Talents Startup Funding

National Natural Science Foundation of China (NSFC): Bo Li Young Scientists Fund (C Class)

## Author contributions

Conceptualization: BL

Methodology: BL, ZDW

Investigation: BL, ZDW, LT

Visualization: ZDW

Funding acquisition: BL

Project administration: BL

Supervision: BL

Writing – original draft: BL, ZDW

Writing – review & editing: BL, ZDW, LT

## Competing interests

The Authors declare that they have no competing interests.

## Data and materials availability

All data are available in the main text or the supplementary materials.

---

**Fig. S1.**
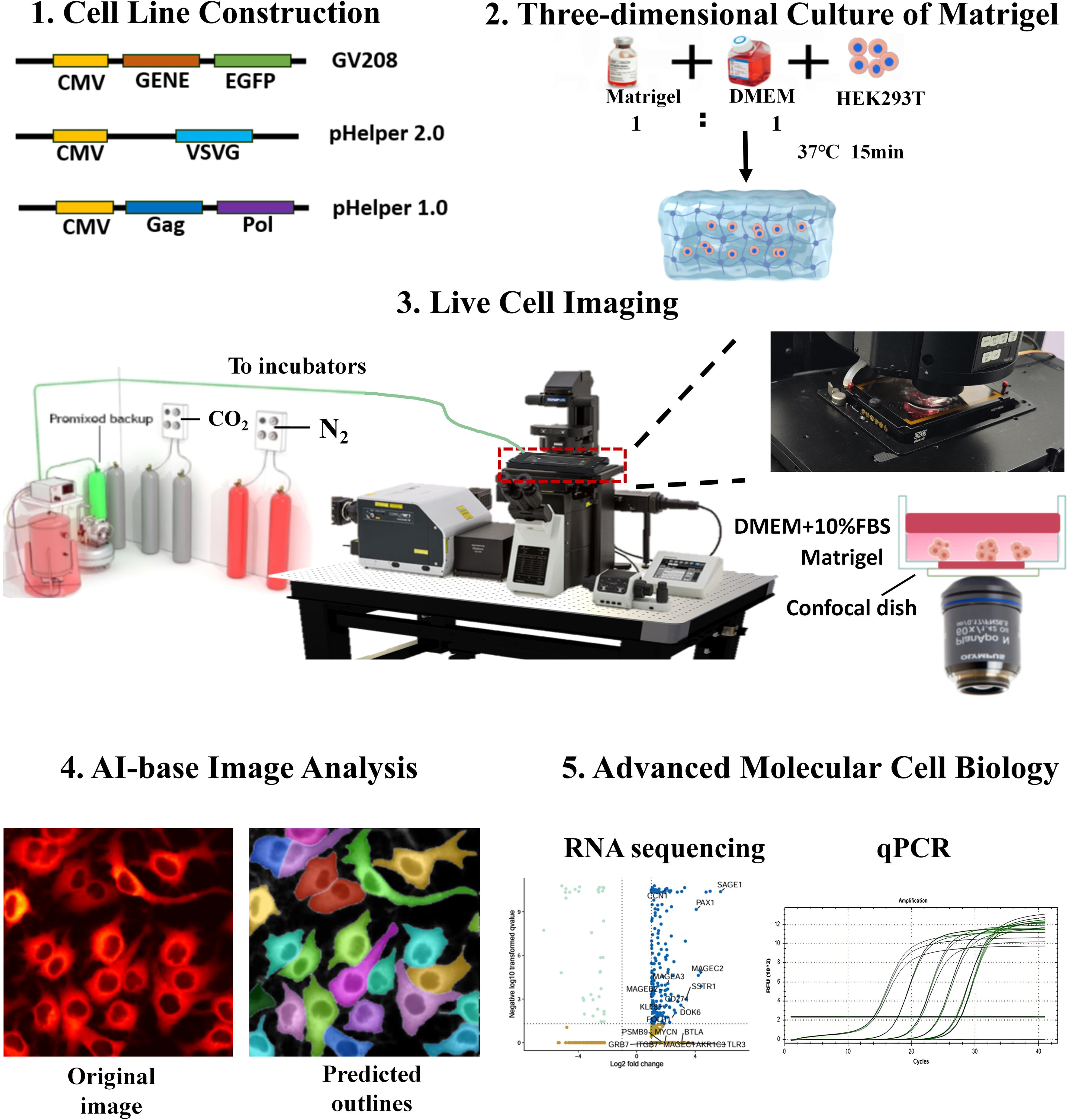

**Fig. S2.**
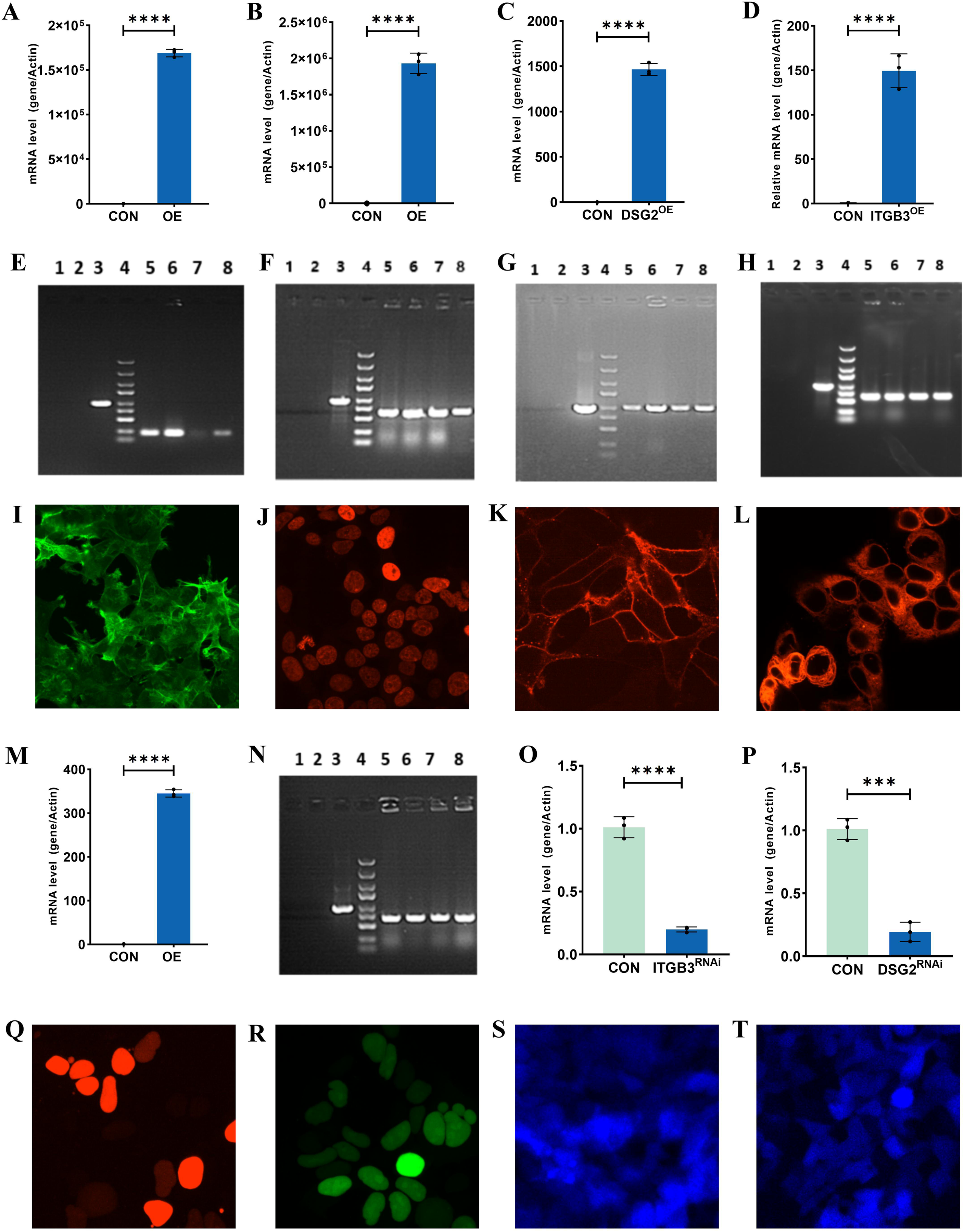

**Fig. S3.**
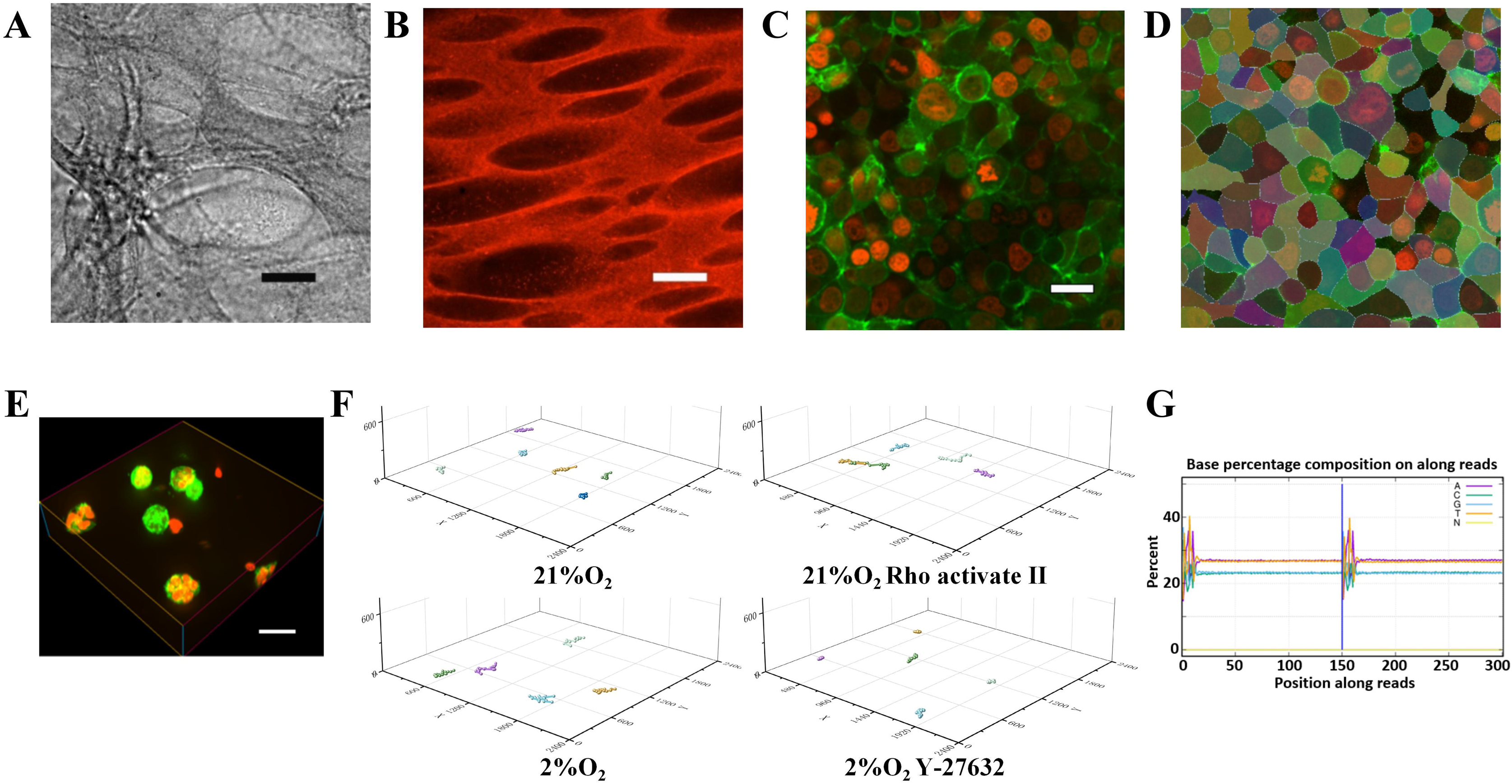

**Fig. S4.**
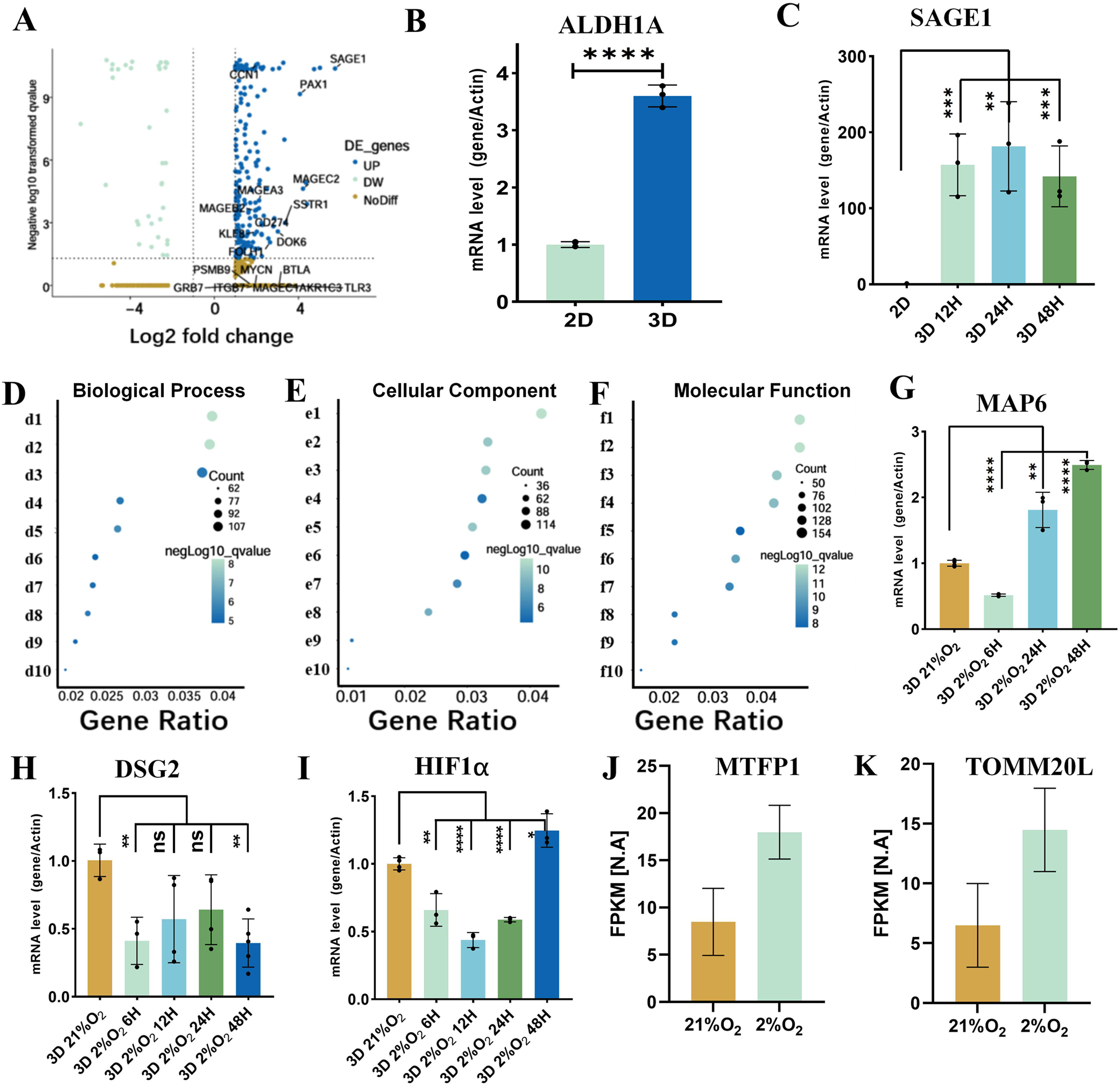

**Fig. S5.**
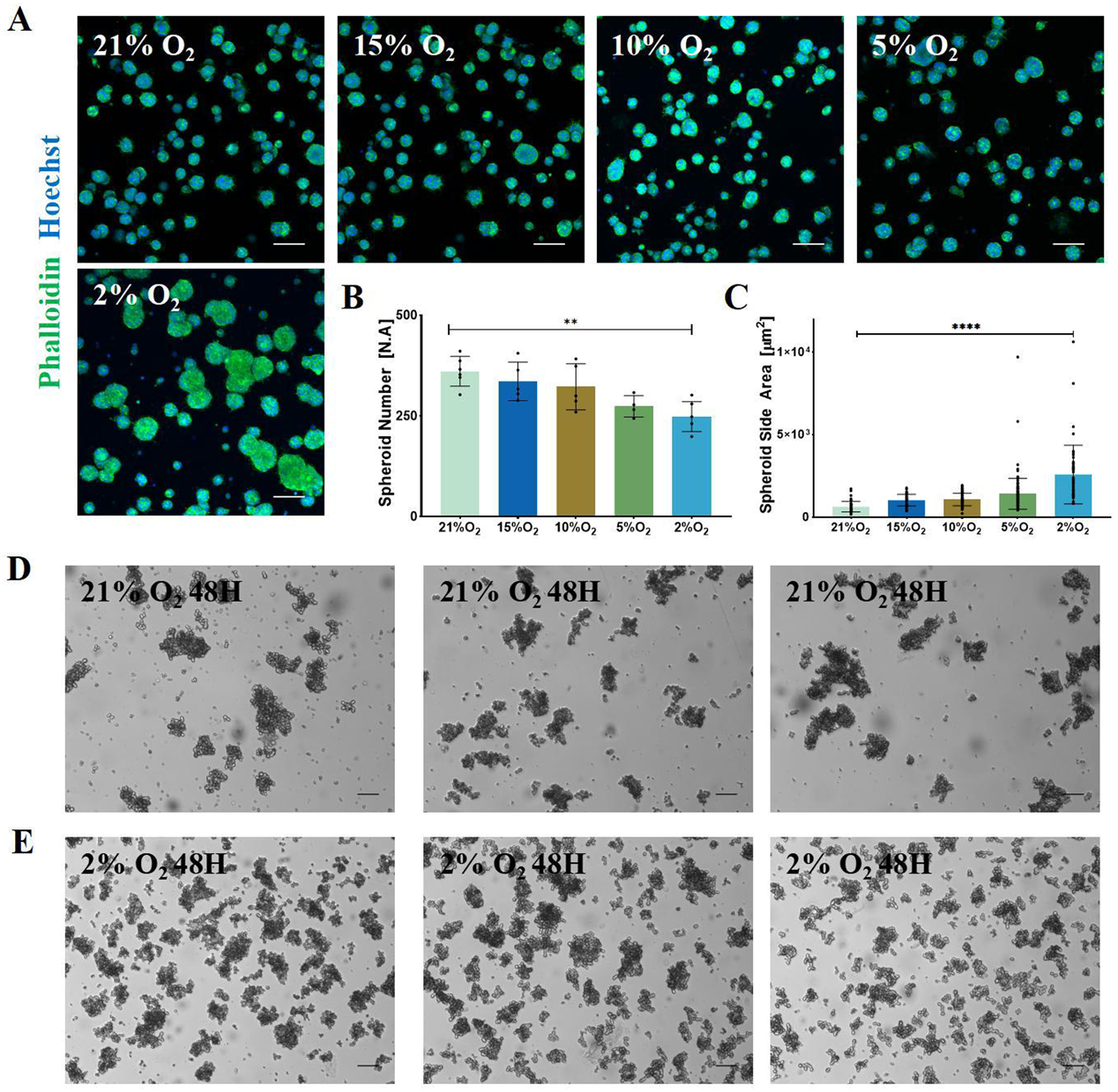

**Fig. S6.**
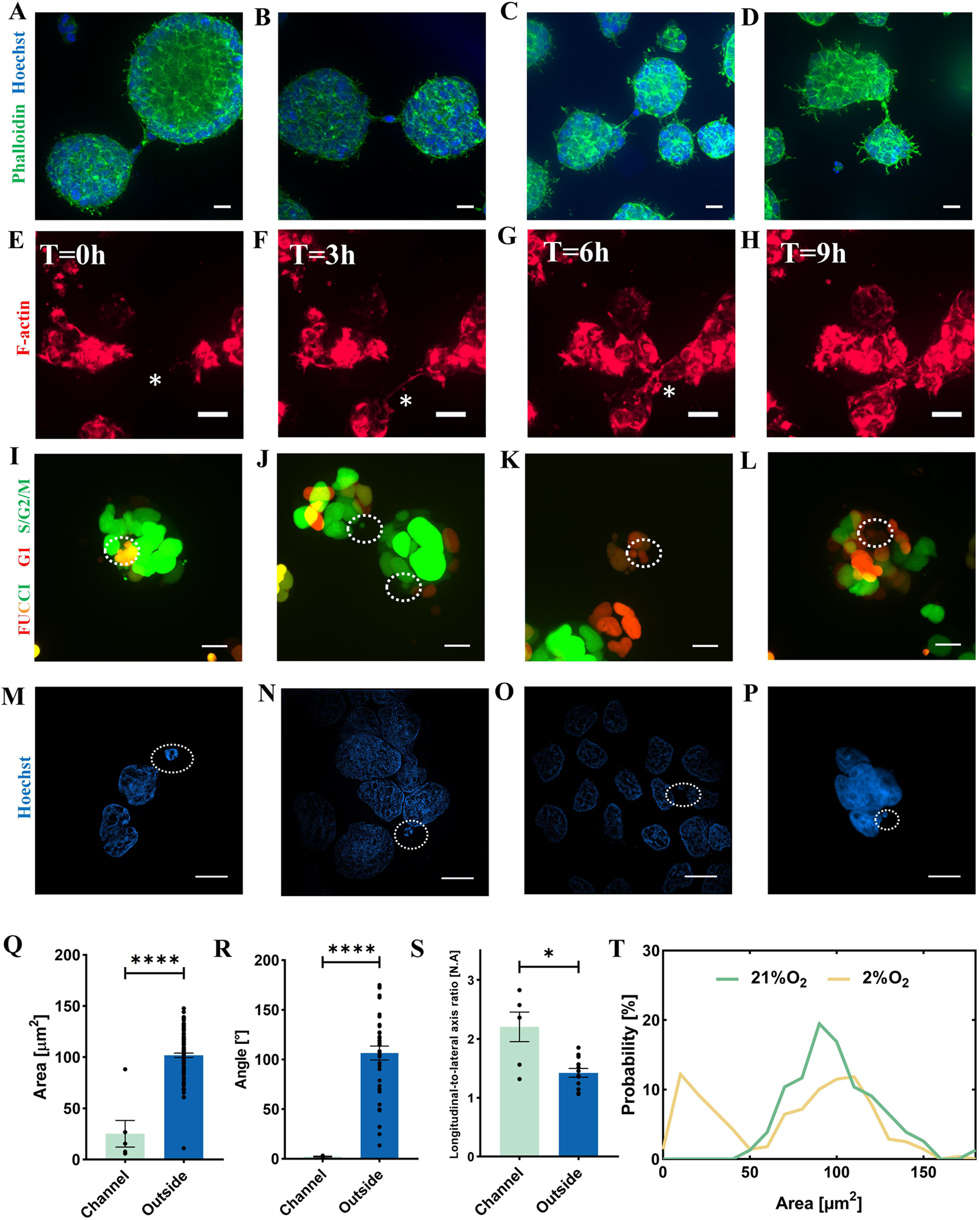

**Fig. S7.**
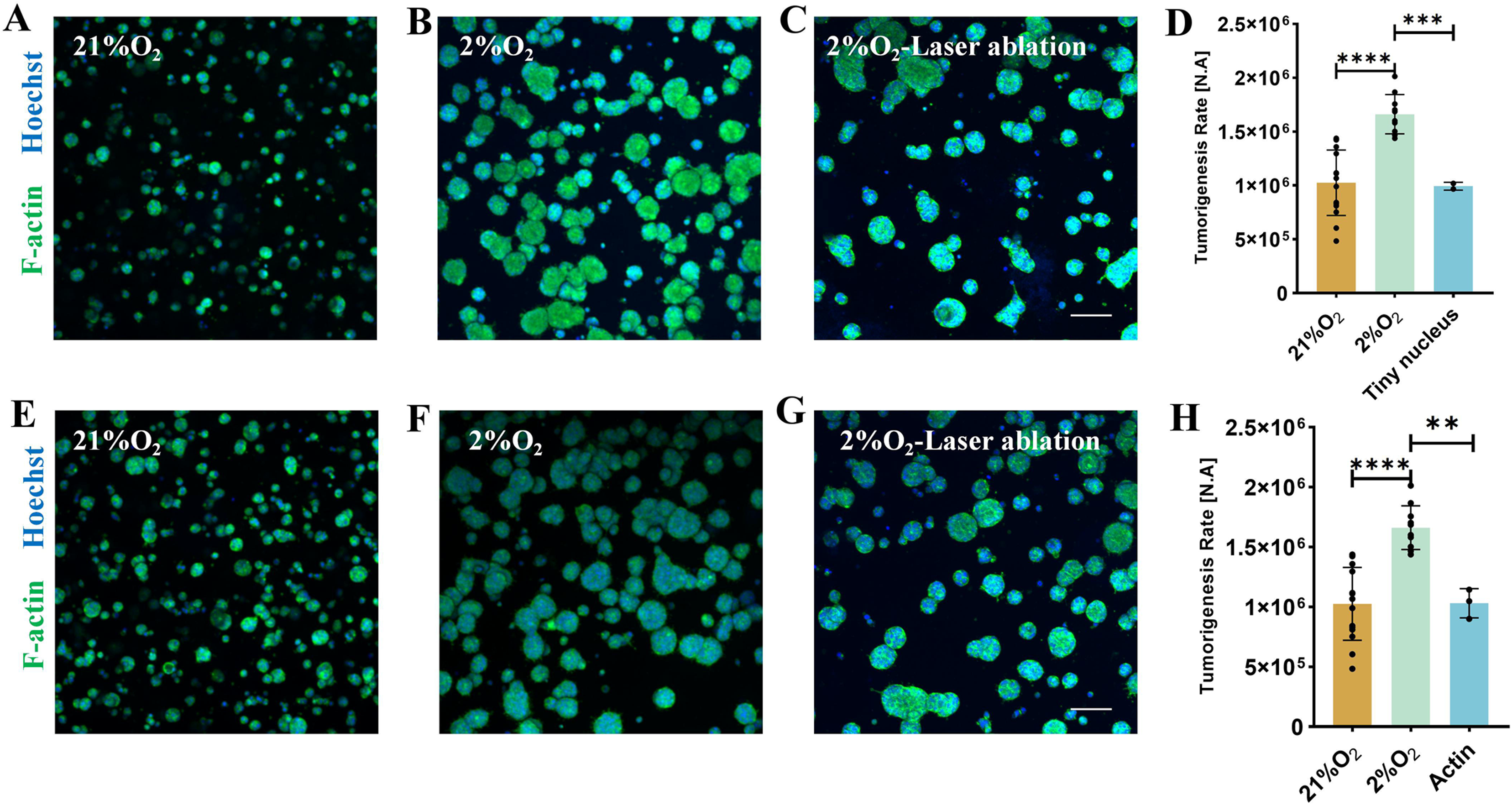

**Fig. S8.**
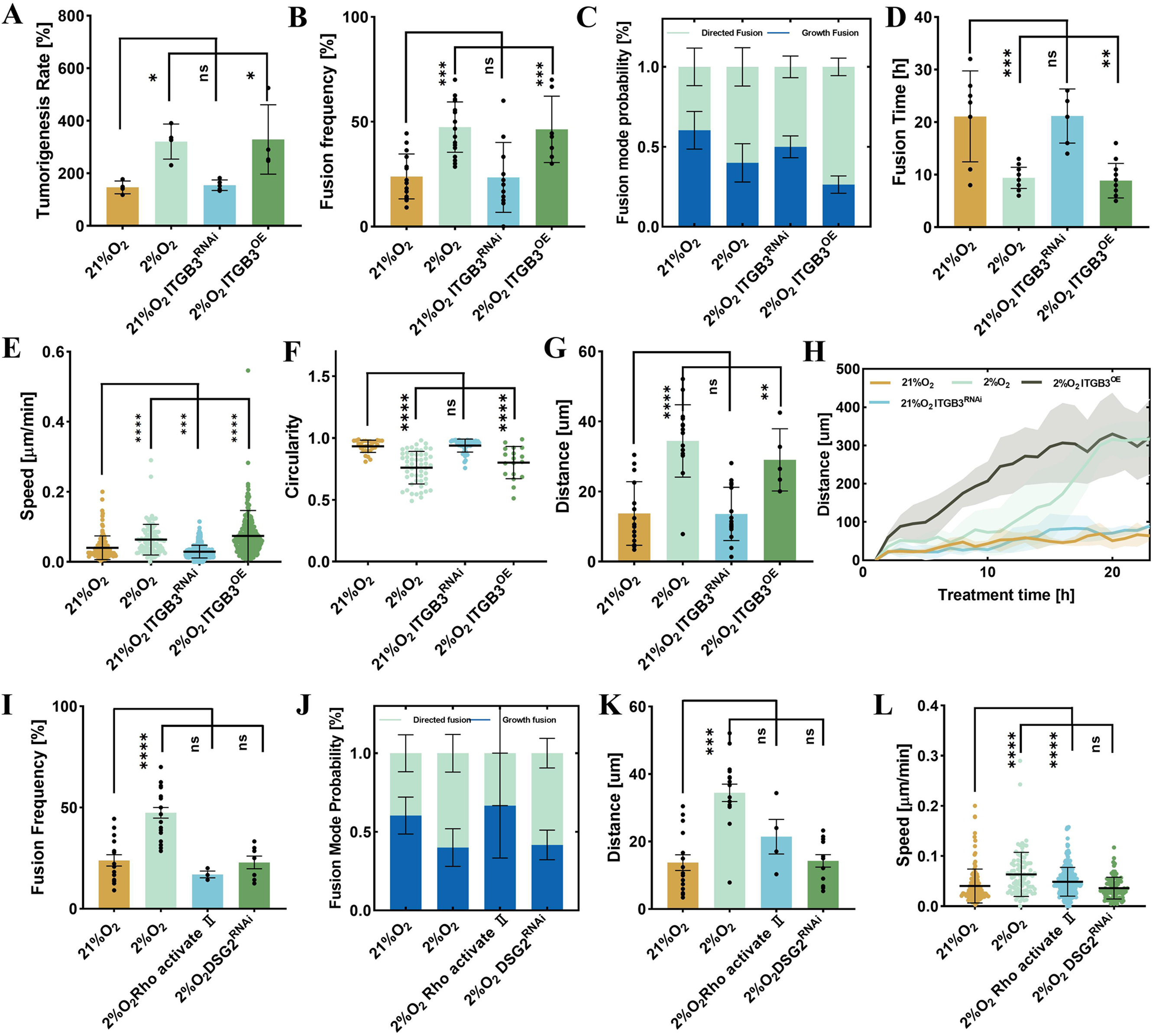

**Fig. S9.**
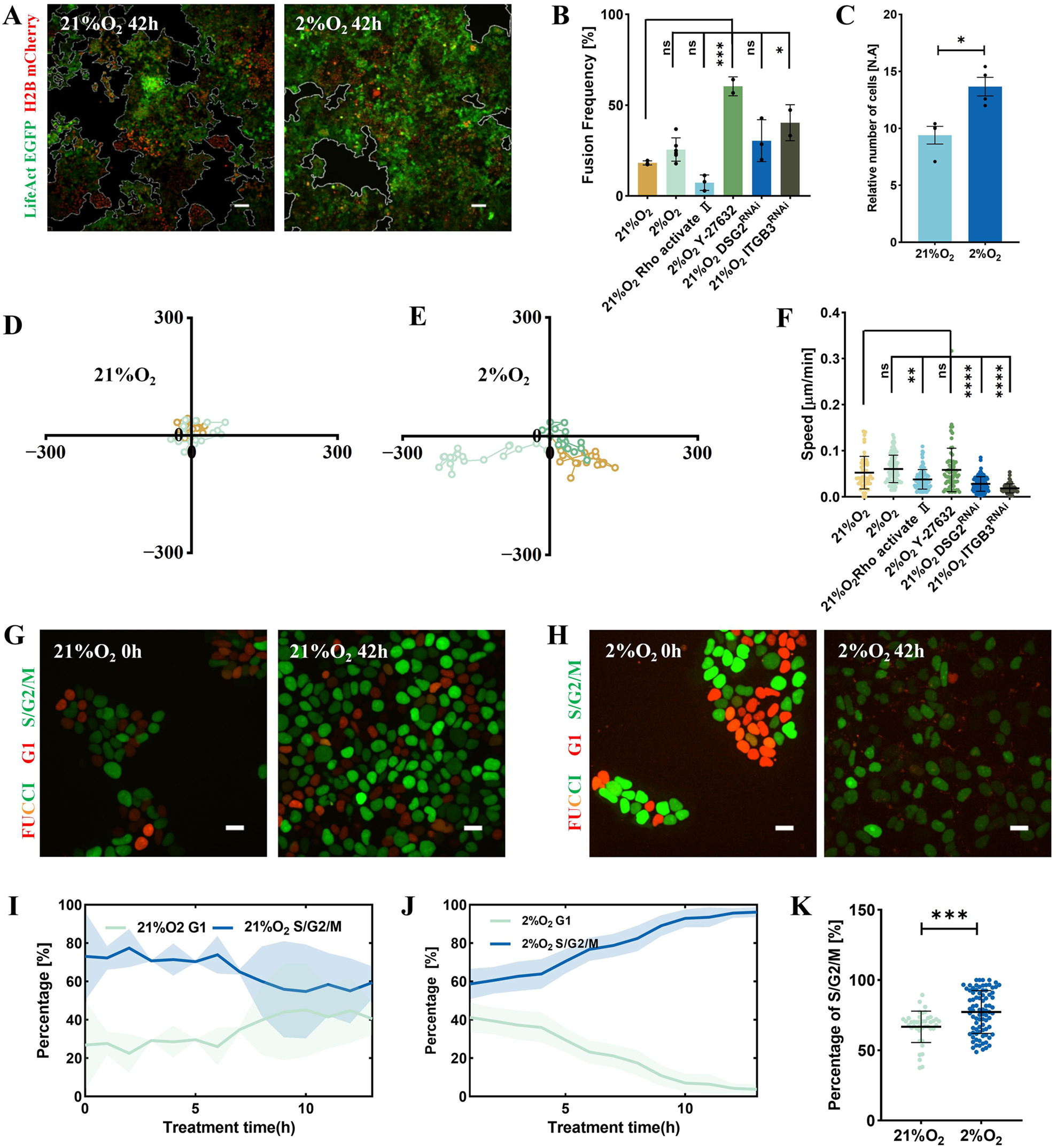

**Fig. S10.**
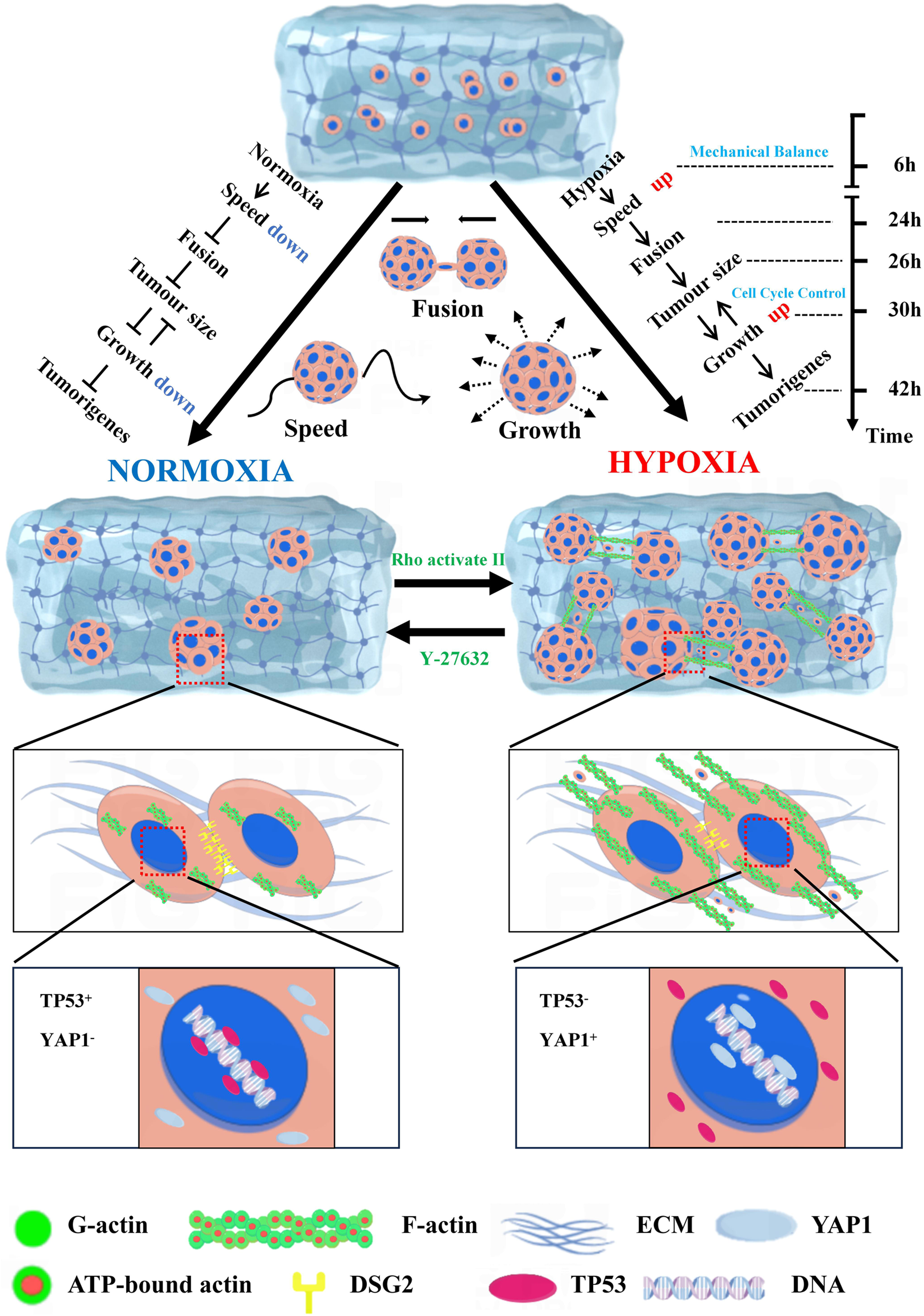

**Fig. S11.**
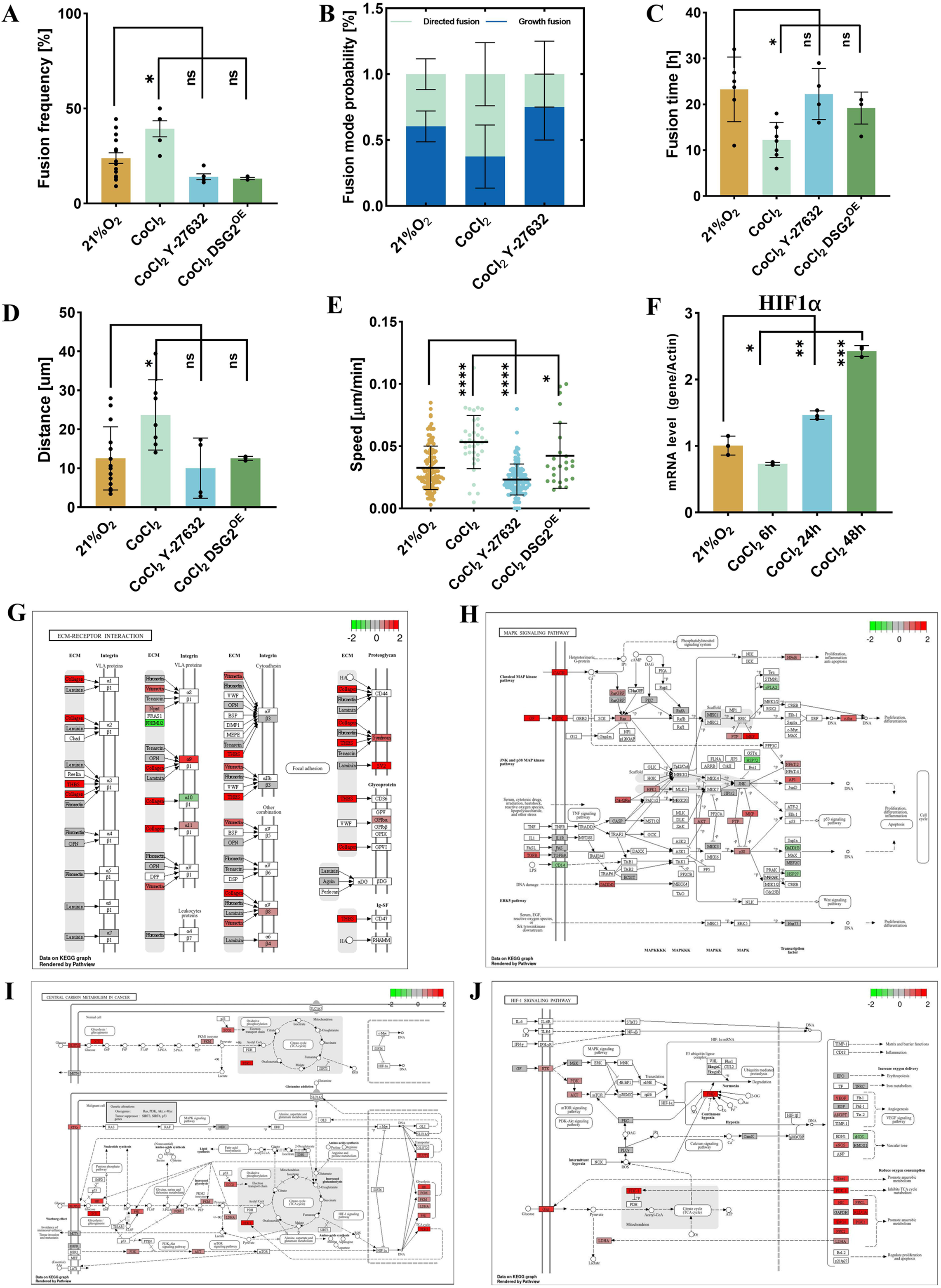

**Fig. S12.**
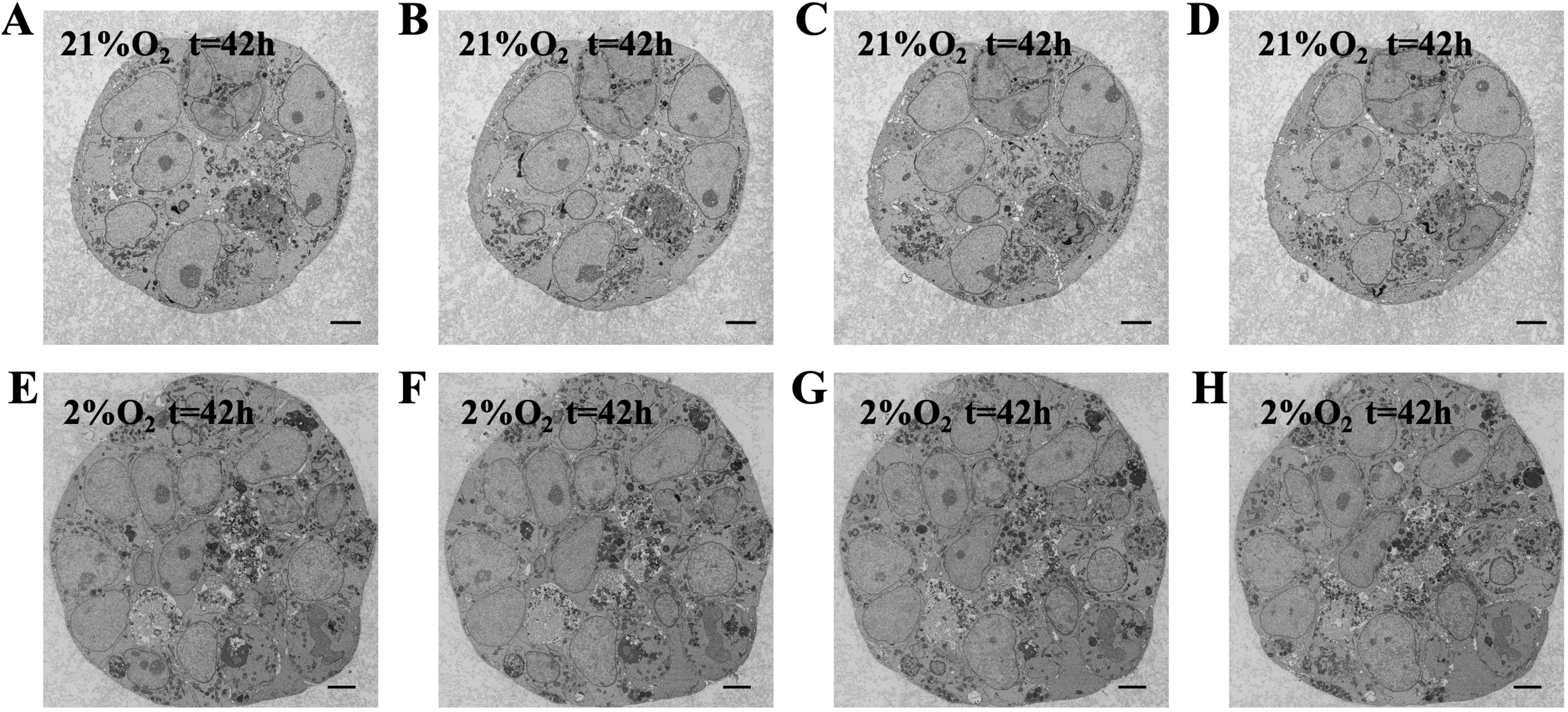

## Notes

### Competing Interest Statement

The authors have declared no competing interest.

### Summary of Updates

Supplementary videos are added in this revision

