## Supplementary Information for "Hypoxia Promotes Early Tumorigenesis through Dynamical-Mechanical Balance and Genetic Instability"

**Research Article**

**This PDF file includes:**

Supplementary Text

Figs. S1 to S12

Tables S1 to S5

Movies S1 to S10

References

**Other Supplementary Materials for this manuscript include the following:**

Movies S1 to S10

**Supplementary Text**

I. **Construction and characterization of cell lines.**

To achieve the visualization of target proteins and validate their related functions, we modified cell lines at the genetic level using lentiviral vector infection (1-3), the procedure and outcome of which are summarized in this section (Fig. S2). This included randomly inserting a green fluorescent Lifeact into the genome, which expresses green fluorescence and binds to F-actin in cells, thereby enabling visualization of cellular F-actin. Additionally, we achieved visualization of the nucleus by inserting the chimera gene with H2B and a red fluorescent, mCherry, into the genome (We visualized the cell nuclei by infecting cells with lentiviral vectors expressing exogenous H2B and red fluorescence). For the overexpression of DSG2 and ITGB3, we introduced these genes as exogenous genes through random insertion, utilizing a broad-spectrum promoter. As a result, the mRNA levels of the target genes in the overexpressing cell lines were upregulated by at least 100-fold (Fig. S2C and D). For RNA interference (RNAi) targeting DSG2 and ITGB3, we randomly inserted shRNA specific to the target proteins into the genome, facilitating continuous production of RNAi against the target genes. Consequently, the mRNA levels of the target genes in the RNAi cell lines were downregulated by at least 70 % (Fig. S2O and P).

The lentiviral vector used in this study was commercially constructed and synthesized (GeneChem Ltd, Shanghai). We packaged the synthesized vector into lentivirus. The lentiviral system consists of the GV lentiviral vector series, pHelper 1.0, and pHelper 2.0 plasmids. The GV lentiviral vector serves as the core of the viral genome and contains essential components of HIV, including the 5' LTR and 3' LTR, as well as viral packaging signals and other auxiliary elements such as the woodchuck hepatitis virus posttranscriptional regulatory element (WPRE). The GV vector can be modified according to specific experimental objectives to insert target genes or fragments for gene function studies (4,5). The pHelper 1.0 vector is an auxiliary plasmid that contains the gag gene, which encodes the major structural protein of the virus; the pol gene, which encodes virus-specific enzymes; and the rev gene, which encodes a regulatory factor that modulates the expression of the gag and pol genes. The pHelper 2.0 vector, also an auxiliary plasmid, contains the VSV-G gene derived from the herpes simplex virus, providing the envelope protein necessary for viral packaging (6,7). Linearized vectors were obtained through restriction enzyme digestion, and target gene fragments were prepared via PCR amplification. The amplification primers were designed to add homologous recombination sequences at their 5' ends, ensuring that the sequences at the 5' and 3' ends of the amplified product matched those of the linearized cloning vector. The reaction system was set up using the linearized vector and the amplified target gene product for the recombination reaction, achieving in vitro circularization of the linearized vector and target gene fragment. The recombinant products were directly transformed, and single clones from the plates were screened through PCR identification. Positive clones were sequenced and analyzed. Correctly cloned bacterial cultures were expanded, and plasmids were extracted to obtain high-purity plasmids for downstream virus packaging (7).

HEK293T cells were seeded in 10 cm culture dishes and incubated in a serum-free medium for 1 hour when the cell density reached approximately 70-80 % in preparation for virus packaging. The constructed plasmid, along with pHelper 1.0 and pHelper 2.0 plasmids, was mixed with serum-free medium and E-trans DNA transfection reagent (GeneChem, 10038555) and incubated at room temperature for 15 minutes. This mixture was then added to the cells, and supernatants were collected every 24 hours. The collected supernatants containing lentivirus were purified using Lenti-X™ Concentrator (PT4421-2). Specifically, 500 g of the viral suspension was centrifuged for 10 minutes, and the clarified supernatant was transferred to a sterile container. One volume of viral purification reagent was mixed with three volumes of the clarified supernatant. The mixture was gently inverted and incubated overnight at 4 °C. Following this, the mixture was centrifuged at 1500 g for 45 minutes at 4 °C, and the supernatant was carefully discarded. The viral pellet was resuspended in 1-2 mL of sterile PBS and stored at -80 °C.

On the day before infection, healthy cells were digested and seeded in 24-well plates, aiming for a cell density of approximately 20 % after 24 hours. Typically, 500 µL of viral solution was mixed with 500 µL of culture medium and subsequently added to the cells. After 12 hours of viral infection, the viral-containing medium was removed and replaced with a serum-containing medium for continued culture. Forty-eight hours later, fluorescence expression was observed under a microscope. Monoclonal cell lines were established for the purpose of avoiding genetic heterogeneity and ensuring a substantial strength of the fluorescence signals. To do this, we continued the cultivation in Petri dishes by adding an appropriate dose of culture medium containing specific antibiotics. The HEK293T-LifeAct cell line was selected using a medium containing G418, while the HEK293T-H2B and HEK293T-FUCCI cell lines were selected using medium containing puromycin. The HEK293T-DSG2 cell line was selected with a medium containing puromycin, and the HEK293T-ITGB3 cell line was selected with a medium containing puromycin. The HEK293T-DSG2 RNAi cell line was selected using a medium containing blasticidin, and the HEK293T-ITGB3 RNAi cell line was selected with a medium containing puromycin. Typically, the culture medium was replaced every two days until no significant cell death was observed. Subsequently, the cells were digested with trypsin to prepare a cell suspension, which was then diluted to an appropriate concentration and seeded into a 96-well plate, ensuring that only one cell was seeded per well. After 24 hours, once the cells had adhered, fluorescence signals were observed again using a confocal microscope. Cells exhibiting positive fluorescence signals were selected for further cultivation. When the cells approached confluence, they were digested with trypsin and reseeded into a 24-well plate for continued culture until a large number of monoclonal cells were obtained for preservation and experimentation. Throughout the entire process of monoclonal cell expansion, culture was performed using media containing lower concentrations of the corresponding antibiotics than those used during selection, until a substantial number of monoclonal cells were achieved (Fig. S2I-L and Fig. S2Q-T).

All cell culture and manipulations of cell lines (8) are conducted in a cell room that satisfies ISO 7 in the ISO 14644-1 system or Class 10,000 in the FS 209E system. The mycoplasma test (9) was done every week, and no positive result has been reported.

We verified the accuracy of the recombinant plasmid using PCR gel electrophoresis and assessed the mRNA expression of the target gene in the constructed cell lines via qPCR (Fig. S2E-H and Fig. S2N). All primers used for qPCR are listed in Table S1. Additionally, we confirmed the localization of the target protein in the cells through confocal microscopy.

The stable cell lines used in this study are summarized in Table S2, and we provide in this paragraph a detailed description of their nomenclature, characteristics, and excitation wavelength, etc. To enable dynamic visualization of the cellular processes, we constructed HEK293T cells expressing LifeAct-EGFP and H2B-mCherry (HEK293T-LifeAct-H2B). LifeAct is a small peptide that specifically binds to F-actin, thereby labeling the actin structures. This binding makes LifeAct an important tool for studying cytoskeletal dynamics. By fusing LifeAct with enhanced green fluorescent protein (EGFP), we can visualize actin filaments. The primary excitation wavelength for EGFP is approximately 488 to 490 nm. The mCherry fluorescent protein is fused with the histone H2B to enable visualization of chromatin. The primary excitation wavelength for mCherry is around 550 to 563 nm. To investigate cell cycle dynamics, we constructed HEK293T cell lines that express the FUCCI system (HEK293T-FUCCI), a dual-color fluorescent probe. This system combines mKO2 (red) and mAG (green) fluorescent proteins with two cell cycle-dependent proteins—Cdt1 and Geminin—allowing us to indicate whether individual live cells are in the G1 phase or the S/G2/M phase. The excitation wavelength for the red fluorescent protein mKO2 is typically around 530 to 550 nm, while the excitation wavelength for the green fluorescent protein mAG is approximately 488 to 495 nm. To investigate the role of intercellular communications and the cell-ECM interaction, we constructed different cell lines that overexpress DSG2 and ITGB3, or reduce their expression through RNA interference. For the overexpression of DSG2(HEK293T-DSG2OE), we built a vector that fuses DSG2 with mCherry and integrated it into the HEK293T genome, thereby simultaneously enhancing DSG2 expression and enabling its visualization. The overexpression strategy for ITGB3 (HEK293T-ITGB3OE) was similar to that of DSG2. For the reduction of DSG2 expression (HEK293T-DSG2RNAi), we constructed a separate cell line that expresses shRNA fused with EBFP targeting DSG2. After being integrated into the HEK293T genome, the sustained expression of the interfering RNA results in decreased DSG2 levels. The method used to lower ITGB3 expression (HEK293T-ITGB3RNAi) was analogous to that employed for DSG2.

II. **Porous ECM gel for 3D culture.**

The porous ECM structure formed from the Matrigel provides the 3-dimensional microenvironment for the tumor cell to survive and proliferate (Fig. S3). This method has been widely used to reproduce 3D cellular processes in vitro, including organoid culture, internal angiogenesis experiments, and tumor growth in nude mice (10-12). To elucidate the microenvironment of the tumor spheroid, we directly image the ECM structure by bright field and confocal microscopy. Empty porous with 40 μm size are homogeneously distributed throughout a 3D Matrigel network (Fig. S3). The size of the pore is much larger compared to that of a typical tumor spheroid, offering sufficient space for the spheroid to move and grow. So any confinement force from the ECM can be excluded. In other words, the ECM serves as a gentle strut for the 3D culture but has little effect on spheroid behavior.

Here we provide more details of fabricating the ECM structure with the appropriate porous size. Mix the Matrigel and DMEM basal medium in a 1:1 ratio and place the mixture on ice. For a 35 mm confocal dish, add approximately 80 µL of the prepared Matrigel; for a four-well confocal dish, add 20 µL of Matrigel to the bottom of each well. Use a pre-chilled 10 µL sterile pipette tip to spread the Matrigel evenly. The amount of Matrigel applied should be appropriate; too much may reduce imaging depth, while too little can lead to cell adhesion without spheroid formation. After using the Matrigel, place the confocal dish at 37 °C for approximately 10 minutes to allow it to solidify. Next, aspirate the cell culture medium and wash the cells twice with PBS, reserving 1 mL of PBS. Add 0.5 mL of 0.25 % trypsin to digest the cells. When the cells round up and begin to detach, add 1 mL of serum-containing medium to terminate digestion. Gently pipette to create a homogeneous cell suspension, transfer the suspension to a centrifuge tube, and centrifuge at 1400 rpm for 5 minutes. Resuspend the cells in the prepared Matrigel solution and add them to the confocal dish containing the Matrigel. Incubate at 37 °C for 10 minutes to allow solidification. Finally, add DMEM medium containing 10 % serum and incubate at 37 °C with 5 % CO₂.

**III. Time-dependent molecular analysis on the function of cancer stemness, hypoxia factor, and cell mechanics.**

Early detection of cancer significantly enhances treatment efficacy and improves survival rates (13-15). However, over 50% of cancers are diagnosed at advanced stages. The progression of tumors follows a continuum from normal tissue to dysregulation, carcinogenesis, proliferation, differentiation, and metastasis. By the time cancer is clinically detected, cancer cells have undergone sufficient genetic and phenotypic alterations, leading to challenges such as drug resistance and recurrence. Therefore, targeted interventions during the early stages of cancer development may yield better anti-tumor outcomes.

One critical challenge in early cancer detection is the lack of research into the early biological characteristics of cancer (13,16), primarily due to the difficulty in observing the first cancer cells that appear in tissues in vivo. We propose an ex vivo model to simulate the tumor microenvironment for studying early cancer biology. The HEK293T cells, when transplanted into three-week-old mice, form tumors within 14 days (17). Histologically, tumors formed by 293T cells are consistent with poorly differentiated breast cancer, exhibiting high expression of vimentin and β-catenin, along with low expression of E-cadherin and cytokeratin (18), which aligns with the expression profile of poorly differentiated tumors (17). In three-dimensional (3D) culture, 293T cells in spheroids display a higher proportion of ALDH1 positive cells and CD44+/CD24- cell populations compared to those in two-dimensional (2D) monolayer cultures, both of which are markers of cancer stem cells. The cells within the 3D spheroids exhibit radio-resistance and upregulate signaling pathways associated with stem cell survival, including the expression of β-catenin, Notch1, and Survivin under radiation exposure. Additionally, spheroid cells show increased expression of mesenchymal genes (vimentin, N-cadherin, ZEB1, Snail, and Slug) as well as pro-metastatic genes (RhoC, Tenascin C, and MTA1) (18,19). Transcriptomic sequencing and qPCR results further demonstrate significant upregulation of cancer embryonic antigen-related genes SAGE1, MAGEC2, and ALDH1A (Fig. S4). After forming spheroid in 3D culture, the 293T cells exhibit typical cancer stem cell characteristics, including self-renewal ability, treatment resistance, and metastatic potential. This cellular model serves as a crucial tool for investigating the molecular and biological mechanisms of cancer stem cells and will aid in the development of novel therapeutic strategies targeting cancer stem cells.

Data on early tumor growth dynamics indicate that under hypoxia conditions, tumor spheroids promote fusion between each other by facilitating actin polymerization. RNA sequencing results show that genes regulating cytoskeletal dynamics are upregulated under hypoxia (Fig. S4). We validated the expression levels of MAP6, a key regulator of the cytoskeleton, at different time points after hypoxia using qPCR, and found that MAP6 expression levels increased following hypoxia (Fig. S4G). We further examined the mRNA expression levels of SAGE1, an embryonic antigen-related gene, at different time points during 3D culture. The results demonstrated that SAGE1 was persistently upregulated under 3D culture conditions. The longest time point we tested was 48 hours in 3D culture, at which time the mRNA level of SAGE1 remained more than 100 times higher compared to 2D culture (Fig. S4C). Our findings suggest that the weakened intercellular interactions under hypoxic 3D culture conditions play a crucial role in the early oncogenic potential of cancer cells. We also measured the mRNA expression of DSG2 at different time points under hypoxic 3D culture conditions and observed a continuous downregulation of DSG2 expression at 48 hours under hypoxia (Fig. S4H).

Additionally, we assessed the expression of HIF1α at various time points after hypoxia, revealing dynamic changes in HIF1α mRNA expression (Fig. S4I). The expression of HIF1α declined by almost 30 % at 6 hours and 24 hours after hypoxia treatment, in sharp contrast with previous studies in 2D systems (20). It is until 48 hours, the HIF1α expression shows a slight increase compared to the control group. But at this time, the directional fusion and enhanced tumorigenesis have already finished. KEGG enrichment at 24 hours shows no sign of enriched HIF1α pathways either. So the time-dependent mRNA test excludes the possibility that the classical HIF1α pathway is responsible for the dynamics of the tumor spheroids.

**IV. Cell segmentation, imaging reconstruction, and data analysis.**

In this section, we provide more information about imaging and data analysis. To obtain the morphology and dynamics of each cell, the outcome image (Fig. S3C and D) of the overlay from CellSens software was imported into CellPose (21,22), where parameters were adjusted to segment the cells. For cell segmentation using CellPose, the "cell diameter" parameter in the segmentation settings was first adjusted according to the size of the cells to be identified. The appropriate fluorescence signal channel for segmentation was then selected, with other parameters left at their default values. Pre-conFig.d segmentation models in CellPose, such as "cyto," "cyto2," and "nuclei," were tested to assess their segmentation performance. The model yielding the best results was chosen for further processing using custom-written Python code for batch processing. The segmentation results were subsequently re-input into CellPose for validation, and any misclassified data were manually corrected. Incorrectly segmented image data can be corrected by re-importing the data into CellPose, where errors can be removed using Ctrl + left-click. The correct cell boundaries are then manually outlined, and the corrected data are saved. All data underwent manual correction to accurately capture the boundary information of each cell (Fig. S3C and D).

During the three-dimensional imaging of tumor spheroids, the full imaging depth of the microscope can reach 200 µm. A field of view in xy plane is 245 µm by 245 µm under 60x oil objective. Using the motorized sample stage, more than ten field of view are recorded simultaneously. The typical diameter of a tumor spheroid after 48 hour hypoxia treatment is 37.024 µm. So the throughput of our experiment can reach more than eighty spheroids per trial. Typically, we acquired video data at 3 µm per layer along the z-axis, resulting in totally 21 layers with an overall z-axis height of 60 µm. Consequently, for analyzing the three-dimensional movement trajectories of the tumor spheroids, we utilized the maximum projection data obtained by stacking the 21 z-axis layers. This method applies for all confocal images presented main text and extended data Fig.s. Another way of 3D reconstruction is to use surface reconstruction algorithms, giving out the side view of the spheroid (Fig. S3E). The side view reconstruction appears to have compromised spatial resolution compared with the top-down view because of the resolution inconsistency of confocal imaging method in xy plane (200 nm) and z scanning (450 nm).

The segmentation of the 3D spheroid follows the same procedure as that of the 2D cell segmentation. Following import of the multi-plane Z-stack images, 3D reconstruction and segmentation were performed using Cellpose (Fig. 4E). The reconstructed spatial information was compared against the 3D models generated by the spinning disk confocal microscope software CellSens (Fig. S3E). Gaps between Z-slices were algorithmically interpolated to derive complete 3D boundary coordinates, which were subsequently used for 3D trajectory analysis (Fig. S3F).

After segmentation using CellPose, the spatial coordinates of the boundaries and inner area of the ROI (Region of Interest) will be obtained, including the brightness of each pixel in the ROI. The ROI could be either a spheroid or cells, depending on the object we are looking into. For analyzing the movement speed and distance of the tumor spheroids, we employed the "Manual Tracking" plugin from ImageJ (23). After setting the appropriate time interval and pixel size, we manually marked the movement trajectories of the tumor spheroids. The more dedicated kinetic analysis was performed using a custom-written MATLAB code. Let the function (I(x, y)) represent the intensity or grayscale values at all positions within the segmented boundary. If (R) is a discrete region, then for each pixel location, the value is typically set to 1 (indicating that we are essentially counting the number of pixels). Thus, (A) can be understood as the total number of pixels within the region (R). The area (A) can be expressed as follows:

$\boldsymbol{A}=\sum_{(\boldsymbol{x},\boldsymbol{y})\in\boldsymbol{R}} \boldsymbol{I}(\boldsymbol{x},\boldsymbol{y})$ ……………………………	**Eq.S1**

The centroid (Cx, Cy) can be calculated using the following formula:

$\boldsymbol{C}_{\boldsymbol{x}}=\frac{\mathbf{1}}{\boldsymbol{A}}\sum_{(\boldsymbol{x},\boldsymbol{y})} \boldsymbol{x}\times\boldsymbol{I}(\boldsymbol{x},\boldsymbol{y})$ ……………………………	**Eq.S2**

$\boldsymbol{C}_{\boldsymbol{y}}=\frac{\mathbf{1}}{\boldsymbol{A}}\sum_{(\boldsymbol{x},\boldsymbol{y})} \boldsymbol{y}\times\boldsymbol{I}(\boldsymbol{x},\boldsymbol{y})$ ……………………………	**Eq.S3**

The perimeter of the ROI (Region of Interest) is denoted as:

$\boldsymbol{P}=\sum_{(\boldsymbol{x},\boldsymbol{y})\in\boldsymbol{B}} \boldsymbol{I}(\boldsymbol{x},\boldsymbol{y})$	………………………	**Eq.S4**

$(x,y)\in B$ represents the summation of all pixel coordinates within the boundary B.

The roundness of the ROI (Region of Interest) is defined as:

$\boldsymbol{C}=\frac{\mathbf{4}\boldsymbol{\pi A}}{\boldsymbol{P}^{\mathbf{2}}}$	……………………………	**Eq.S5**

where A is the area and P is the perimeter.

Velocity can be calculated by measuring the displacement over a unit time interval. The displacements in the x-direction and y-direction within that unit time are denoted as $\Delta\boldsymbol{X}$ and$\Delta\boldsymbol{Y}$, respectively.

$\Delta\boldsymbol{x}=\boldsymbol{x}_{\boldsymbol{t}}-\boldsymbol{x}_{\boldsymbol{t}-\mathbf{1}}$	…………………………	**Eq.S6**

$\Delta\boldsymbol{y}=\boldsymbol{y}_{\boldsymbol{t}}-\boldsymbol{y}_{\boldsymbol{t}-\mathbf{1}}$	…………………………	**Eq.S7**

Therefore, the velocity (v) can be expressed as:

$\boldsymbol{v}=\sqrt{{\Delta\boldsymbol{x}}^{\mathbf{2}}+{\Delta\boldsymbol{y}}^{\mathbf{2}}}$	………………………	**Eq.S8**

To calculate the direction of motion of the tumor sphere, we first compute the vector difference between the positions at two adjacent time points. This is expressed as:

$\vec{\boldsymbol{B}}=(\boldsymbol{x}-\boldsymbol{x}_{\boldsymbol{P}\mathbf{1}},\boldsymbol{y}_{\boldsymbol{P}\mathbf{2}}-\boldsymbol{y}_{\boldsymbol{P}\mathbf{1}})$	…………………………	**Eq.S9**

Let A be the velocity vector and B be the displacement vector. The cosine of the angle between the two vectors can be expressed as:

$\cos\boldsymbol{\theta}=\frac{\boldsymbol{A}\times\boldsymbol{B}}{\left| \boldsymbol{A} \right|\times\left| \boldsymbol{B} \right|}$	………………………………	**Eq.S10**

The projection of the velocity vector onto the direction of the displacement vector can be expressed as:

$\boldsymbol{projection}=\left| \boldsymbol{A} \right|\cos(\boldsymbol{\theta})$	……………………………	**Eq.S11**

**V. Verification and analysis of transcriptomic data**

To conduct the transcriptomic analysis, we extract the RNA from the cells First, we implanted cancer cells into a three-dimensional (3D) culture system. Then, the 3D-cultured cancer cells were placed in an incubator set at 37 ℃ with 5% CO2 and 21% O₂, or in a hypoxic incubator at 37 ℃ with 5% CO2 and 2% O2, depending on the experimental requirements, for 12 to 48 hours. After the treatment period, the culture dishes containing the cells were removed, the culture medium was aspirated, and the cells were washed once with PBS. RNA extraction was performed using the TransZol Up Plus RNA Kit (ER501-01-V2). The microcentrifuge tubes were removed from the -80 ℃ freezer and placed on ice until thawed. After thawing, 200 µL of RNA Extraction Agent, TransZol, was added into each 3.5 cm culture dishes for cell cultivation, and the sample was pipetted repeatedly until no visible precipitate remained in the lysate. The mixture was vortexed at room temperature for 5 minutes and then centrifuged at 10,000 g for 15 minutes at 4 ℃. At this stage, the sample separated into three layers: a colorless aqueous phase (top layer), a middle layer, and a pink organic phase (bottom layer). The RNA is located in the aqueous phase. The colorless aqueous phase (400 µL) was carefully transferred to a new centrifuge tube, and an equal volume of anhydrous ethanol was added. The solution was gently mixed by inverting the tube. The mixture, including any precipitate, was then transferred to a spin column and centrifuged at 12,000 g at room temperature for 30 seconds. The flow-through was discarded, and 500 µL of CB9 buffer was added. The sample was centrifuged at 12,000 g at room temperature for 30 seconds, and the flow-through was discarded. This step was repeated once. Then, 500 µL of WB9 buffer was added, and the sample was centrifuged at 12,000 g at room temperature for 30 seconds. The flow-through was discarded, and the washing step was repeated once more. Finally, the sample was centrifuged at 12,000 g for 2 minutes to completely remove any residual ethanol. The spin column was transferred into an RNase-free tube, and 40 µL of RNase-free water was added to the center of the column. The column was incubated at room temperature for 1 minute, followed by centrifugation at 12,000 g for 1 minute to elute the RNA. The extracted RNA was stored at -80 ℃.

The prepared RNA samples used in transcriptome sequencing were delivered to BGI Genomics for sequencing. The purity and integrity of the RNA were assessed through agarose gel electrophoresis in our laboratory before submitting the samples for further quality checks by the company. After passing the double-check procedures, the sequencing started. The library construction and sequencing process includes the following steps. First, the mRNA is extracted using oligo-Dt and the total amount of it is quantified. Then, the mRNA chain is fragmented by chemical method. The use of metal ions, such as zinc ions, can catalyze the cleavage of phosphodiester bonds within RNA molecules, thereby fragmenting the RNA into smaller segments. Random primers are added for the synthesis of single-stranded cDNA from the fragmented mRNA. With the synthesized double-stranded cDNA, a ploy-A tail and ligation of adapters were used for the end repair. The prepared cDNA further underwent PCR amplification, quality assessment, and circularization to establish the library. The circular DNA molecules were rolled up to generate DNA nanoballs (DNBs). Finally, the DNBs were sequenced on the DNBSEQ platform.

After sequencing, the raw data undergoes filtering to remove contaminants, adapters, and low-quality data. Some raw sequences may contain adapter sequences or a small number of low-quality bases. We perform a series of data processing steps to eliminate impurities and obtain high-quality data. This analysis is conducted using SOAPnuke (24), a filtering software developed by BGI. The filtering parameters for SOAPnuke are set as follows: "-n 0.001 -l 20 -q 0.4 --adaMR 0.25 --polyX 50 --minReadLen 150." The filtering steps are described as follows. An adapter filtering was conducted to discard any sequencing read matching an adapter sequence at 25.0 % or more (allowing for a maximum of two mismatches). Read length filtering was then applied, and reads shorter than 150 bp were removed. Removal of N bases discards reads that contain 0.1% or more N bases. PolyX filtering deletes reads with a polyX length (where X can be A, T, G, or C) exceeding 50 bp. If the proportion of bases with a quality score below 20 exceeds 40.0 %, the low-quality data filtering removes the entire read. Finally, the output reads are assigned quality values based on the Phred+33 scale. Following the filtering of the raw data, we monitor data quality to ensure the integrity of the sequencing results (Fig. S3G).

To support opinions in the corresponding parts of the main text, we provide here a more detailed analysis of the transcriptomic data. Transcriptomic data show substantial differences in the gene expression pattern between the HEK293T cells in 3-dimensional and 2-dimensional cultures. In 3D, a significant number of genes that control cancer stemness are upregulated (Table S3). For example, we found SAGE1 (Squamous Aggressive Growth Enhancer 1) is upregulated by 53.43 fold under hypoxic conditions. SAGE1 is upregulated in various squamous cell carcinomas and is therefore considered a potential diagnostic or prognostic biomarker for assessing disease progression and patient treatment response. Studies have suggested that SAGE1 may promote tumor growth and development by modulating cell cycle regulation, enhancing cell proliferation, and inhibiting apoptosis. Additionally, SAGE1 has been implicated in increasing the invasiveness and migratory capacity of cancer cells, thereby facilitating distant metastasis (25,26). In terms of mechanics, the differences between 3D and 2D cultures are complex. The genes that are relevant to cell-cell adhesion and cell-ECM interaction do not exhibit collective upregulation or downregulation. For example, although CDH18, CDH11, and DSG2 are all responsible for cell-cell adhesion (27-30), their gene expression shows very different trends of upregulation, downregulation, and no change, respectively, in 3D culture (Table S3). Therefore, it is difficult to judge from the transcriptomic data whether the cell-cell adhesion changes in 3D culture. The same situation is for the genes that control cell-substrate interaction. These data suggest the major difference between 3D and 2D culture lies in the cancer stemness. Compared with the growth in adherence with the substrate, the anchorage-independent environment in 3D culture makes the genetic expression of the HEK293T cells closer to that of tumor cells.

When comparing the gene expression between hypoxia and normoxia conditions in 3D culture, the changes in the mechanics are more prominent (Table S4). Genes responsible for actin polymerization exhibit collective upregulation and those for cell-cell adhesion in general are downregulated. The expression of both INTB1 and INTB3 exhibits no significant change. All these sequencing outcomes are consistent with the PHAi, RNAi, and OE experiments reported in the main text. Besides cell mechanics, our transcriptomic data underwent Gene Ontology (GO) enrichment analysis, with Biological Process results showing significant enrichment of genes involved in extracellular matrix (ECM) organization and canonical glycolysis, which suggest the involvement of extracellular matrix remodeling and metabolism (Table S4). ECM organization refers to processes involving the assembly, maintenance, and remodeling of the extracellular matrix, a complex network of proteins and polysaccharides secreted by cells (31,32). ECM is crucial for cellular functions such as morphology, differentiation, migration, and signal transduction (33). It provides physical support and protection to cells and helps maintain tissue structural integrity (34). Components of the ECM serve as carriers of signaling molecules, facilitating intercellular communication. The structure and composition of the ECM influence cellular migration and are vital for processes such as embryonic development (35,36), wound healing (37-39), and cancer metastasis (32,40). Furthermore, ECM interacts with cell surface receptors to regulate differentiation and cell fate. Canonical glycolysis refers to the classic glycolytic pathway (41,42), in which glucose is broken down into pyruvate. Glycolysis is one of the primary mechanisms for ATP production in cells, particularly under hypoxic conditions (41,43). This pathway is critical for cell survival and proliferation, especially in rapidly proliferating cells, such as cancer cells (44,45).

Under hypoxic conditions, the upregulation of genes associated with these processes indicates that cancer cells are actively involved in tumor microenvironment remodeling (46). Cancer cells actively utilize and alter their surrounding environment to promote tumor progression, fostering an environment conducive to tumor formation (47-49). Moreover, glycolysis is a fundamental metabolic process, particularly in cancer cells, where its aberrant activation and dysregulation play a critical role in tumor initiation, progression, and metastasis. Tumor cells are often in a state of rapid proliferation, resulting in an elevated energy demand (50). Glycolysis provides a fast means of ATP production under hypoxic conditions, thus meeting the energy needs of tumor cells. Intermediates generated during glycolysis, such as 3-phosphoglyceraldehyde and phosphoenolpyruvate, can be used for the synthesis of nucleic acids, lipids, and proteins, supporting the rapid proliferation of cancer cell. Additionally, the NADH and NADPH produced through glycolysis contribute to antioxidant defense mechanisms, protecting tumor cells from oxidative stress-induced damage (51,52). In summary, under hypoxic conditions, tumor cells actively reprogram their behavior, including metabolic pathways and protein expression, to facilitate tumorigenesis and progression.

In brief, we identified enrichment in signaling pathways related to glycolysis and cell-extracellular matrix interactions. As mentioned earlier, the expression of glycolytic enzymes was significantly upregulated under hypoxic conditions, and genes related to extracellular matrix components secreted by cancer cells were also significantly upregulated. These findings are consistent with the altered energy metabolism of tumors and the remodeling of the tumor microenvironment.

Although thousands of genes are upregulated or downregulated in response to hypoxia, no existing signal pathway was identified. Some examples of classical signal pathways are summarized in Fig. S11. The MAPK (Mitogen-Activated Protein Kinase) signaling pathway plays a critical role in the initiation, progression, and metastasis of cancer (53,54). This pathway consists of three main branches: the ERK (Extracellular signal-Regulated Kinase) pathway, the JNK (c-Jun N-terminal Kinase) pathway, and the p38 MAPK pathway (55-57). These pathways regulate various biological processes such as cell proliferation, differentiation, survival, migration, and apoptosis through a series of cascade reactions (55,58). Ras is one of the key initiators in the MAPK pathway, receiving signals from cell surface receptors and transmitting them to downstream Raf kinases. Our sequencing data indicate that under hypoxic conditions, Ras expression is upregulated, along with the expression of downstream genes such as c-fos, MKP, and AP1. Additionally, we observed the downregulation or no significant change in the expression of some genes. Hypoxia-inducible factor 1 (HIF-1) is a transcription factor activated under hypoxic conditions (43,59). Its activation not only helps cells adapt to low-oxygen environments but also plays a role in various physiological and pathological processes, including tumor development. In our signal enrichment analysis, while the expression of HIF1α did not show a significant change, all downstream target genes of this pathway were significantly upregulated. This indicates that a series of downstream target genes and their associated activities are activated in response to hypoxia.

**VI. RNAi, PHAi, and OE experiments.**

The gene expression and pharmaceutical perturbation is a major approach we used to identify the effects of various molecular factors. We provide more details of such experiments in this section. The contents include the working principles of each perturbation method, the procedures and doses used in the experiments, and the raw outcome of the experiments.

Rho Activator II (CN03) was obtained from Cytoskeleton, provided as a grayish-white lyophilized solid. The active site of CN03 is based on the catalytic domain of the bacterial cytotoxic necrotizing factor (CNF) toxins (60-62). The catalytic domain is covalently attached to a proprietary cell penetrating moiety. CN03 activates Rho GTPase isoforms by deamidating glutamine-63, which is located in the Switch II region of these GTPases. This modification converts glutamine-63 to glutamate, which blocks intrinsic and GAP-stimulated GTPase activity, resulting in constitutively active Rho (61). CN03 robustly increases the level of GTP-bound RhoA within 2-4 h after addition to the culture medium. Moreover, the targeted action of this activator makes it a far more attractive tool for the study of Rho GTPase signaling than classic indirect activators (LPA) that concomitantly activate other signaling pathways (Ras, PI3K, and PLC). Each vial contains 20 µg of CN03 protein with a purity exceeding 80 %. The lyophilized protein should be stored in a desiccated state at 4 °C for no more than 6 months. For reconstitution, centrifuge at 1400 rmp for 5 min to collect the product at the bottom of the tube and resuspend each vial in 200 µl of sterile water, place on ice for 10 minutes prior to mixing, and mix by gently pipetting up and down to yield a concentration of 0.1 µg/µl. During use, the compound was diluted in a culture medium to achieve a final concentration of 2 µg/mL for cell treatment.

Y-27632 was purchased from Yeasen Biotechnology and is provided as a powder with a purity exceeding 98 %. Y-27632 is an orally bioavailable, ATP-competitive selective inhibitor of Rho-associated protein kinase (ROCK). It is a potent, cell-permeable, reversible, and selective inhibitor, with a Ki value of 140 nM for p160 ROCK (63-65). The powder should be aliquoted and stored at -20 °C dry environment for no more than 2 years. To prepare a stock solution, 10 mg of Y-27632 was dissolved in 312 µL of DMSO to create a 100 mmol/L solution, which was then diluted 10-fold with DMSO to achieve a concentration of 10 mmol/L. This solution was stored at -20 °C. For cell treatment, the solution was further diluted in culture medium to a final concentration of 30 µM.

Under hypoxia conditions, the tumor spheroid formation rate nearly doubled compared to normoxia conditions, resulting in larger volumes and a greater number of tumor spheroids. Analysis of live-cell imaging data revealed that tumor spheroids exhibited more complex dynamic behavior under hypoxia. The fusion frequency between tumor spheroids increased significantly, with the time required for two spheroids to fuse being markedly reduced (average fusion time under hypoxia was approximately 6.4 hours, compared to 21.1 hours under normoxia). Additionally, we observed that tumor spheroids underwent greater deformation under hypoxia conditions. Thus, we hypothesized that the increased fusion frequency under hypoxia is closely related to cytoskeleton properties that mainly involve dynamical-mechanical coupling. To validate the hypothesis above, we employed PHAi, RNAi, and OE experiments to examine the major components of the cytoskeleton one by one. The detailed results of all perturbation experiments are summarized in Table S5. In the next paragraph, we present an extended discussion to support the opinions in the main text.

Rho Activator II, which enhances RhoA levels, promoted actin polymerization, while Y-27632 inhibited Rho, thereby suppressing actin polymerization. Our results demonstrated that the addition of Rho Activator II under normoxia conditions significantly increased the fusion frequency of tumor spheroids (23.9 % in normoxia vs. 49.5 % Rho Activator II treatment) and notably decreased fusion time (21.1 hours in normoxia vs. 6.4 hours after treatment), leading to increased tumor formation rates. Conversely, the addition of Y-27632 under hypoxia conditions reduced tumor formation rates and fusion frequency (47.5 % in hypoxia vs. 13.4 % after Y-27632 treatment), with fusion time greatly increased (9.4 hours in hypoxia vs. 28.2 hours after treatment). This suggests that promoting actin polymerization under hypoxia conditions enhances the fusion frequency of tumor spheroids, ultimately leading to increased tumor formation rates. Tumor spheroids consist of multiple cells, and intercellular interactions may influence the overall dynamics of the spheroids. The results of RNA sequencing indicated that genes mediating intercellular interactions, such as PCDH20 and PCDH15, are downregulated under hypoxia conditions. By reducing DSG2 expression in tumor spheroids via RNA interference under normoxia conditions, we observed a slight increase in tumor formation rates (147 % vs. 207 %). In contrast, overexpression of DSG2 under hypoxia conditions resulted in decreased tumor formation rates (321 % vs. 188 %), indicating that intercellular interactions also impact tumor dynamics.

**VII. Effects of cell-ECM interaction.**

The interactions between tumors in the tumor microenvironment significantly influence tumor growth. Transcriptome sequencing results indicated that the gene expression related to the interaction between cells and the extracellular matrix did not change. We investigated the effect of the interaction between tumor spheroids and the extracellular matrix (ECM) on tumor formation rates. Under normoxia conditions, we reduced the expression levels of ITGB3 in tumor spheroids using RNA interference (RNAi), theoretically weakening the interaction between the spheroids and the ECM and allowing greater freedom for the spheroids. Experimental results indicated that the reduction of ITGB3 did not affect the tumor formation rate (normoxia: 147 % vs. normoxia with ITGB3 RNAi: 155 %) (Fig. S8A, nor did it change the fusion frequency of the tumor spheroids (normoxia: 23.9 % vs. normoxia with ITGB3 RNAi: 23.5 %) (Fig. S8B). Furthermore, under hypoxia conditions, the overexpression of ITGB3 to enhance spheroid-ECM interactions also did not impact the tumor formation rate (hypoxia: 321 % vs. hypoxia with ITGB3 overexpression: 329 %). The expression of ITGB3 exhibits little effect on the dynamics of the spheroid, like the distance before fusion (Fig. S8G) and speed (Fig. S8E), either. These numbers are summarized in Table S5. Therefore, we propose that the interactions between tumor spheroids and the ECM may not play a significant role in the growth strategies observed in our study. We also examined the three-dimensional collagen gel network in which our tumor spheroids were cultured. Our findings revealed that the network consists of stacked, multi-layered two-dimensional structures, each containing multiple pores with an average diameter of approximately 40 µm (Fig. S3). The pore sizes are larger than the initial diameter of the tumor spheroids, suggesting that the spheroids exist in a relatively free three-dimensional environment. This may explain why interactions between cells and the ECM have a limited effect during the early growth phase of tumors.

**VIII. Optimize rather than “the more the better”**

In the main text, we propose an opinion that the properties of the cytoskeletons are optimized for the proper functioning of the spheroid. Emphasizing “optimize” means there is no simple linear relationship between the spheroid dynamics and the expression of any cytoskeleton components. Instead, multiple factors are tuned and balanced into the most suitable receipt for the directional fusion. This picture distinguishes the delicate bio-adaptation of tumor cells and spheroids from the scope of non-living models. In this section, we offer more experimental evidence to support our opinion further.

In the main text, we report that actin polymerization is promoted and cell-cell adhesion is weakened in response to hypoxia (Fig. 1D-G). However, if we further enhance actin polymerization or weaken cell-cell adhesion under hypoxic conditions with PHAi and RNAi approach, the directional fusion and tumorigenesis won’t be further encouraged. In surprising contrast, the dynamics and tumorigenesis fall back to the value of normoxia (Fig. S8I-L). There is no simple linear relationship between directional fusion and spheroid stiffness either. In general, actin polymerization advances cell polarization and makes them more dynamic, and it also leads to higher stiffness. However, weakening cell-cell adhesion usually makes the spheroid softer. These two cytoskeleton factors have the opposite effect on the spheroid stiffness. The RNAi experiments on DSG2 tell the same story. Further weakening cell-cell interaction under hypoxic conditions makes the tumorigenesis behaviors similar to those under normoxic conditions (Fig. S8I-L). The fact that no certain signal pathway could be identified from the sequencing data also suggests the systematic optimization of many factors (Fig. S11). In summary, to realize the bio-function of the directional fusion, multiple factors in the cytoskeleton are balanced to the most appropriate states.

**IX. Scanning tunneling microscope experiments.**

To further investigate the differences between tumor spheroids under hypoxic and normoxic conditions, we performed scanning electron microscopy (SEM) imaging on three-dimensionally cultured tumor spheroids (66). Imaging was conducted along the z-axis at 70 nm intervals, with continuous acquisition lasting over 12 hours. More than 300 images were captured for each treatment group, yielding an overall three-dimensional reconstruction depth exceeding 20 μm. Analysis of the SEM images revealed clearly discernible mitochondrial structures (67), including cristae, at the resolution achieved. We quantified the number of mitochondria per cell and found that spheroids under normoxic conditions contained approximately 140 mitochondria per cell, whereas those under hypoxic conditions contained about 240 mitochondria per cell. This increase in mitochondrial number under hypoxia may be closely linked to cellular metabolic adaptations, supported by our live-cell imaging results indicating more dynamic behavior in hypoxic tumor spheroids.

Furthermore, we observed that hypoxic tumor spheroid cells exhibited a bimodal distribution in lysosomal content (68): approximately 35% of cells contained abundant lysosomes, while the remaining cells possessed few lysosomes. This pronounced heterogeneity suggests a form of phase separation based on lysosomal abundance within the same spheroid. The underlying cause of this considerable variability warrants further investigation. Finally, tiny nuclei were detected via SEM in hypoxic tumor spheroids, typically localized at the spheroid periphery, which is consistent with observations from confocal imaging.

**X. Chemically induced hypoxia by CoCl_2_**

Cobalt chloride (CoCl2) is one of the most commonly used models to induce chemical hypoxia. It can stabilize the expression of HIF-1α and HIF-2α in cells under normoxia conditions for relatively prolonged periods, facilitating the handling and analysis of samples within a broader time window (69). The stabilization and accumulation of HIF-1α occur because these chemical agents inhibit the activity of prolyl hydroxylases (PHDs), interfere with the hydroxylation of HIF-1α, and suppress the ubiquitin-dependent 26S proteasomal degradation pathway (69-71).

To double-check the phenomena observed under physical hypoxia in the 3-gas incubator, we conducted a chemical hypoxia experiment by adding Cobalt Chloride (CoCl2) with the following procedures. CoCl2 was purchased from Sigma (catalog number 232696). A total of 0.12984 g of CoCl2 was weighed and dissolved in 10 mL of PBS to prepare a 100 mM CoCl2 solution. The prepared solution was then filtered through a 0.22 µm filter, aliquoted into sterile centrifuge tubes, and stored at -20 °C in the dark. During the experiments, 3 µL of the prepared CoCl2 solution was added to 1 mL of culture medium before imaging. Imaging procedures were carried out as previously described.

As summarized in Fig. S11, following CoCl2 treatment, the fusion frequency of tumor spheroids increased (39.4 % with CoCl2 treatment vs. 23.9 % under normoxia), and the fusion time was shortened (12 hours with CoCl2 treatment vs. 23 hours under normoxia). The overall behaviors of the tumor spheroids are consistent under chemical and physical hypoxia. Additionally, we monitored the expression of HIF1α at various time points post-CoCl2 treatment, finding that HIF1α mRNA expression was upregulated following CoCl2 exposure (Fig. S11F). In contrast, no discernible upregulation of HIF1α occurs in the physical hypoxia samples (Fig. S4I). This inconsistency in molecular pathways implies that the classical HIF1α pathway is not the major inducing factor of the directional fusion.

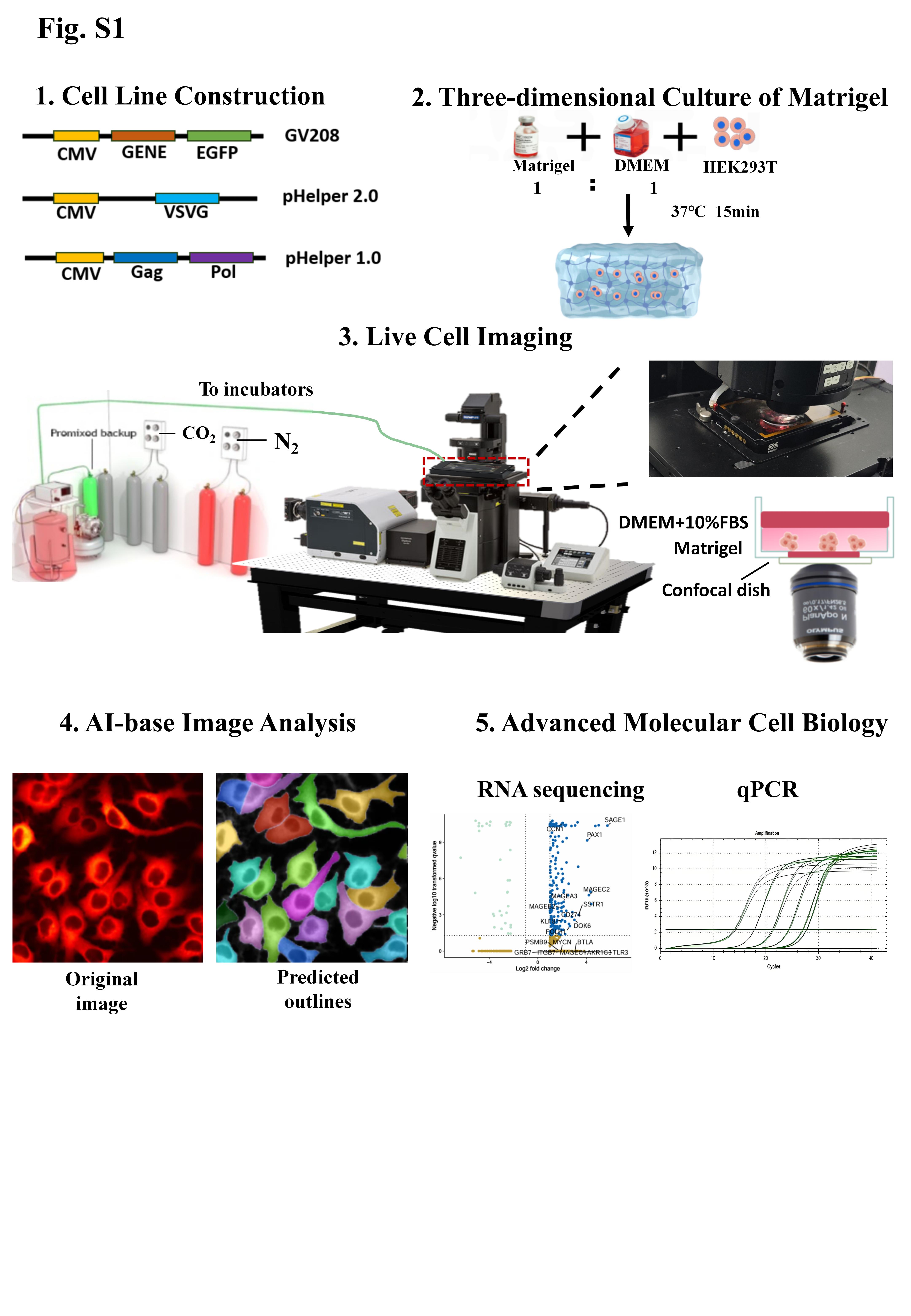

**Fig. S1. Schematics of the experimental design and workflow of this study.** In the sample preparation stage, lentivirus is used to construct cell lines expressing target proteins with fluorescent tags. The cells are seeded into a three-dimensional matrix gel network, and spheroids are allowed to form over 24 hours. Live-cell imaging is then performed on the integrated platform composed of a spinning disk confocal microscope and a three-gas incubator. The whole dynamical processes over extended periods are recorded under various oxygen concentrations with minimal phototoxicity. The obtained imaging data are processed by AI-based cell segmentation. The dynamic information of the tumor spheroid is extracted using homebuilt MATLAB codes and ImageJ plugin. Advanced molecular cell biology techniques, including transcriptomic sequencing and qPCR, are used to further elucidate the molecular mechanisms behind the dynamical phenotype.

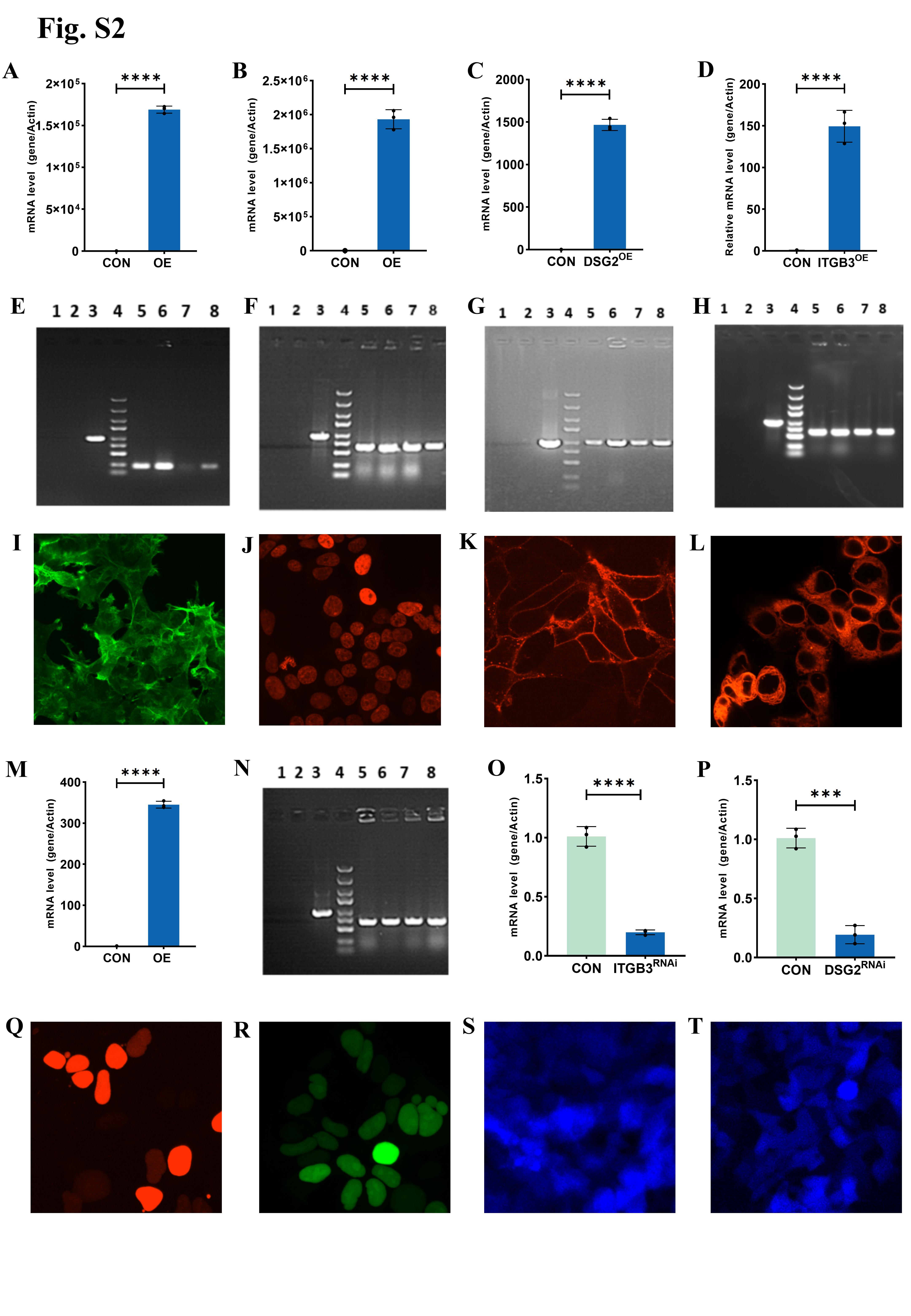

**Fig. S2. Validation of the techniques, materials, and methods used in this study.** (**A-D**), qPCR quantification of mRNA expression levels of Lifeact labeled with green fluorescence to indicate F-actin in HEK293T-Actin-H2B (**A**), H2B in HEK293T-Actin-H2B (**B**), DSG2 in HEK293T-DSG2OE (**C**) and ITGB3 in HEK293T-ITGB3OE (**D**) cell lines. (**E-H**), Electrophoresis results of PCR-amplified target genes inserted into the expression vector. (**E**) corresponds to Lifeact, (**F**) to H2B, (**G**) to DSG2, and (**H**) to ITGB3. Lane 1: negative control (ddH₂O); Lane 2: negative control (empty vector); Lane 3: positive control (GAPDH); Lane 4: DNA ladder, with bands from top to bottom corresponding to 5 kb, 3 kb, 2 kb, 1.5 kb, 1 kb, 750 bp, 500 bp, 250 bp, and 100 bp; Lanes 5–8: transformants. (**I-L**), Representative confocal images of the established cell lines expressing green fluorescence for F-actin (**I**), red fluorescence for H2B (**J**), red fluorescence for DSG2 (K), and red fluorescence for ITGB3 (**L**). (**M**), qPCR quantification of mRNA expression levels of Geminin in the fluorescently labeled cell line for Cdt1 and Geminin. (**N**), Agarose gel electrophoresis analysis of PCR products amplifying the target gene Geminin after its insertion into the expression vector. Lane 1: negative control (ddH₂O); Lane 2: negative control (empty vector); Lane 3: positive control (GAPDH); Lane 4: DNA ladder with bands (from top to bottom): 5 kb, 3 kb, 2 kb, 1.5 kb, 1 kb, 750 bp, 500 bp, 250 bp, and 100 bp; Lanes 5–8: transformants. (**O, P**), qPCR quantification of mRNA expression levels of ITGB3 in the ITGB3 RNAi cell line (**O**) and DSG2 in the DSG2 RNAi cell line (**P**). (**Q, R**), Cell cycle profiling of the established cell line: Cdt1 (red, G1 phase) and Geminin (green, S/G2/M phases). (**S, T**), RNAi validation: blue fluorescence indicates shRNA transfection for ITGB3 (**S**) and DSG2 (**T**). For all statistical plots, unpaired Student’s t-test (n > 3 experiments, data are mean ± SEM) was conducted for all data with **** P < 0.0001, *** P < 0.001, ** P < 0.1, * P <0.5, ns P > 0.5. The error bars are the standard error of the mean of the data.

**Fig. S3. The workflow comprised a 3D culture system, robust cell segmentation, and stringent RNA-seq quality control.** (**A, B**), Three-dimensional network structure of the matrix gel under bright field (**A**) and confocal illumination (**B**). The scale bar is 20 µm. (**C**), Confocal microscopy images of two-dimensional cultured cells, with red fluorescence labeling H2B and green fluorescence labeling F-actin. (**D**) The data from (**C**) were imported into CellPose 2.0 and after adjusting the relevant parameters, the cell segmentation was conducted. The shaded area and the dashed line are the cell body and cell boundary detected by CellPose, respectively. (**E**), 3D reconstruction of the tumor spheroid composed of HEK293T-Actin-H2B cells from the z-stack of confocal microscopy images. The reconstruction is done by CellSens Dimension Desktop 4.2.1, with red fluorescence labeling H2B and green fluorescence labeling F-actin. (**F**), The live-cell imaging data of 3D-cultured tumor spheroids under different experimental conditions were subjected to 3D segmentation via Cellpose, followed by the analysis of their 3D motion trajectories. (**G**), Distribution of bases after filtering the transcriptome data from 24-hour normoxia culture. The show the base positions within the reads on the x-axis and the proportion of each base type on the y-axis; different colors represent different base types. Under normal conditions, samples exhibit separation of AT and CG bases. The observed jitter in the first few base pairs is attributed to the instability of random primers and enzyme-substrate binding during the sequencing reaction, representing normal variability inherent to the sequencing process.
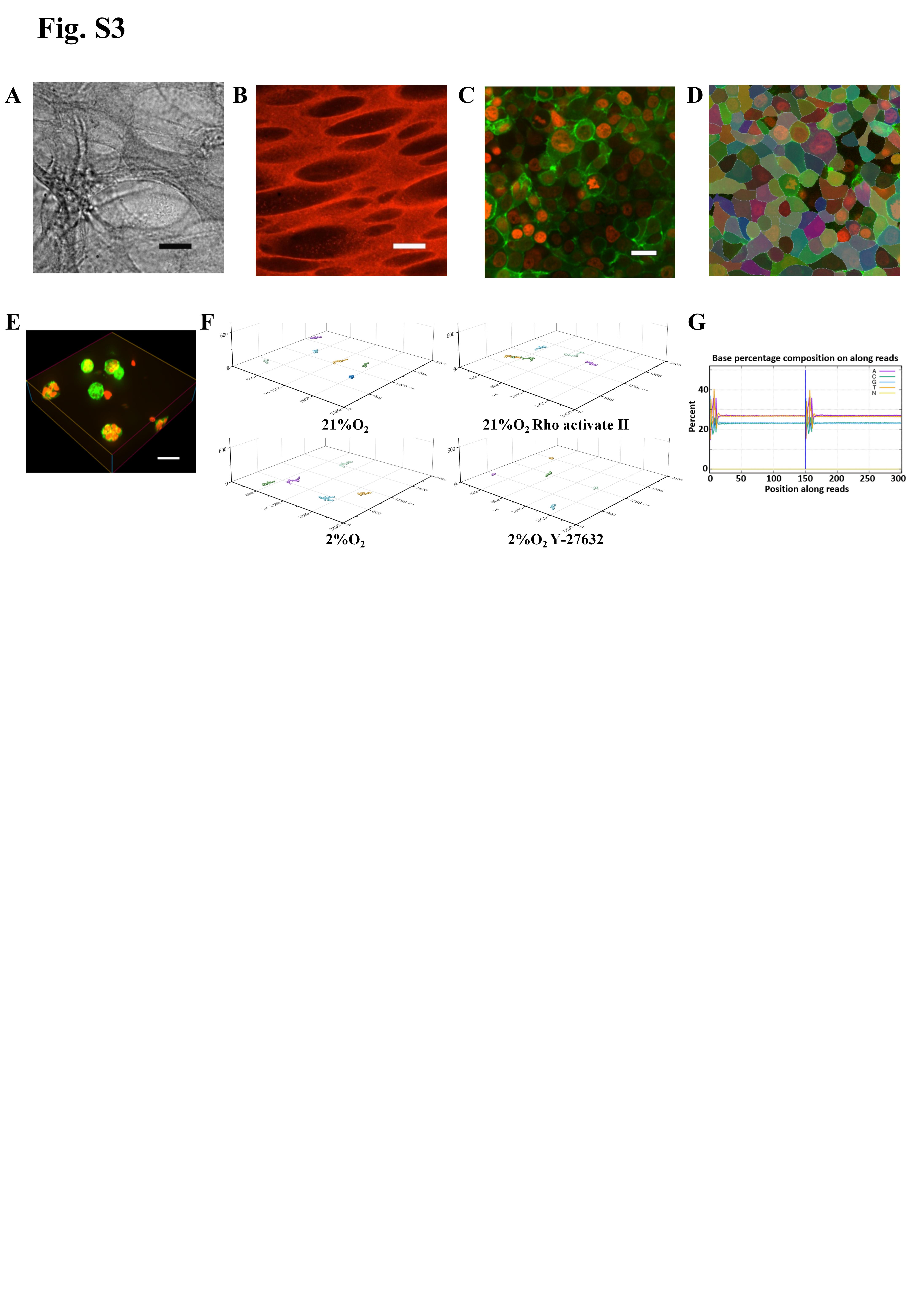

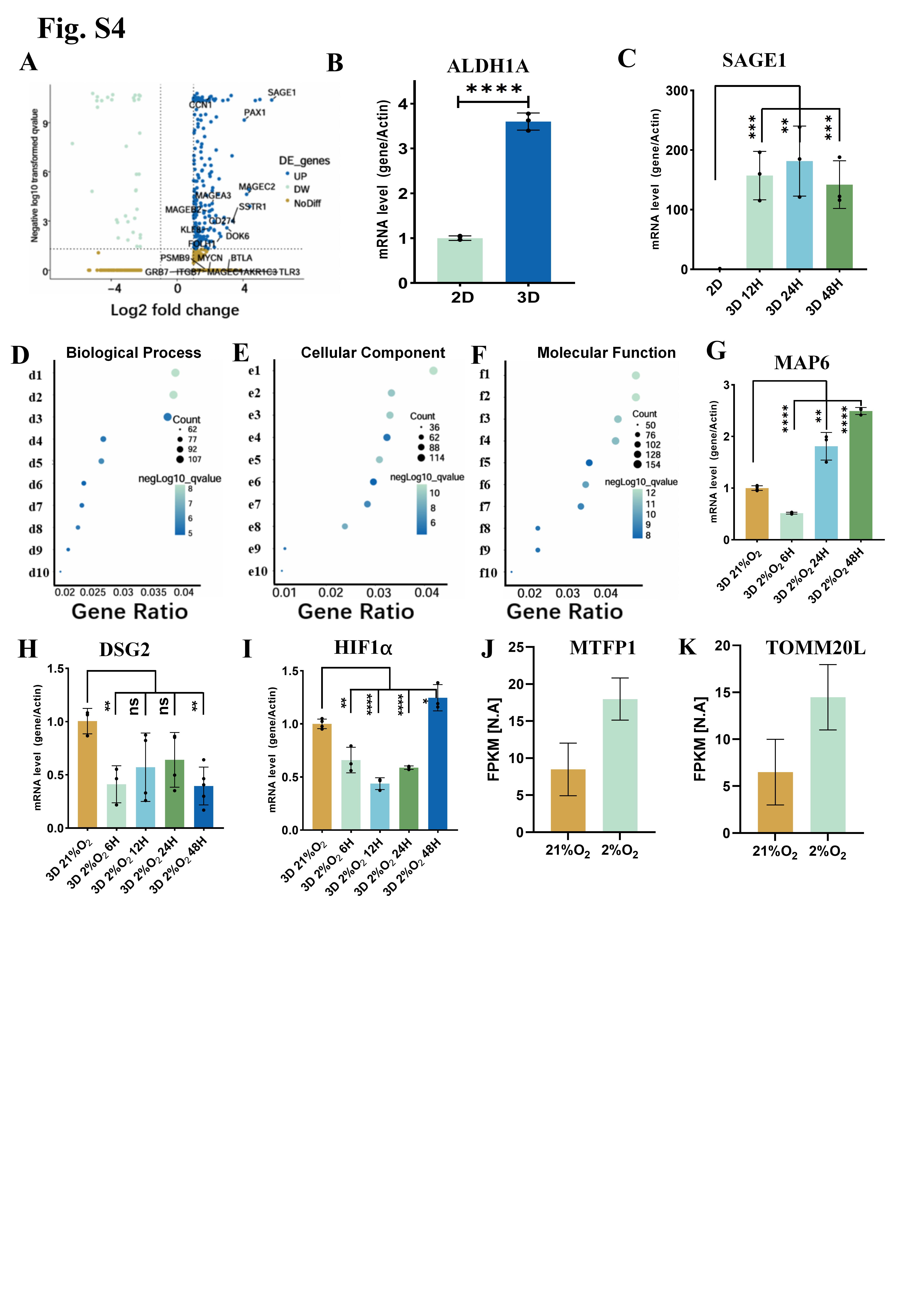

**Fig. S4. Molecular analysis on relevant bio-chemical factors.** (**A**), Volcano plots of differentially expressed genes under 3D and 2D culture. Cancer-related genes are indicated by black arrows. (**B, C**), mRNA expression levels of ALDH1A (**B**) and SAGE1(**C**) in two-dimensional and three-dimensional cultured cells. The different time points in (**C**) refer to the time after the cells are subjected to hypoxia. (**D-F**), KEGG enrichment analysis of differentially expressed genes on biological process (**D**), cellular component (**E**), and molecular function (**F**). The bubble plot illustrates the top ten representative results. The y-axis is described as follows. d1:extracellular structure organization, d2:extracellular matrix organization,, d3: positive regulation of cell adhesion, d4: cell−cell adhesion via plasma−membrane adhesion molecules, d5: G protein−coupled receptor signaling pathway, coupled to cyclic nucleotide second messenger, d6: potassium ion transport, d7: adenylate cyclase−modulating G protein−coupled receptor signaling pathway, d8: potassium ion transmembrane transport, d9: multicellular organismal signaling, d10:regulation of blood pressure; e1: collagen−containing extracellular matrix, e2: transporter complex, e3: transmembrane transporter complex, e4: synaptic membrane, e5: ion channel complex, e6: apical plasma membrane, e7: endoplasmic reticulum lumen, e8: cation channel complex, e9: potassium channel complex, e10: voltage−gated potassium channel complex; f1: passive transmembrane transporter activity, f2: channel activity, f3: ion channel activity, f4: metal ion transmembrane transporter activity, f5: monovalent inorganic cation transmembrane transporter activity, f6: gated channel activity, f7: cation channel activity, f8: voltage−gated ion channel activity, f9: voltage−gated channel activity, f10: potassium channel activity The size of each bubble represents the number of genes involved, the color indicates the significance level (p-value). Data are from three independent experiments. (**G-I**), mRNA expression levels of, MAP6 (**G**), DSG2 (**H**), and HIF1α (**I**) in three-dimensional cultured cells. The different time points in the plots refer to the time after the cells are subjected to hypoxia. (**J, K**), Quantitative analysis of the normalized gene expression levels of the mitochondrial membrane protein MTFP1 (**J**) and the mitochondrial transport complex subunit TOMM20L (**K**) under normoxic and hypoxic conditions from transcriptome sequencing data. For all statistical plots, unpaired Student’s t-test (n > 3 experiments, data are mean ± SEM) was conducted for all data with **** P < 0.0001, *** P < 0.001, ** P < 0.1, * P <0.5, ns P > 0.5. The error bars are the standard error of the mean of the data.

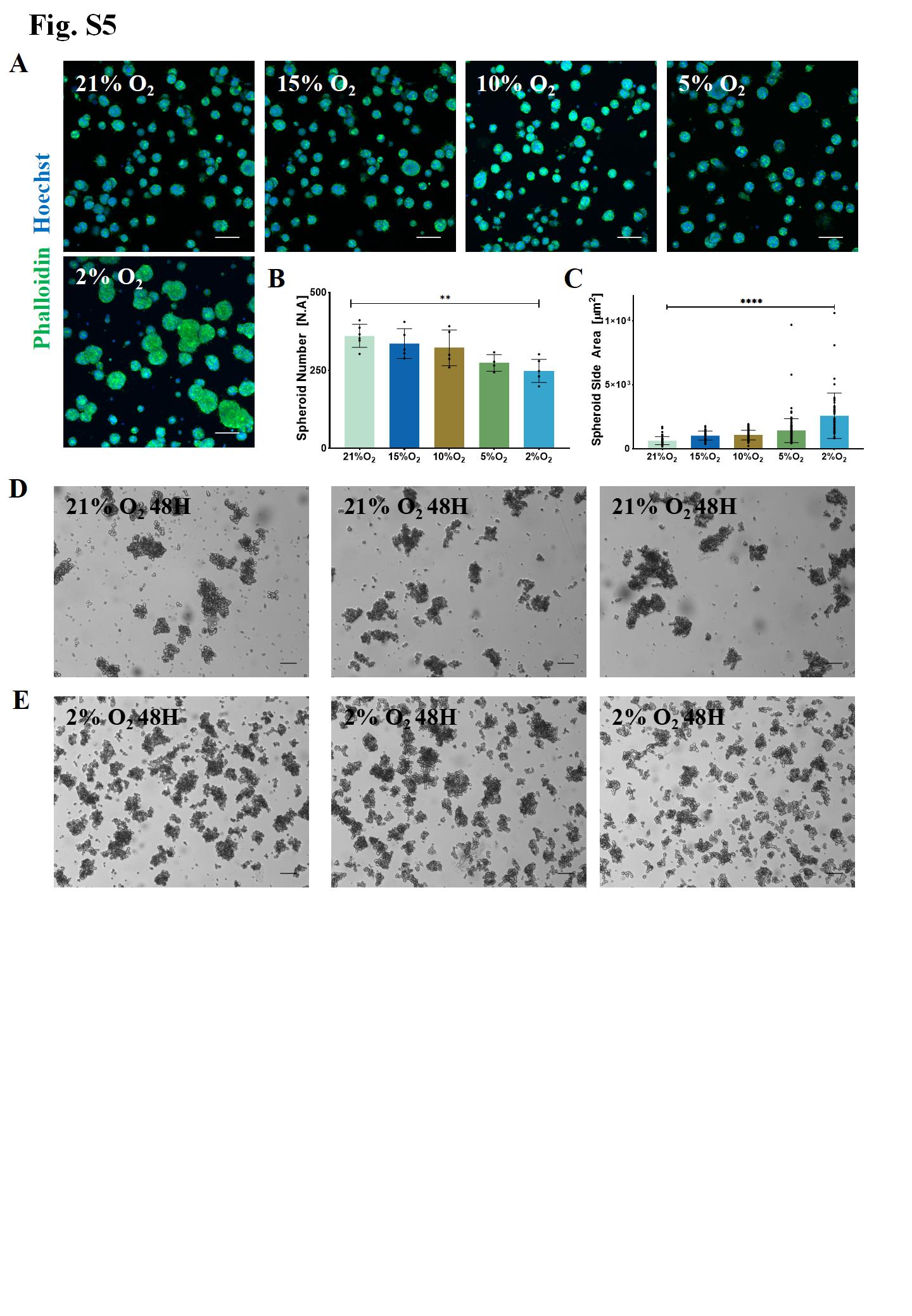

**Fig. Tumor spheroid growth of HEK293T under three-dimensional culture with different oxygen concentrations and tumor spheroid growth of NCI-H69 under 2% oxygen culture condition.** (**A**), Confocal micrographs of HEK293T cells cultured in 3D Matrigel under different oxygen concentrations for 48 h. Images are representative of more than three independent experiments. The green channel is Phalloidin which labels F-actin, and the blue channel is Hoechst which labels nuclei. The scale bar is 20 µm. (**B**), Quantification of tumor spheroid numbers following 48 h of culture under different oxygen conditions. (**C**), Average maximum cross-sectional area of tumor spheroids after 48 hours treatment under different oxygen concentrations. (**D**), Representative brightfield microscopic images of NCI-H69 cells after 48 h of culture under 21% oxygen. The scale bar is 200 µm. (**E**), Representative brightfield microscopic images of NCI-H69 cells after 48 h of culture under 2% oxygen. The scale bar is 200 µm.

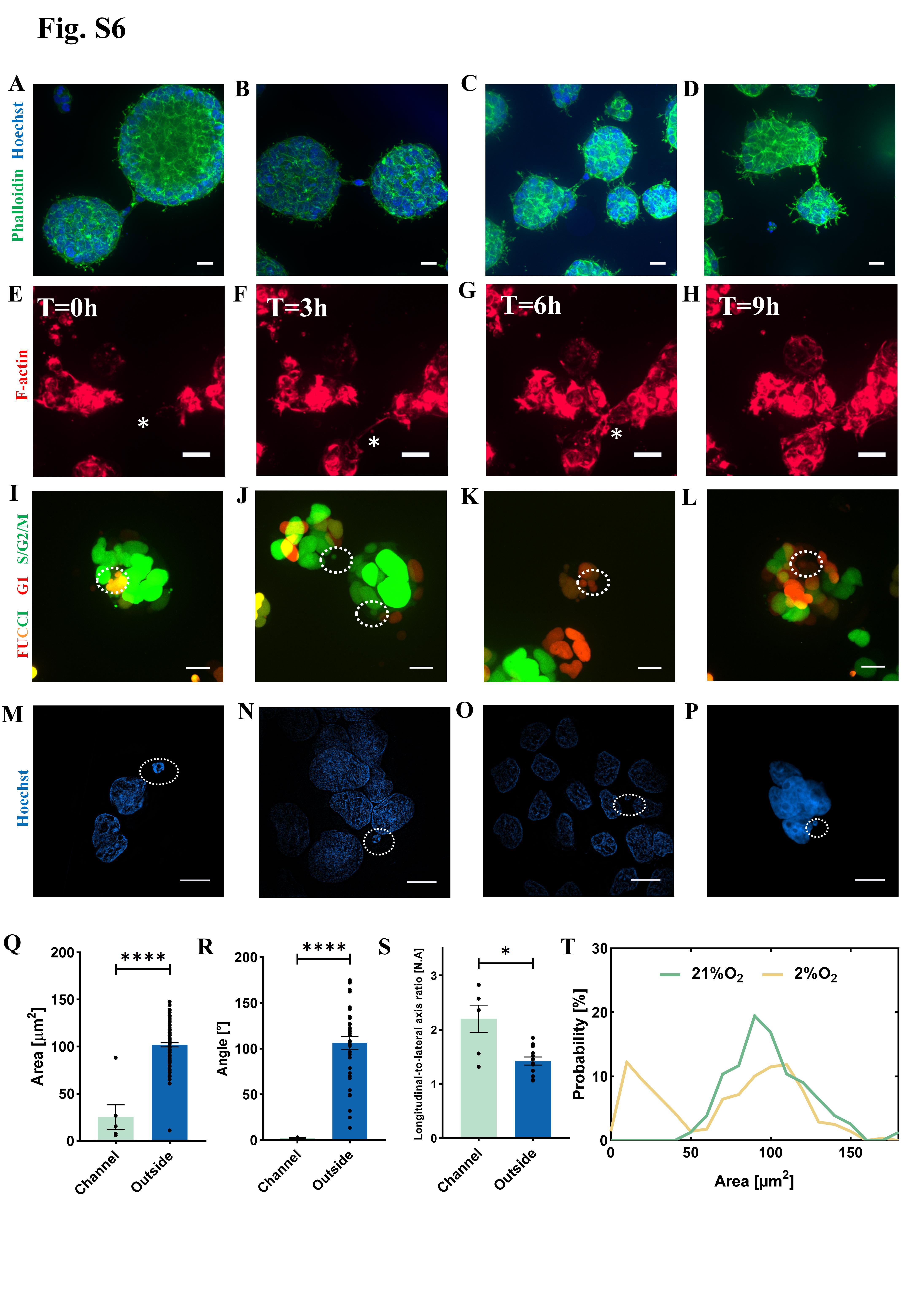

**Fig. S6. Additional analysis on the outreaching F-actin and tiny cells.** (**A-D**), Representative confocal images out of three independent experiments demonstrating the channels formed by outreaching F-actin between tumor spheroids under hypoxia conditions. The green channel is Phalloidin which labels F-actin, and the blue channel is Hoechst which labels nuclei. The scale bar is 20 µm. (**E-H**), Representative time-lapse confocal images showing the formation of channels between two tumor spheroids through F-actin. Red fluorescence represents F-actin labeled with SiR-actin. The scale bar represents 20 µm. (**I-L**), Representative confocal images out of two independent experiments demonstrating that the nuclei of the tiny cells locate between two fusing tumor spheroids under hypoxia conditions. The white dotted circle indicate the position of the tiny cells. The cells are labeled with FUCCI, in which the color indicates different stages of the cell cycle. - Red marks hCdt1, indicating cells are in the G1 phase, and green marks hGem, indicating cells are in the S/G2/M phases. The scale bar is 10 µm. (**M-P**), Representative structured illumination microscopy (SIM) images of tumor spheroids out of three independent immunostaining experiments proving that tiny cells contain chromosome materials. The cell nuclei are labeled with Hoechst. The white dotted circle indicates the tiny cells. The scale bar is 10 µm. (**Q**), Average area of nuclei inside and outside the channels formed by F-actin during tumor spheroid fusion. (**R**), The angle between the long axis of cell nuclei and the direction of the channel itself. (**S**), Ratio of the long axis length to the short axis length of cell nuclei inside and outside the F-actin channels. (**T**), Probability distribution of nucleus sizes calculated from the immunostaining data under hypoxia and normoxia conditions in 3D control. For all statistical plots, unpaired Student’s t-test (n = 3 experiments, data are mean ± SEM) was conducted for all data with **** P < 0.0001, *** P < 0.001, ** P < 0.1, * P <0.5, ns P > 0.5. The error bars are the standard error of the mean of the data.

**Fig. S7. Laser ablation experiments on the outreaching F-actin and tiny cells.** (**A-C**), Representative confocal images showing the effects of laser ablation on micronuclei after culturing for 42 hours under normoxia conditions (**A**) or for 42 hours under 2% oxygen conditions (**B**), laser ablation followed by continued hypoxia culture for an additional 24 hours (**C**). The green fluorescence indicates F-actin labeled with phalloidin, while the blue fluorescence marks the nuclei with Hoechst. The scale bar is 100 µm. (**D**), Tumor spheroid sizes under different culture conditions. (**E-G**), Representative confocal images depicting the channels formed between two tumor spheroids after laser ablation following 42 hours of culture under normoxia conditions (**E**) or 42 hours under 2% oxygen conditions (**F**), and laser ablation then continued hypoxia culture for an additional 42 hours (**G**). The green fluorescence indicates F-actin labeled with phalloidin, while the blue fluorescence marks the nuclei with Hoechst. The scale bar is 100 µm. (**H**), Tumor spheroid sizes under different culture conditions. For all statistical plots, unpaired Student’s t-test (n = 3 experiments, data are mean ± SEM) was conducted for all data with **** P < 0.0001, *** P <0.001, ** P < 0.1, * P <0.5, ns P > 0.5. The error bars are the standard error of the mean of the data.
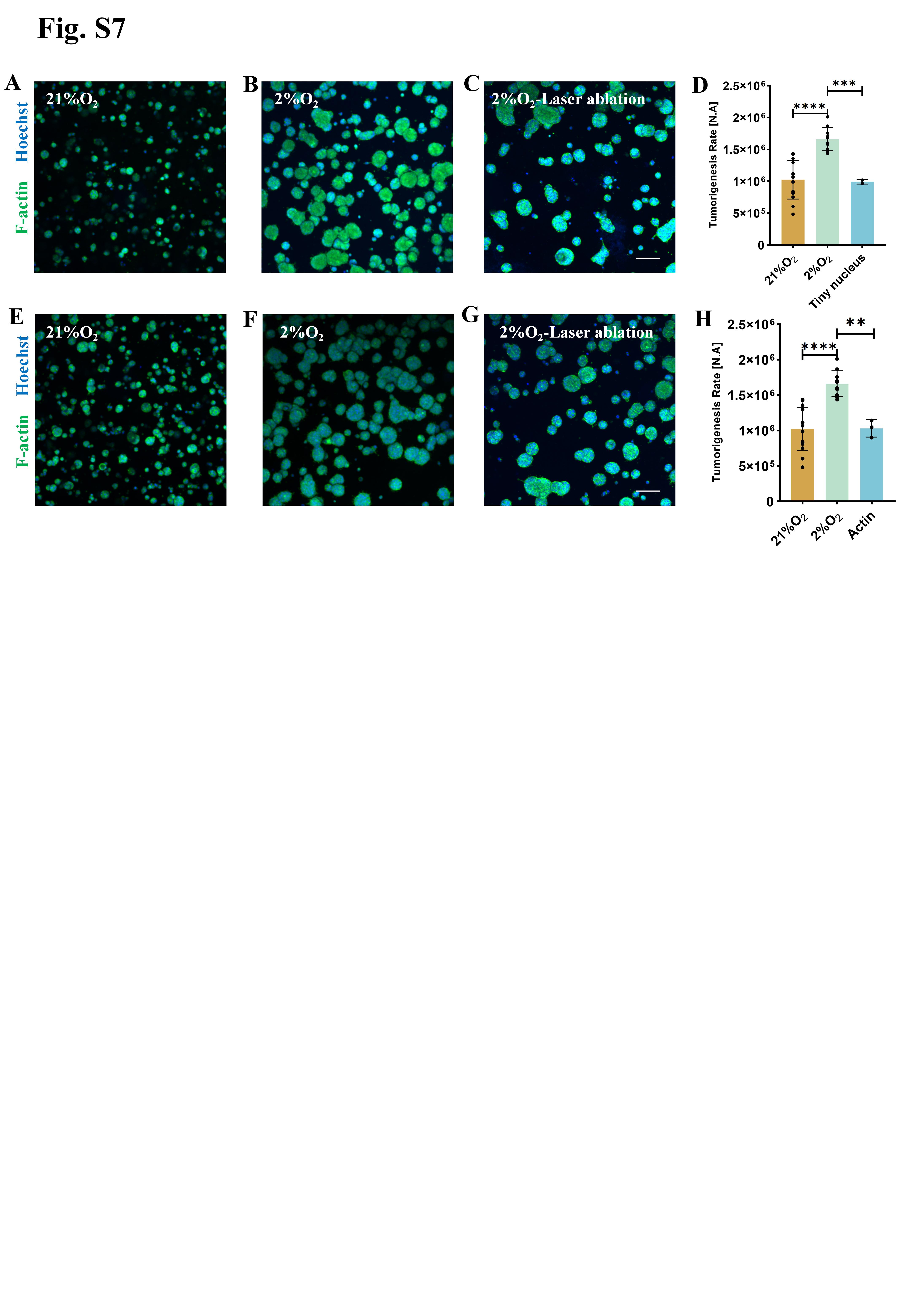

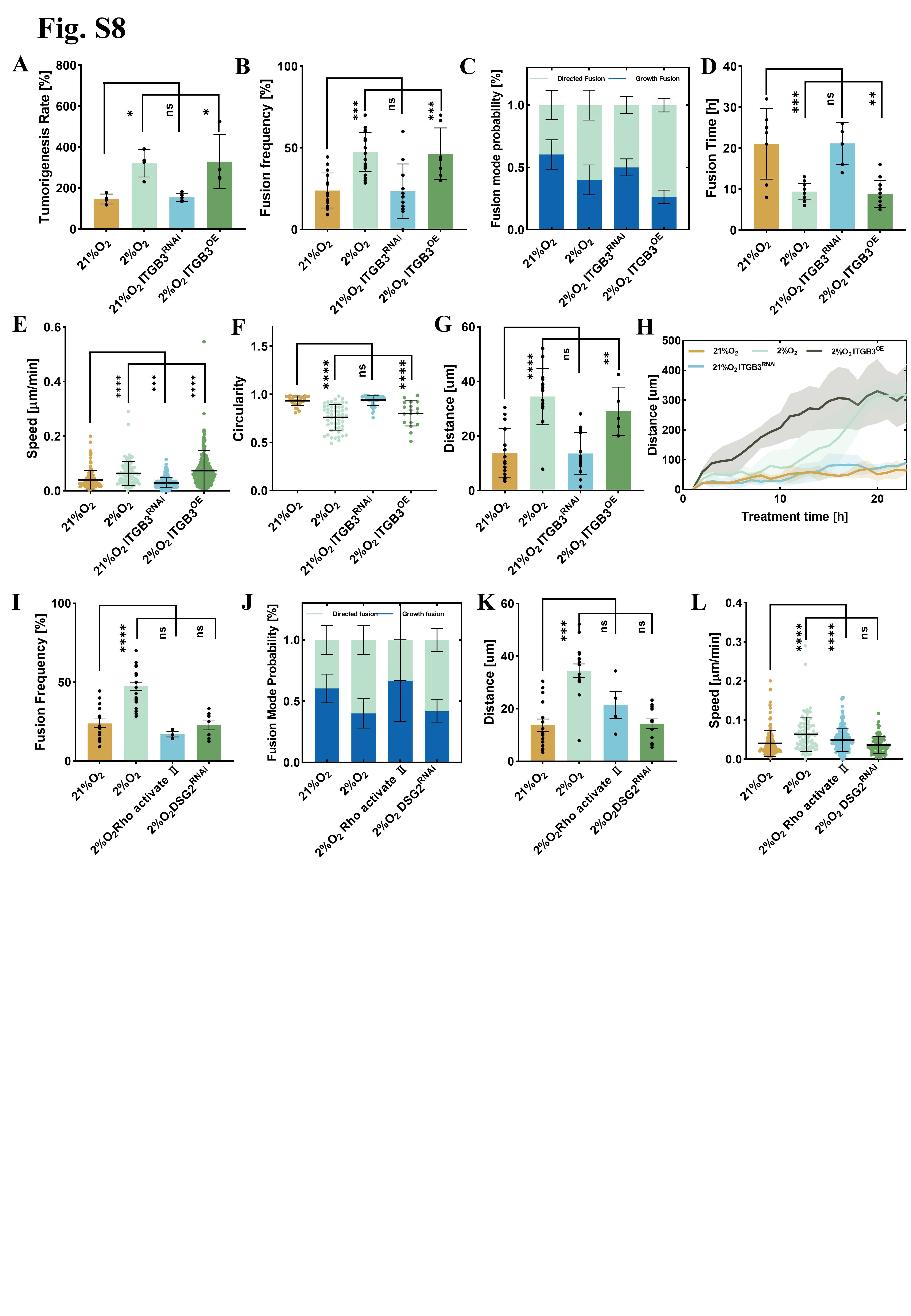

**Fig. S8. The Impact of ITGB3, DSG2, and the Regulation of Actin Assembly on Tumor Development.** (**A**), Tumorigenesis rate of HEK293T cells under different conditions of normoxia, hypoxia, ITGB3-OE, and ITGB3-RNAi treatments. (**B**), Fusion frequency of tumor spheroids under different culture conditions. (**C**), Proportion of fusion and growth modes in the fusion under different culture conditions of normoxia, hypoxia, ITGB3-OE, and ITGB3-RNAi treatments. (**D**), Time required for the directional fusion under different culture conditions of normoxia, hypoxia, ITGB3-OE, and ITGB3-RNAi treatments. (**E**), Speed in 3-dimension of tumor spheroids under different culture conditions of normoxia, hypoxia, ITGB3-OE, and ITGB3-RNAi treatments. (**F**), Circularity of the tumor spheroid during directional fusion under different culture conditions. (**G**), The distance between two tumor spheroids before directional fusion occurs under various conditions of normoxia, hypoxia, ITGB3-OE, and ITGB3-RNAi treatments. (**H**), The distance between the non-fused tumor spheroids and their initial positions over time under various conditions of normoxia, hypoxia, ITGB3-OE, and ITGB3-RNAi treatments. (**I**), Fusion frequency of tumor spheroids under different culture conditions of normoxia, hypoxia, PHAi, and RNAi treatments. (**J**), Proportion of directed and growth modes in the fusion under different culture conditions of normoxia, hypoxia, PHAi, and RNAi treatments. (**K**), The distance between two tumor spheroids before directional fusion occurs under various conditions of normoxia, hypoxia, PHAi, and RNAi treatment. (**L**), Speed in 3-dimension of tumor spheroids under different culture conditions of normoxia, hypoxia, PHAi, and RNAi treatment. For all statistical plots, an unpaired Student’s t-test (n > 3 experiments, data are mean ± SEM) was conducted for all data with **** P < 0.0001, *** P < 0.001, ** P < 0.1, * P <0.5, ns P > 0.5. The error bars are the standard error of the mean of the data.

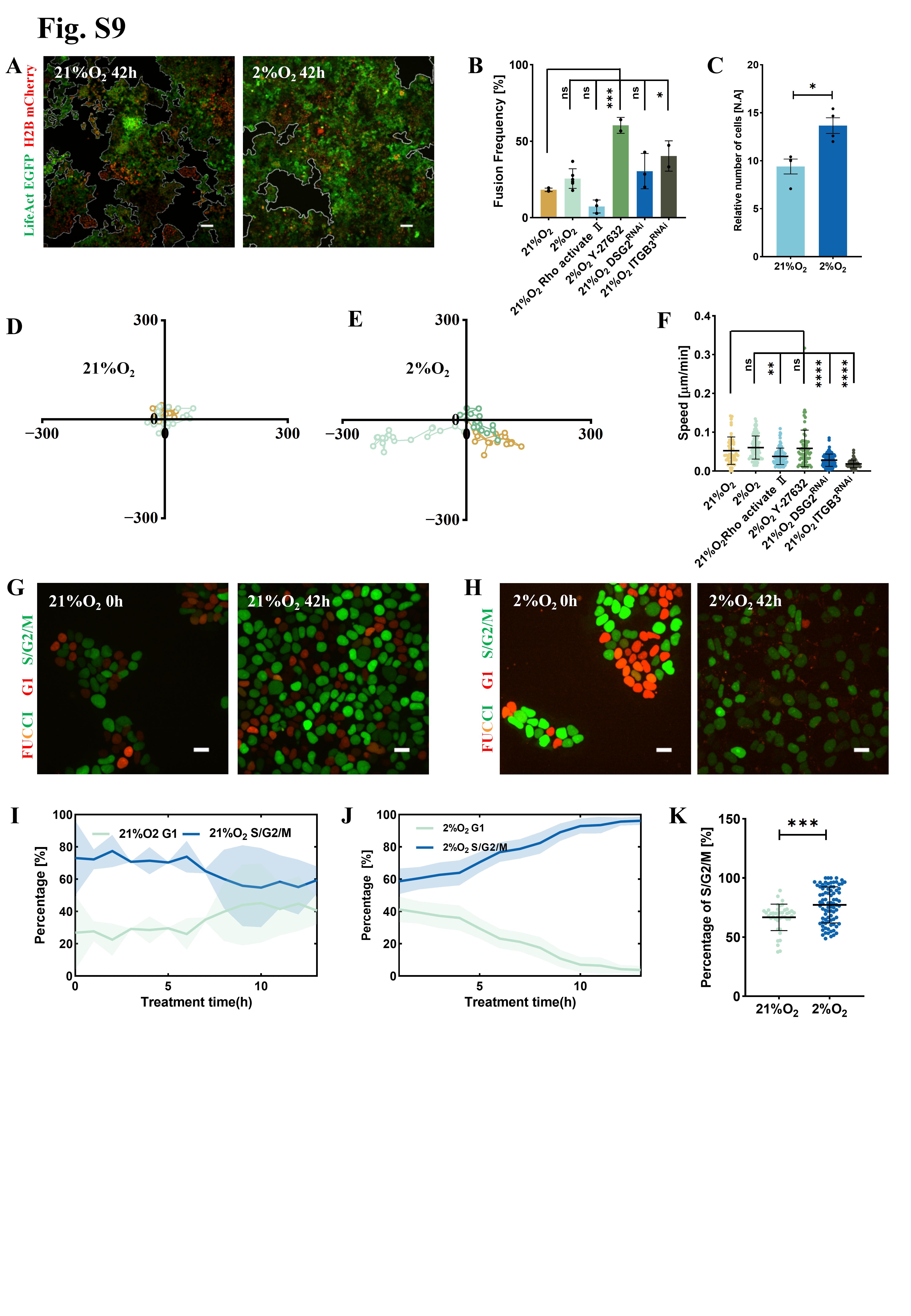

**Fig. S9. Comparison between the cell behaviors in 3D and 2D cultures under hypoxia.** (**A**), Confocal images of HEK293T cells cultured under normoxia conditions (left) and hypoxia conditions (right) for 42 hours. The white lines indicate the borders of the cell colonies. The green and red channels are LifeAct and H2B, respectively. The scale bar is 100 µm. The images are representative ones from more than three independent experiments. (**B**), Frequency of cell cluster fusion within ten hours under various culture conditions. (**C**), Relative cell number changes 42 hours after seeding. (**D, E**), Typical trajectories of HEK293T cell clusters under normoxia (**D**) and hypoxia (**E**) conditions in two-dimensional culture. (**F**), Speed of HEK293T cells under different culture conditions in 2D. (**G, H**), Confocal images of HEK293T-FUCCI cells from 3 independent experiments under nomoxia (**G**) and hypoxia (**H**) conditions in 2D culture. The left and right panels are images taken at 0 hours and 42 hours, respectively, after cell seeding. The green and red channels are hGem and hCdt1, respectively. The scale bar is 20 µm. (**I, J**), Proportion of cells in G1 phase and S/G2/M phases under normoxia (**I**) or hypoxia (**J**) conditions as a function of time. (**K**), Proportion of HEK293T cells in the S/G2/M phases under hypoxia and normoxia conditions in 2D culture. For all statistical plots, unpaired Student’s t-test (n > 3 experiments, data are mean ± SEM) was conducted for all data with **** P < 0.0001, *** P < 0.001, ** P < 0.1, * P <0.5, ns P > 0.5. The error bars are the standard error of the mean of the data.

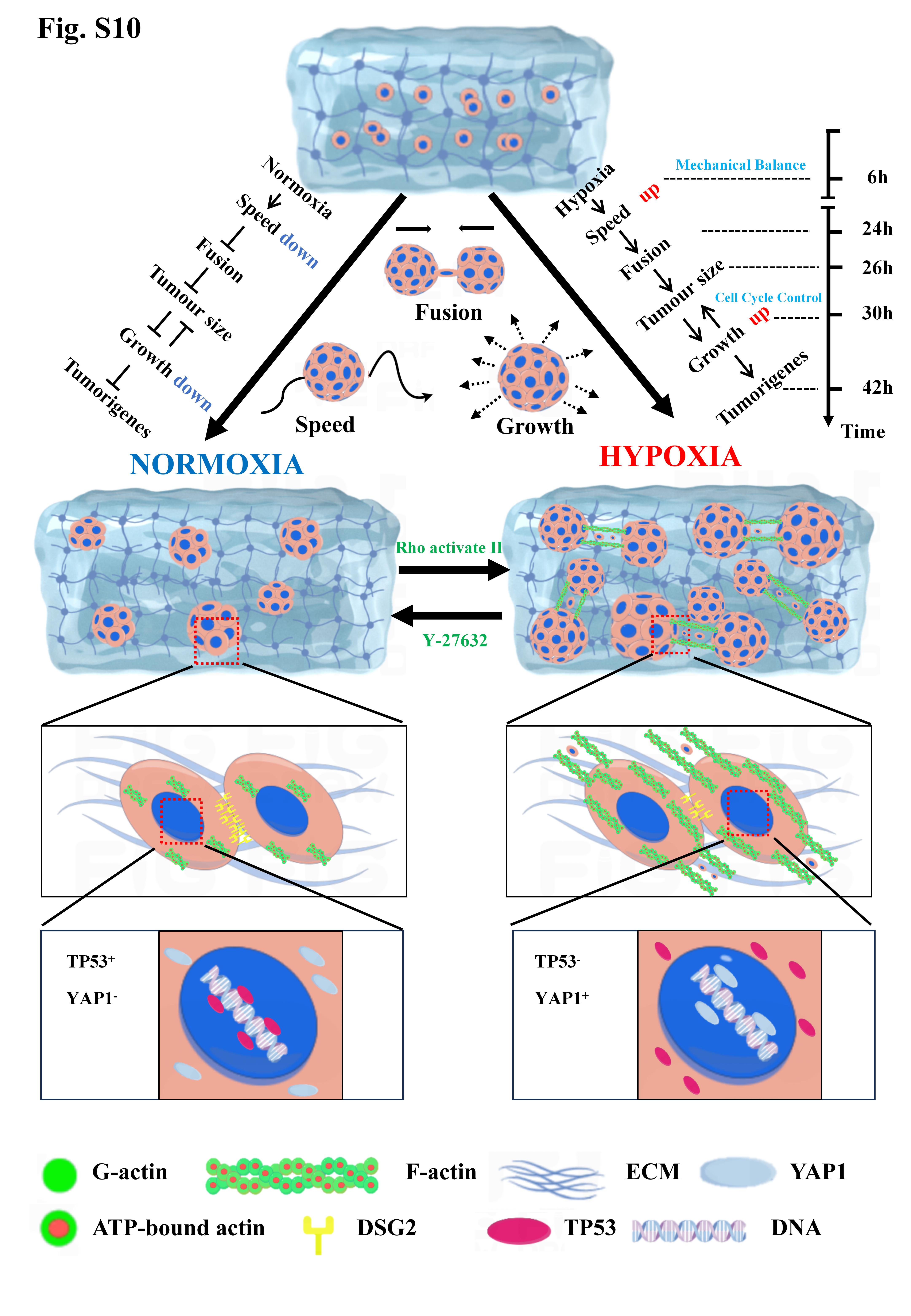

**Fig. S10. Schematic summary of early tumorigenesis under normoxia and hypoxia conditions.** The overall tumorigenesis is significantly promoted by hypoxia treatment due to the altered dynamical phenotype of the tumor spheroids, including the major contribution from directional fusion and minor contribution from enhanced growth. At the cellular level, the directional fusion is directed by the outstretched F-actin and tiny cells that bridge the gaps between the fusion spheroids. At the molecular level, cytoskeleton proteins and the nucleus are optimized for the structural basis for the dynamics and mechanics of the tumor spheroids, leading to their proper biological function in adaptation to hypoxia.

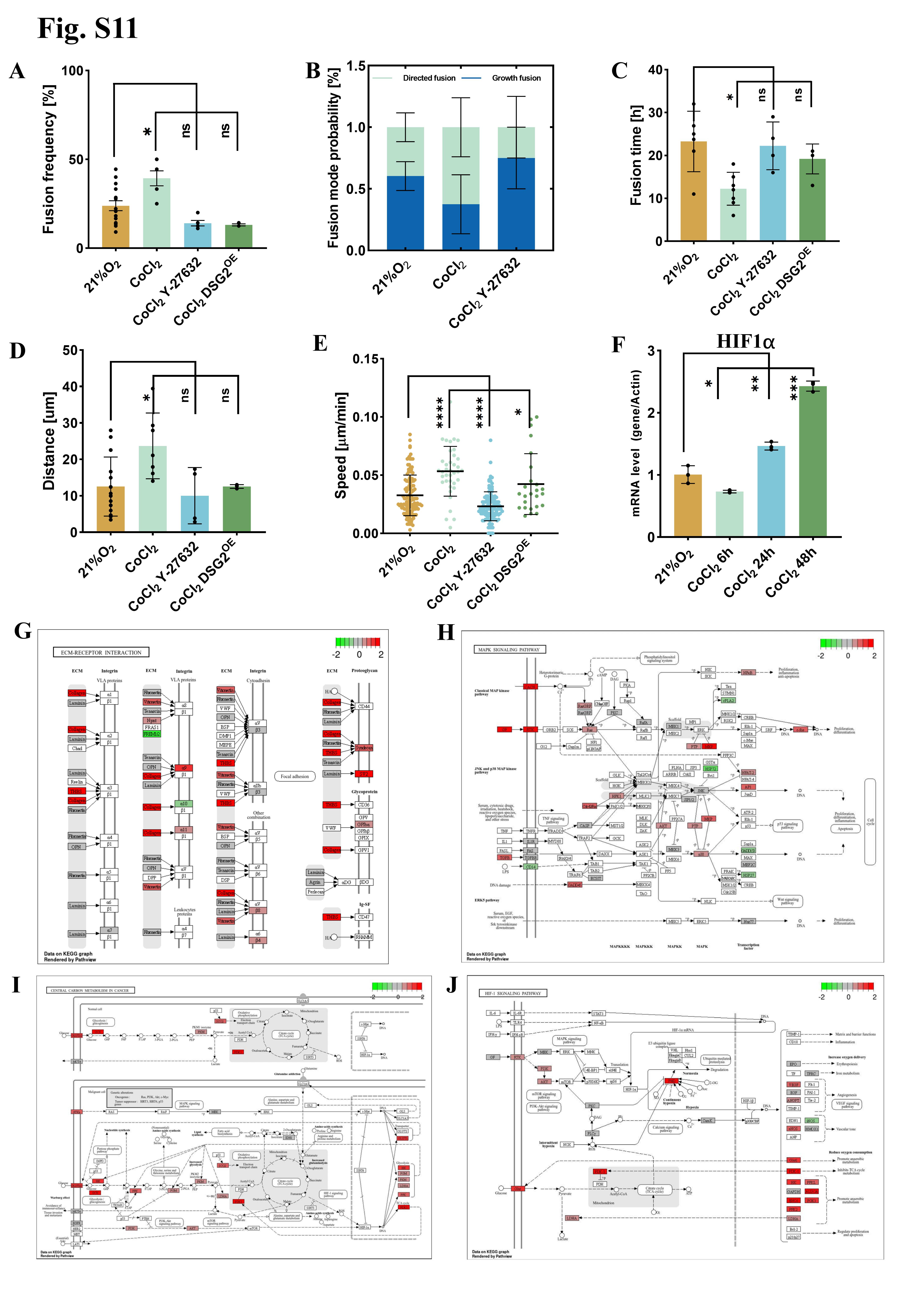

**Fig. S11. Effects of Cobalt Chloride-induced chemical hypoxia on tumors and pathway enrichment analysis of transcriptome sequencing data under hypoxic conditions.** (**A**), Fusion frequency of tumor spheroids under different culture conditions. (**B**), Proportion of directional and growth modes in the fusion under different culture conditions of normoxia, CoCl2, PHAi treatments. (**C**), Time required for the directional fusion under different culture conditions of normoxia, CoCl2, PHAi, and DSG2-OE treatments. (**D**), The distance between two tumor spheroids before directional fusion occurs under various conditions of normoxia, CoCl2, PHAi, and DSG2-OE treatments. (**E**), Speed in 3-dimension of tumor spheroids under different culture conditions of normoxia, CoCl2, PHAi, and DSG2-OE treatments. (**F**), mRNA expression levels of HIF1α in cells subjected to CoCl2-induced chemical hypoxia at different time points. (**G**), Enrichment of the cancer-related MAPK signaling pathway. (**H**), Enrichment of hypoxia-inducible factor-related signaling pathways. (**I**), Enrichment of carbon metabolism signaling pathways in cancer cells that support the energy demand of rapid proliferation. (**J**), Enrichment of adhesion molecule-related signaling pathways. For all statistical plots, unpaired Student’s t-test (n > 3 experiments, data are mean ± SEM) was conducted for all data with **** P < 0.0001, *** P < 0.001, ** P < 0.1, * P <0.5, ns P > 0.5. The error bars are the standard error of the mean of the data.

**Fig. S12. Representative scanning electron microscopy (SEM) images of cancer spheroids under normoxic and hypoxic conditions in three-dimensional (3D) culture after 42 hours, showing different Z-axis planes.** (**A-D**), SEM images of the same cancer spheroid cultured under 21% oxygen for 42 hours, captured at different depths. The scale bar is 5 µm. (**E-H**), SEM images of the same cancer spheroid cultured under 2% oxygen for 42 hours, captured at different depths. The scale bar is 5 µm.

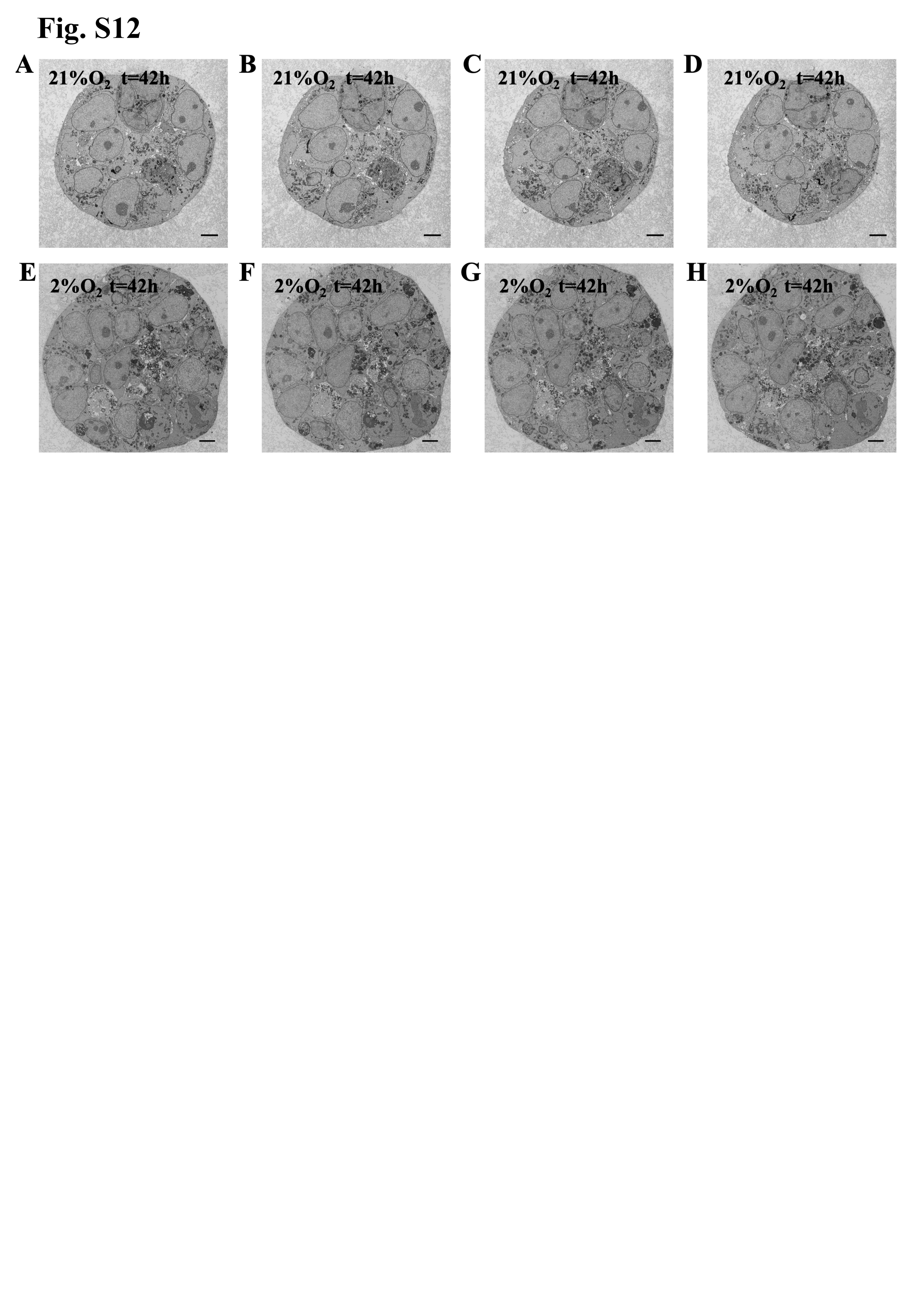

**Table S1.**

List of primers required for qPCR validation of relevant gene mRNA expression levels

| **Gene** | **primer** |
| --- | --- |
| actin-F | CATGTACGTTGCTATCCAGGC |
| actin-R | CTCCTTAATGTCACGCACGAT |
| GMNN-F | GCCCTGGGGTTATTGTCCC |
| GMNN-R | AGCGCCTTTCTCCGTTTTTCT |
| ALDH1A1-F | GCACGCCAGACTTACCTGTC |
| ALDH1A1-R | CCTCCTCAGTTGCAGGATTAAAG |
| SAGE1-F | ACTTCAAACGAGTCAACCAACT |
| SAGE1-R | TCTAACCACGAGGACATACTCTT |
| MAP6-F | TTGATCGCAGAAGAATACGCAG |
| MAP6-R | GCCTTCCTCGTGGGCTTAT |
| DSG2-F | TGCTGCTTCTCCTGATCTGC |
| DSG2-R | CAGATCCTCTCCCTCCCGAA |
| HIF1α-F | GAACGTCGAAAAGAAAAGTCTCG |
| HIF1α-R | CCTTATCAAGATGCGAACTCACA |
| LifeAct-F | TGCTTCAGCCGCTACCC |
| LifeAct-R | AGTTCACCTTGATGCCGTTC |
| H2B-F | CACTACGACGCTGAGGTCAAG |
| H2B-R | TGGTGTAGTCCTCGTTGTGGG |
| ITGB3-F | CAGGCCCAAGCTGAACCTAA |
| ITGB3-R | GAGAAGTCGTCACACTCGCA |

**Table S2.**

Summary of all cell lines used in this study

| **ID** | **Name** | **Description** | **Excitation wavelength [nm]** |
| --- | --- | --- | --- |
| 1 | HEK293T LifeAct | Visualize F-actin | 488 |
| 2 | HEK293T H2B | Visual nucleus | 561 |
| 3 | HEK293T FUCCI | Visualization and analysis of cell division cycle | 488/561 |
| 4 | HEK293T DSG2OE | DSG2 was overexpressed | 561 |
| 5 | HEK293T ITGB3OE | ITGB3 was overexpressed | 561 |
| 6 | HEK293T DSG2RNAi | DSG2 expression levels were persistently decreased | 405 |
| 7 | HEK293T ITGB3RNAi | ITGB3 expression levels were persistently decreased | 405 |
| 8 | HEK 293T WT | Human embryonic kidney cell line | N.A |

**Table S3.**

Comparison of gene expression levels in 3D and 2D culture conditions

| **Gene** | **Regulation** | **Fold change** | **Description** |
| --- | --- | --- | --- |
| Sarcoma antigen 1 (SAGE1) | up | 53.43 | Cancer embryonic antigen, involved in tumor occurrence and development |
| MAGE Family Member C2（MAGEC2） | up | 18.58 | Cancer embryonic antigen, involved in tumor occurrence and development |
| Paired Box 1（PAX1） | up | 16.81 | Promote the occurrence and progression of cancer |
| CD274 Molecule（CD274） | up | 5.10 | Drive the progression of cancer |
| （MAGE Family Member H1（MAGEH1） | up | 4.73 | Tumor-specific antigen |
| RAS Guanyl Nucleotide Releasing Factor 1（RASGRF1） | up | 4.77 | Associated with tumorigenesis |
| MAGE Family Member A3（MAGEA3） | up | 4.38 | Cancer embryonic antigen, involved in tumor occurrence and development |
| Protocadherin Gamma Subfamily B5（PCDHGB5） | up | 21.23 | cell-cell interactions |
| Protocadherin Gamma Subfamily A12（PCDHGA12） | up | 4.64 | cell-cell interactions |
| Cadherin 18（CDH18） | up | 3.03 | cell-cell interactions |
| Cadherin 11（CDH11） | down | -18.07 | cell-cell interactions |
| Desmoglein 2（DSG2） | No change | -1.42 | cell-cell interactions |
| Integrin Beta 7（ITGB7） | up | 3.79 | cell-extracellular matrix adhesion |
| Integrin Alpha X（ITGAX） | down | -7.25 | cell-extracellular matrix adhesion |
| Integrin Beta 3（ITGB3） | No change | -1.51 | cell-extracellular matrix adhesion |

**Table S4.**

Comparison of gene expression levels in hypoxia and normoxia in 3D culture conditions

| **Gene** | **Regulation** | **Fold change** | **Description** |
| --- | --- | --- | --- |
| Microtubule-Associated Protein 6 (MAP6) | up | 2.75 | Regulates microtubules polymerization |
| Dihydropyrimidinase-Like 4 (DPYSL4) | up | 2.90 | Promotes microtubule polymerization |
| EPH Receptor B3 (EPHB3) | up | 2.29 | Regulates the polymerization of the actin cytoskeleton |
| Neural Precursor Cell Expressed, Developmentally Down-Regulated 9 (NEDD9) | up | 2.61 | Regulates the dynamic rearrangement of the cytoskeleton |
| Myocardial Z-Line Associated Protein（MYZAP） | up | 2.11 | Affects the dynamic rearrangement of actin filaments |
| Fascin Actin-Bundling Protein 1（FSCN1） | up | 2.03 | Overregulation of actin filament bundling |
| Retinoic Acid-Inducible Gene I（RIGI） | up | 1.84 | Activation of RhoA leads to the polymerization of actin filaments |
| Collagen Type XVI Alpha 1 Chain（COL16A1） | up | 2.75 | Involved in the assembly and stabilization of the extracellular matrix, providing physical support to the cells |
| Matrix Metallopeptidase 23B（MMP23B） | up | 2.67 | Degrading various components of the extracellular matrix, a process crucial for tissue remodeling, cell migration |
| Hyaluronan Synthase 2（HAS2） | up | 2.31 | Synthesize hyaluronic acid |
| Collagen Type XII Alpha 1 Chain（COL12A1） | up | 2.17 | Involved in the assembly and stabilization of collagen fibers in the extracellular matrix |
| Glucokinase（GCK） | up | 3.34 | Catalyze the reaction between glucose and ATP to produce glucose-6-phosphate (G6P) and ADP |
| Aldolase C（ALDOC） | up | 3.18 | Catalyze a key reaction in the glycolytic pathway, where fructose-1,6-bisphosphate (F1,6BP) is cleaved to produce glyceraldehyde-3-phosphate (G3P) and dihydroxyacetone phosphate (DHAP). |
| Phosphoglycerate Kinase 1（PGK1） | up | 2.41 | Catalyze the reaction between glyceraldehyde-3-phosphate (G3P) and phosphoenolpyruvate (PEP) to produce 1,3-bisphosphoglycerate (1,3-BPG) and ATP |
| Protocadherin 15（PCDH15） | down | -4.00 | Ensures tight junctions between cells |
| Protocadherin 20（PCDH20） | down | -3.25 | Participates in cell-cell adhesion |
| Mucin 19（MUC19） | down | -2.66 | Participates in cell surface adhesion |
| Claudin 23（CLDN23） | down | -2.50 | Encodes a tight junction protein |
| Tight Junction Protein 2（TJP2） | down | -2.26 | Encodes a tight junction protein |
| CD34 Molecule（CD34） | down | -2.18 | Encodes a cell surface glycoprotein |
| Protocadherin Gamma Subfamily A12（PCDHGA12） | down | -1.80 | Encodes a cell adhesion protein |
| Cadherin 4（CDH4） | down | -1.75 | Promotes cell-cell adhesion and interactions. |
| Integrin Beta 1（ITGB1） | No change | 1.04 | cell-extracellular matrix adhesion |
| Integrin Beta 3（ITGB3） | No change | -1.27 | cell-extracellular matrix adhesion |

**Table S5.**

Summary of tumorigenesis and kinetics under different culture conditions

|  | **Tumorigenesis Rate [%]** | **Fusion Frequency [%]** | **Fusion Mode** | **Fusion Time [h]** | **Speed**  **[μm/min]** | **Circularity** |
| --- | --- | --- | --- | --- | --- | --- |
| HEK293T 21%O2 | 147 | 23.9 | Growth | 21.1 | 0.04 | 0.93 |
| HEK293T 2%O2 | 321 | 47.5 | Directional | 9.4 | 0.06 | 0.76 |
| HEK293T 21%O2 Rho activate Ⅱ | 424 | 49.5 | Directional | 6.4 | 0.06 | 0.62 |
| HEK293T 2%O2 Y-27632 | 123 | 13.4 | Growth | 28.2 | 0.02 | 0.95 |
| HEK293T 21%O2  DGS2RNAi | 207 | 38.2 | Directional | 12.1 | 0.05 | 0.88 |
| HEK293T 21%O2  ITGB3RNAi | 155 | 23.5 | Directional | 21.2 | 0.03 | 0.94 |
| HEK293T 2%O2  DGS2OE | 188 | 23.0 | Growth | 14.9 | 0.05 | 0.88 |
| HEK293T 2%O2  ITGB3OE | 329 | 46.4 | Directional | 8.87 | 0.07 | 0.80 |

**Legends for Supplementary Movies**

**Movie S1.** A typical time-lapse confocal movie in a large field of view that compares the tumorigenesis process under normoxia (**left**) and hypoxia (**right**) conditions in 3D culture. Imaging was performed using a 10x objective lens, with F-actin labeled with green fluorescence and H2B with red fluorescence. Images were acquired every hour over a total duration of 42 hours. The laser wavelengths used for imaging were 488 nm and 561 nm, with a laser intensity of 20 % and an exposure time of 200 ms. For each z-scanning, the image was captured every 8 μm over the range of 168 μm, resulting in a total of 21 images. The move is about 18514 times the real time. The scale bar is 100 μm.

**Movie S2.** A typical time-lapse confocal movie in a zoom-in field of view that compares the spheroid dynamics under normoxia (**left**) and hypoxia (**right**) conditions in 3D culture. Imaging was performed using a 60x oil objective lens, with F-actin labeled with green fluorescence and H2B with red fluorescence. Imaging was performed at 1-hour intervals over a total duration of 42 hours. The laser wavelengths used for imaging were 488 nm and 561 nm, with a laser intensity of 100 mW and an exposure time of 200 ms. For each z-scanning, the image was captured every 5 μm over the range of 65 μm, resulting in a total of 13 images. The movie is 21600 times the real-time. The scale bar is 20 μm.

**Movie S3.** A typical time-lapse confocal movie that demonstrates the directional fusion of the tumor spheroids. Both top-down (left) and side view (right) are provided. A typical time-lapse confocal movie that demonstrates the active fusion of the tumor spheroids. Both top-down (**left**) and side view (**right**) are provided. The imaging parameters are the same as those of **Movie S2**. The movie is 21600 times the real-time. The scale bar is 20 μm.

**Movie S4.** A typical time-lapse confocal movie that demonstrates the growth fusion of the tumor spheroids. Both top-down (**left**) and side view (**right**) are provided. The imaging parameters are the same as those of **Movie S2**. The movie is 18800 times the real time. The scale bar is 20 μm.

**Movie S5.** A typical time-lapse confocal movie that compares the response of a tumor spheroid after laser ablation under normoxia (**left**) and hypoxia (**right**) conditions. The intensity of the 405 nm ablation laser was set to 50 mW, with each tumor spheroid subjected to 4-5 laser exposures. Throughout the imaging process, a 488 nm laser was utilized, capturing a single-layer image of the maximum section of the spheroid at a frame rate of 200 fps. The movie is 51 times the real-time. The scale bar is 20 μm.

**Movie S6.** A typical time-lapse confocal movie that demonstrates the behavior of the tumor spheroids after pharmaceutical interference of Rho Activator II (**left**) and Y-27632 (**right**). The PHAi for Rho Activator II was conducted under normoxia conditions and that for Y-27632 was done under hypoxia conditions. The imaging parameters are the same as those of **Movie S2**. The movie is 18900 times the real-time. The scale bar is 20 μm.

**Movie S7.** A typical movie that highlights the F-actin channel and tiny cells locating between the tumor spheroids under 3D hypoxia conditions. Green fluorescence was used to label F-actin with phalloidin, and blue fluorescence to label the cell nucleus with Hoechst 33258, as detailed in the experimental methods. Imaging was performed using 405 nm and 488 nm lasers, with a power of 100mW, laser intensity at 20%, and an exposure time of 200 ms. The Z-axis motor of the microscope was employed to move the sample stage along the Z-axis, acquiring continuous optical sections at different Z-axis positions, with a 2 μm interval between adjacent layers. The acquired two-dimensional images were then aligned using Cellvis software and stacked in Z-axis order. These images were processed and analyzed via algorithms to reconstruct the three-dimensional structure of the sample. The scale bar is 40 μm.

**Movie S8.** A typical movie for the dynamics of the outstretched F-actin that guides directional fusion. Imaging was performed using a 40x objective lens, with F-actin labeled by red fluorescence. Imaging was conducted hourly over a period of 42 hours. To capture F-actin dynamics more effectively, the samples were first cultured in a hypoxic incubator for 24 hours, followed by hypoxic imaging in a live-cell workstation. A 640 nm laser was used with a power of 100 mW and an exposure time of 200 ms. For each Z-scan, images were acquired at 3 μm intervals within a 60 μm range, yielding a total of 21 images. The movie is 18000 times the real time. The scale bar is 20 μm.

**Movie S9.** A typical time-lapse confocal movie that demonstrates the tiny cells under hypoxia conditions. The imaging parameters are the same as those of **Movie S2**. The movie is 18900 times the real time. The scale bar is 20 μm.

**Movie S10.** A typical time-lapse confocal movie that compares cell cycle dynamics in the tumor spheroid under normoxia (**left**) and hypoxia (**right**) conditions in 3D culture. Cells in the G1 phase are labeled with red fluorescence, and cells in the S/G2/M phases are labeled with green fluorescence. Other imaging parameters are the same with those of **Movie S2**. The movie is 19800 times the real time. The scale bar is 20 μm.
